# Biomechanical Phenotyping Reveals Unique Mechanobiological Signatures of Early-Onset Colorectal Cancer

**DOI:** 10.1101/2025.07.15.664829

**Authors:** Nicole C. Huning, Munir H. Buhaya, Victor V. Nguyen, Afeefah Khazi-Syed, Haider A. Ali, Adil Khan, Angela Fan, Robert C. Fisher, Zhikai Chi, Indu Raman, Guangchun Chen, Chengsong Zhu, Mengxi Yu, Andrew R. Jamieson, Sara Roccabianca, Victor D. Varner, Cheryl M. Lewis, Emina H. Huang, Jacopo Ferruzzi

## Abstract

While both incidence and mortality of sporadic average-onset colorectal cancer (AO CRC, above 50 years of age) are in constant decline, sporadic early-onset colorectal cancer (EO CRC, under 50 years of age) is rising rapidly. Yet, the causes behind this rise remain poorly understood. Epidemiological studies indicate that lifestyle and environmental exposures may result in chronic inflammation, which is known to trigger tissue fibrosis. This study tests the hypothesis that fibrotic remodeling and biomechanical stiffening of colorectal tissues represent measurable hallmarks and potential drivers of EO CRC. Using primary human tissues, this work shows that EO CRC is associated with changes in collagen microstructure, increased stiffness and elevated viscosity of primary tumors. Spatial transcriptional profiling and immunostaining reveal pro-fibrotic signatures in stromal cells, alongside enhanced Yes-associated protein (YAP) mechanotransduction and proliferation in epithelial cells of EO CRC tissues. Mechanistically, increasing matrix stiffness in vitro promotes proliferation of epithelial cells in 2D and 3D colorectal cancer models. Together, these findings establish EO CRC as a disease marked by early and widespread biomechanical remodeling, suggesting that a fibrotic and stiffened tissue microenvironment may orchestrate EO CRC tumor initiation.

## INTRODUCTION

Colorectal cancer (CRC) ranks as the third most diagnosed cancer worldwide and the second leading cause of cancer-related death in the United States.^[1–3]^ Over the past decades, increased rates of routine colonoscopy screening have led to a steady decline in the incidence of average-onset colorectal cancer (AO CRC), which affects patients aged 50 years or older.^[4]^ In contrast, early-onset colorectal cancer (EO CRC), which affects patients under 50 years of age,^[5]^ has been rising by 0.5% to 2.4% annually for 3 decades, comprising 12% of all CRCs diagnosed in the United States since 2020.^[6]^ Based on current estimates, the most rapid increase in EO CRC will be experienced by individuals between 20 and 34 years old.^[7]^ Younger patients are also more likely to present with advanced-stage disease.^[8]^ If the current trend persists, the incidence of EO CRC is expected to increase by 124% by the year 2030.^[9]^ In response, independent panels of experts now recommend starting colonoscopy screening at age 45 instead of 50.^[10,11]^ However, these updates in medical guidelines were made in the absence of a clear mechanistic understanding of EO CRC pathogenesis or progression. Thereby, critical gaps in knowledge persist and hinder the development of tailored screening tools and treatments strategies for younger patients.

EO CRC has no unique mutational drivers differentiating it from AO CRC.^[12–14]^ The rapid rise in incidence over just 3 decades suggests that genetic changes alone are insufficient to explain the emergence of this disease.^[15]^ EO CRC often presents at a late stage at diagnosis with aggressive pathological features including mucinous, signet ring, and poorly differentiated tumors.^[12]^ Further, the majority of EO CRC cases are microsatellite stable (MSS) and sporadic in nature (i.e., cannot be attributed to genetic germline mutations).^[16]^ EO CRC is often located within the distal colon or rectum, a feature referred to as left-sidedness that is more pronounced than in AO CRC, which occurs more frequently in the proximal colon.^[12,17]^ The *birth-cohort effect* hypothesis^[18]^ suggests that early life exposures are critical to understanding the etiology of EO CRC, with adults born in the 1990s having a higher risk of developing both colon and rectal cancers than adults born in the 1950s.^[19]^ Emerging evidence supports the consideration of exposomal elements (that is, the totality of nongenetic environmental exposures) in the genesis of EO CRC.^[9]^ This literature highlights how external exposures (e.g., diet and physical activity) and internal exposures (e.g., the gut microbiota) may drive oncogenesis via chronic inflammation.^[20–22]^ In the context of EO CRC, it is conceivable that both the birth-cohort effect and lifestyle exposures are permissive of inflammation.^[23]^ In turn, chronic inflammation may also promote microenvironmental alterations, further contributing to disease development.^[9,24]^

The tumor microenvironment (TME) undergoes profound alterations during colorectal carcinogenesis, with extracellular matrix (ECM) remodeling emerging as a central feature that influences cancer progression. Activation of tissue-resident fibroblasts drives the deposition and reorganization of a collagen-rich ECM, fundamentally altering the biomechanical properties of tissues.^[25–28]^ These ECM modifications create a mechanically distinct microenvironment characterized by increased tissue stiffness and altered matrix architecture. In breast^[29]^ and pancreatic^[30]^ cancers, such microenvironmental remodeling has been demonstrated to drive malignant phenotypes through mechanotransduction and altered cell-matrix interactions. Although the role of ECM remodeling in CRC progression remains incompletely understood, with conflicting reports of both pathogenic^[31,32]^ and protective^[33,34]^ effects, prior studies documented altered tissue stiffness in AO CRC^[35–38]^. Critically, biomechanical characterization of the TME in EO CRC remains unexplored, representing a significant gap in the understanding of this disease’s distinct pathophysiology.

To fill this gap in knowledge, we quantified the biomechanical properties of primary tumors and matched normal tissues from AO and EO CRC patients. We hypothesized that tissues from EO CRC patients are more fibrotic and thus become biomechanically stiffer. Our results consistently reveal that collagen remodeling and mechanical stiffening are greater in EO CRC tissues than AO CRC tissues, with structural and mechanical abnormalities occurring also in histologically normal tissues from EO CRC patients. Furthermore, we show that these changes in connective tissue structure and mechanics trigger altered mechanotransduction and increased proliferation in colonic epithelial cells. Our findings suggest that biomechanical stiffening may serve as both a driver and diagnostic predictor of EO CRC. Overall, we identify increased tissue stiffness as a quantifiable hallmark of EO CRC, which could serve as a potential biomarker to stratify cancer risk in young populations.

## RESULTS

### EO CRC demonstrates distinct left-sided and distal anatomical localization

To investigate the structural and biomechanical properties of EO CRC, we collected colonic tissue specimens harvested from patients undergoing surgical resection. Our study cohort included 19 patients with AO CRC (68 ± 11 years of age) and 14 patients with EO CRC (44 ± 2 years of age) with demographic information and clinical characteristics listed, respectively, in **Table S1** and **S2**. Both groups included men and women of mixed ethnicities and races (Table S1 and S2) with a body mass index (BMI) that was on average elevated (32.6 ± 8.2 for AO CRC and 27.2 ± 3.5 for EO CRC), without statistically significant differences between the two cohorts. Consistent with patients who present for curative resection, tumor stage distribution was similar between groups, with most patients having stage II or III CRC. The most striking difference between the groups was tumor location (**Figure S1**). The most frequent location for AO CRC was the cecum (36.8%) and the least frequent location was the rectum (0%). Conversely, the most frequent location for EO CRC was the sigmoid colon (42.9%) and the least frequent location was the right colon along with the hepatic and splenic flexures (0%). Despite potential selection bias in our small patient cohort, our data are similar to data listed in large epidemiological studies^[12]^ showing that EO CRC more frequently localizes to the left colon and rectum than in AO CRC (**Figure 1**A and S1).

**Figure 1.**
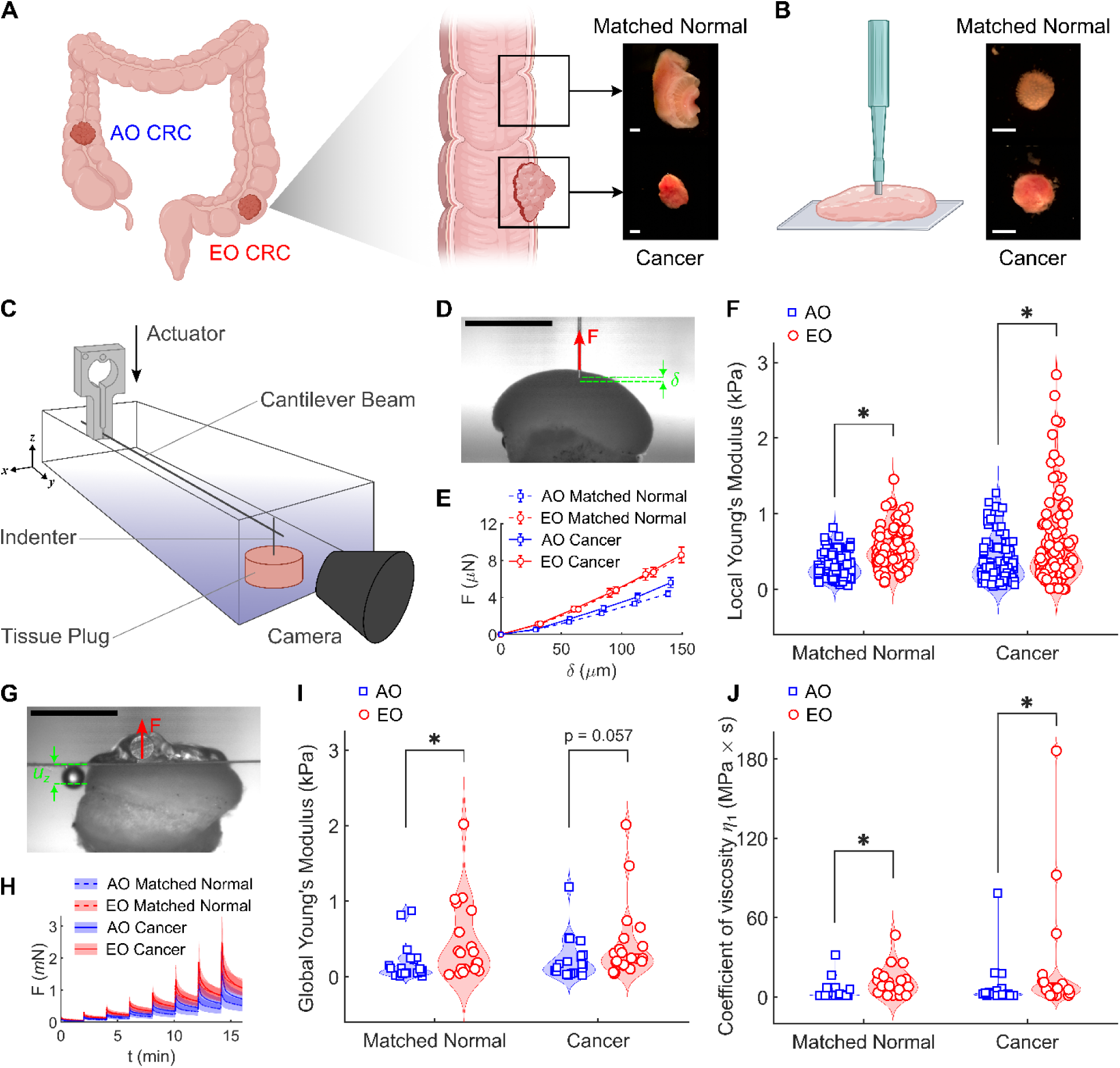
Primary tissues from sporadic early-onset colorectal cancer (EO CRC) patients exhibit increased material stiffness and viscosity across length scales. (**A**) Cancer and matched normal tissues were harvested from average-onset (AO) and early-onset (EO) colorectal cancer (CRC) patients undergoing surgical resection and (**B**) cylindrical specimens for mechanical characterization were prepared using a 3 mm biopsy punch. Scale bars, 2 mm. (**C**) Schematic of the mechanical testing setup, in which a cantilever beam of known stiffness was equipped with an indenter for local (mesoscale) testing or with a flat platen for global (macroscale) testing. (**D**) Representative camera caption showing the imposed indentation (δ) and the resulting force (F) measured locally from a colorectal tissue specimen. Scale bar, 2 mm. (**E**) Average force-indentation curves and (**F**) associated stiffness values (quantified as local Young’s moduli) measured for the four groups (n = 94 for AO Matched Normal, n = 89 for AO Cancer, n = 80 for EO Matched Normal, n = 102 for EO Cancer). (**G**) Representative camera caption showing the imposed displacement (u_z_) and the resulting force (F) measured globally from a colorectal tissue specimen. Scale bar, 2 mm. (**H**) Average force evolution over time upon 8 consecutive stress-relaxation steps and (**I**) associated stiffness values (quantified as global Young’s moduli) and (**J**) fast coefficient of viscosity measured for the four groups (n = 21 for AO Matched Normal, n = 21 for AO Cancer, n = 19 for EO Matched Normal, n = 21 for EO Cancer). Mechanical data are presented as mean ± SEM while associated quantifications of stiffness and viscosity are presented as violin plots with data points indicating individual measurements. Statistical significance was assessed using a Scheirer-Ray-Hare test with Dunn’s post-hoc test and Bonferroni correction. * indicates significant differences (*P* < 0.05) between cancer and matched normal tissues within onset groups (AO, EO) and between onset groups within tissue types (matched normal, cancer). Schematics in (**A**) and (**B**) created using Biorender.com.

### EO CRC tissues exhibit elevated material stiffness and viscosity

For each patient, we collected primary tumor tissues as well as matched normal tissues (retrieved at least 5 cm from the tumor) and extracted cylindrical tissue plugs (3 mm in diameter) using a biopsy punch for ex vivo biomechanical testing (Figure 1B). We quantified the mechanical properties of such tissue plugs using a computer-controlled system, which estimates tissue-generated forces by measuring the deflection of an elastic cantilever beam (Figure 1C). We assessed tissue stiffness at two distinct length scales via mesoscale (tens of micrometers) and macroscale (millimeter range) measurements, requiring distinct modeling approaches. For mesoscale properties, we used a cylindrical indenter having a radius of 38.1 µm (Figure 1D) to conduct local force-indentation tests (Figure 1E) and to estimate the local Young’s modulus under the assumption of linear elastic behavior (Figure 1F). We modified previous protocols^[39,40]^ and used the cylindrical indenter to impose quasi-static small deformations (up to 5% of the total height for each tissue plug) and measured the resulting elastic forces generated on the mucosal surface (**Figure S2**A-H). At this length scale, both cancer and matched normal tissues from EO CRC were materially stiffer than their AO counterparts (Figure 1F). Within each patient cohort, cancer and matched normal tissues showed no statistically significant differences in stiffness, although cancer tissues revealed a broader distribution of tissue stiffness than matched normal tissues (Figure S2I-J), consistent with other reports.^[36,41]^

To validate these mesoscale findings, we assessed macroscale tissue properties by pairing the cantilever beam with a 6 mm × 6 mm square platen (Figure 1G). Following our previous work^[42,43]^, we performed stress-relaxation tests in unconfined compression (Figure 1H) and used the steady-state stress values to estimate the global Young’s modulus (Figure 1I). We used this method to impose large deformations (up to 20% of the total height for each tissue plug) and measured the resulting viscoelastic forces generated by the tissues while recording the radial tissue expansion due to the Poisson effect (**Figure S3**A-G). These combined measurements allowed us to estimate the true (Cauchy) stress developed within each tissue upon compression (Figure S3H). By combining the neo-Hookean shear moduli and the Poisson’s ratios measured from each specimen, we calculated Young’s moduli that were comparable to those obtained using local indentation (Figure 1I), thereby validating our local measurements.

In addition to greater stiffness, EO tissues also exhibited greater viscosity (Figure 1J and S3I-K). Using a nonlinear Maxwell-Wiechert model appropriately modified from previous reports^[44,45]^, we modeled the stress-relaxation data (**Figure S4**A-B) to capture the nonlinear and viscoelastic properties of colorectal tissues. We found that both matched normal and cancer tissues from EO CRC were more viscous than AO CRC tissues (Figure 1J). Indeed, human colorectal tissues displayed a mild but noticeable nonlinearity, which could be modeled by incorporating a nonlinear spring in parallel to two Maxwell elements (Figure S4C-D). Quantification of individual model parameters revealed no significant differences between AO and EO CRC in the nonlinear spring parameters ( *c_a_* , *c_b_* ) (Figure S4E-F). Instead, we found that the coefficient of viscosity for the fast Maxwell element (*η*_1_ ) was higher in both matched normal and cancerous EO tissues (Figure 1J) while the slow Maxwell element (*η*_2_ ) and the Maxell spring constant ( *c_m_* ) were consistently higher only in EO matched normal tissues (Figure S4G-I). Taken together, our multiscale biomechanical data interpreted using both linear elastic and nonlinear viscoelastic modeling reveal that human tissues from EO CRC were materially stiffer and more viscous than human tissues from AO CRC.

### EO CRC displays consistent patterns of fibrotic collagen remodeling

To investigate the structural basis of the observed changes in tissue mechanics, we examined histological cross-sections from formalin-fixed and paraffin-embedded (FFPE) tissues and quantified the changes in collagen fiber content, distribution, and organization. We used Masson’s Trichrome (MTC) staining to differentiate collagen (blue) from cellular elements (red) and a performed a colorimetric image-based quantification to determine collagen and cytoplasm area fractions (**Figure 2**A and **S5**A-E). Matched normal tissues preserved a well-organized structure comprising mucosa, submucosa, muscularis propria, and serosa. Accordingly, the collagen content was higher in the normal submucosa with respect to the normal mucosa (Figure S5F-G). Conversely, cancer tissues lost such layered organization and displayed interspersed stromal and epithelial compartments. When comparing collagen content between the normal submucosa and the cancer stroma in AO and EO CRC tissues, we found that both AO and EO cancer tissues had lower collagen content with respect to their normal submucosa, which likely reflects an increased tumor cellularity (Figure 2B-C). However, EO cancers retained more collagen than AO cancers, suggesting enhanced matrix deposition and remodeling in EO CRC (Figure 2C). To assess collagen fiber structure, we performed Picrosirius Red (PSR) staining in combination with polarized light (POL) imaging (Figure 2D). This approach allows us to distinguish collagen fiber thickness, packing, and maturity based on the color of birefringent fibers: red denotes thicker, tightly packed, and mature fibers; orange-yellow represent intermediate thickness and packing; green indicates thinner, loosely packed, and immature fibers^[46–49]^ (**Figure S6**). EO tissues, both cancer and matched normal, contained higher proportions of thick (red) collagen fibers than AO tissues (Figure 2E-F), which mirrors the increased tissue stiffness observed biomechanically (Figs. 1F and 1I). Although PSR staining is widely used to assess fibrosis, it lacks subtype specificity, and POL imaging is dependent on the orientation of the collagen fiber bundles^[50]^. To address the latter concern, we quantified the architectural organization of stromal collagen using multiphoton label-free Second Harmonic Generation (SHG) imaging^[51–54]^ from hematoxylin and eosin (H&E) stained slides (Figure 2G). We analyzed the SHG images using Ridge Detection^[55]^ to quantify mean fiber length, width, orientation, and SHG signal intensity across all pixels (**Figure S7**A-C). Compared to AO normal tissues, EO matched normal tissues exhibited shorter collagen fibers with a higher SHG signal intensity, reflecting a greater density of collagen fibers (Figure S7D). SHG signal intensity decreased in tumors relative to matched normal tissues in both groups paralleling a reduction in collagen fiber width (Figure S7D-E). Notably, we found that EO CRC collagen fibers were longer and more aligned with respect to fibers in EO normal or AO cancer samples (Figure 2H-I and S7F-G), a pattern reminiscent of the aligned collagen found in invasive breast cancers^[56]^. This finding is particularly relevant as it may link collagen alignment with the highly invasive and metastatic behavior reported in EO CRC.^[9]^

**Figure 2.**
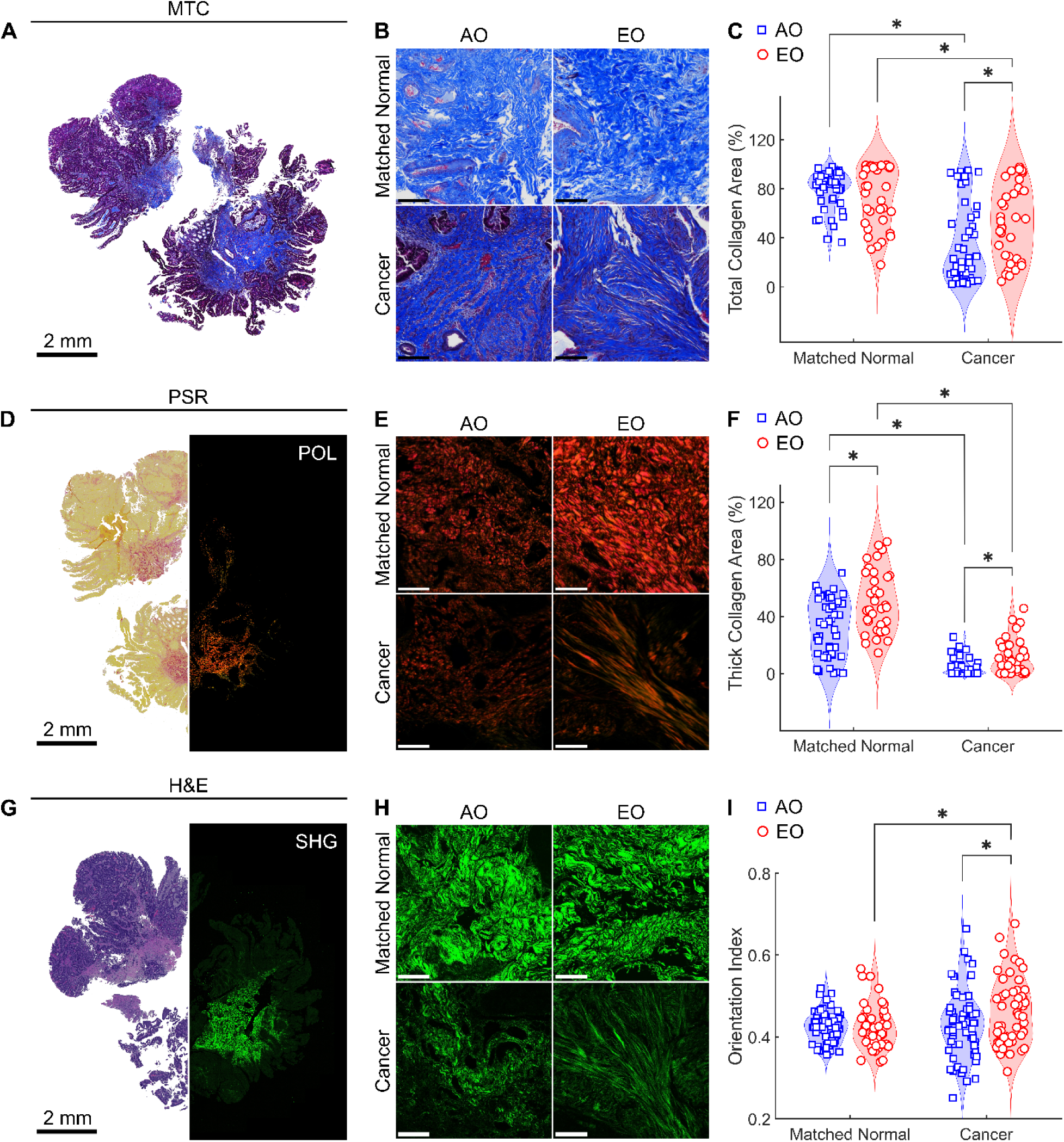
Tissues from sporadic EO CRC patients exhibit collagen remodeling consistent with fibrosis. (**A**) Masson’s Trichrome (MTC) staining of a patient tissue sample and (**B**) representative zoomed-in images (1 mm × 1 mm) for the four groups. (**C**) Quantitative assessment of the area fraction occupied by collagen in the normal submucosa and in the tumor stroma (n = 45 for AO Matched Normal, n = 45 for AO Cancer, n = 35 for EO Matched Normal, n = 35 for EO Cancer). (**D**) Picrosirius Red (PSR) staining of a patient tissue sample imaged under brightfield illumination (left) and under polarized light (POL, right) and (**E**) representative zoomed-in images (1 mm × 1 mm) for the four groups. (**F**) Quantitative assessment of the area fraction occupied by thick collagen fibers (red) in the normal submucosa and in the cancer stroma (n = 45 for AO Matched Normal, n = 45 for AO Cancer, n = 35 for EO Matched Normal, n = 35 for EO Cancer). (G) Hematoxylin and eosin (H&E) staining of a patient tissue sample imaged under brightfield illumination (left) and using multiphoton Second Harmonic Generation (SHG, right) imaging, with (H) representative zoomed-in images (1 mm × 1 mm) for the four groups. (**I**) Quantitative assessment of collagen fiber alignment via a dimensionless Orientation Index (OI) in the normal submucosa and in the cancer stroma (n = 65 for AO Matched Normal, n = 64 for AO Cancer, n = 47 for EO Matched Normal, n = 55 for EO Cancer). Unless otherwise specified, scale bars indicate 200 µm. Data are presented as violin plots with data points indicating individual measurements. Statistical significance was assessed using a Scheirer-Ray-Hare test with Dunn’s post-hoc test and Bonferroni correction. * indicates significant differences (*P* < 0.05) between cancer and matched normal tissues within onset groups (AO, EO) and between onset groups within tissue types (matched normal, cancer).

These data demonstrate that EO CRC exhibits consistent fibrotic features with denser and more aligned collagen fibers across both tumor and adjacent normal tissues. These structural alterations support the increased tissue stiffness observed in EO CRC and point to fibrosis as a defining feature of the EO CRC TME.

### EO CRC presents unique cell morphologies and epithelial-mesenchymal phenotypes

To explore the cellular characteristics underlying a dysregulated ECM in EO CRC, we performed high-resolution immunofluorescence imaging on a subset of tissue samples stained with PanCK, an epithelial marker; Vimentin, a mesenchymal marker; and CD45, an immune marker (**Figure 3**A). Although CD45 successfully labeled immune cells, the minimal immune infiltration resulted in low signal intensity, leading to its exclusion from downstream quantifications. Matched normal tissues maintained spatially distinct epithelial and mesenchymal compartments, whereas cancer tissues displayed a disorganized pattern of epithelial cell clusters surrounded by mesenchymal cells (Fig 3B). We conducted single-cell segmentation using an established deep-learning algorithm^[57]^, enabling quantification of 2,155,046 cells from 11 patient-derived samples across the four groups (**Figure S8**). For each segmented cell, we extracted features including morphology (cell area and aspect ratio) and marker expression (PanCK and Vimentin intensity) (Figure 3C). Across all samples, EO tissues exhibited consistent morphological distinctions. The cell area was smaller in both normal and tumor tissues from EO CRC than in AO CRC tissues, although cancer cells were larger than matched normal cells within each group (Figure 3D). The cell aspect ratio showed greater variability: EO normal tissues had higher aspect ratios (i.e., were more elongated) while EO cancer had lower aspect ratios (i.e., were more rounded) than their corresponding AO counterparts (Figure 3E). After segmentation, we z-normalized the fluorescence intensity of the PanCK (epithelial) and Vimentin (mesenchymal) markers to facilitate comparisons of cellular identity markers across groups. We found that PanCK expression was lower in EO than in AO in both normal and tumor tissues (Figure 3F). In contrast, Vimentin expression followed the opposite pattern: EO cells had higher Vimentin levels than AO cells, despite cancer tissues generally expressing lower Vimentin than matched normal tissues (Figure 3G).

**Figure 3.**
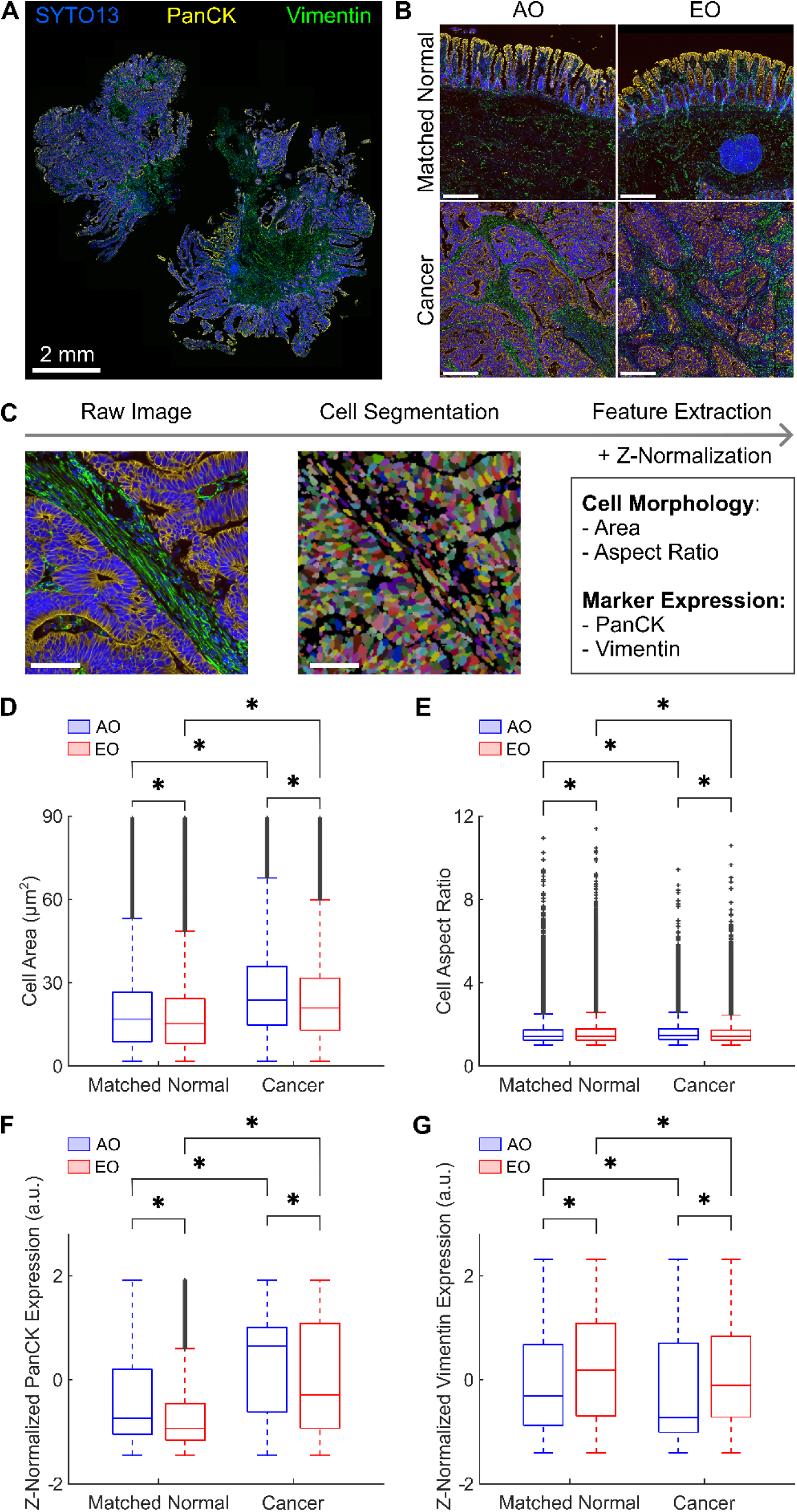
Tissues from EO CRC patients reveal distinct epithelial and mesenchymal cellular phenotypes. (**A**) Immunofluorescence staining of a patient tissue sample and (**B**) representative zoomed-in images (1 mm × 1 mm) for the four groups. Pseudocoloring shows the SYTO13 nuclear marker in blue, the epithelial marker PanCK in yellow, and the stromal marker Vimentin in green. Unless otherwise specified, scale bars indicate 200 µm. (**C**) Image analysis pipeline for immunofluorescence images includes cell segmentation and extraction of single-cell features. Scale bars, 50 µm. Box plots for (**D**) cell area, (**E**) cell aspect ratio, (**F**) z-normalized PanCK expression, and (**G**) z-normalized Vimentin expression show differences in cell morphology and marker expression across the four groups. The expressions of epithelial and stromal markers were z-normalized to facilitate comparisons across a large dataset (n = 176,681 for AO Matched Normal, n = 753,993 for AO Cancer, n = 379,626 for EO Matched Normal, n = 844,746 for EO Cancer). Statistical significance was assessed using a Scheirer-Ray-Hare test with Dunn’s post-hoc test and Bonferroni correction. * indicates significant differences (*P* < 0.05) between cancer and matched normal tissues within onset groups (AO, EO) and between onset groups within tissue types (matched normal, cancer).

To further characterize the observed cellular heterogeneity, we applied unsupervised Louvain clustering^[58]^, which identified four dominant cell clusters based on the average cell morphology and marker expression: epithelial large (PanCK-high, Vimentin-low, cell area-high), epithelial-small (PanCK-high, Vimentin-low, cell area-low), stroma (PanCK-low, Vimentin-high, cell area-high), and immune (PanCK-low, Vimentin-high, cell area-low) (**Figure S9**). This classification yielded spatial distributions that reflected qualitatively the cellular patterns observed in raw immunofluorescence images (Figure S9). Reanalysis of the cellular morphology and marker expression results by cluster confirmed that both EO normal and cancer cells displayed consistently smaller areas across all phenotypes (**Figure S10**). This result potentially reflected the elevated compressive stress experienced by all cell types within the TME^[59–61]^. Stromal cells in EO tissues had a consistently high aspect ratio (Figure S10), mirroring the greater collagen fiber alignment observed in EO cancers (Figure 2I). This finding suggests that fibroblast-like mesenchymal cells actively polarize and align the tumor ECM in EO CRC. Regarding marker expression, both epithelial-large and epithelial-small cells in EO CRC co-expressed higher levels of PanCK and Vimentin (Figure S10), which indicated a mixed epithelial-mesenchymal phenotype. Overall, our single-cell morphological analysis demonstrated that EO CRC exhibited altered cellular morphologies (e.g., smaller areas among all cells, elongated stromal cells) and more epithelial-mesenchymal characteristics than AO CRC.

### Mesenchymal cells in EO CRC present pro-fibrotic transcriptional signatures

Mesenchymal cells, particularly cancer-associated fibroblasts (CAFs), play a critical role in sculpting the TME and in promoting cancer progression.^[62–64]^ Given the increased tissue stiffness, collagen remodeling, and stromal alignment in EO CRC, we next examined spatially resolved RNA expression in FFPE tissue samples from AO and EO CRC patients using the GeoMx® RNA platform.^[65,66]^ We focused first on 13 Vimentin-positive regions of interest (ROIs) from matched normal samples (6 ROIs from AO tissues and 7 ROIs from EO tissues) and 40 Vimentin-positive ROIs from cancer samples (17 ROIs from AO tissues and 23 ROIs from EO tissues) for spatial transcriptomic analysis (**Figure 4**A). Differential expression analysis between EO cancer and AO cancer revealed 32 differentially expressed genes (DEGs, adjusted *P* value < 0.05, |log2 Fold Change| > 1) in stromal cells, with 30 genes upregulated and only 2 genes downregulated in EO cancer (**Figure S11**). Gene ontology (GO) analysis revealed significant enrichment of pathways involved in ECM organization, collagen metabolism, and angiogenesis in EO cancer (Figure 4B). Based on this evidence, we conducted a Gene Set Enrichment Analysis (GSEA) using a curated intestinal fibrosis gene set.^[67]^ We discovered that a transcriptional program promoting intestinal fibrosis is active in EO CRC (Figure 4C) in accordance with our biomechanical and histological findings. We next examined key genes within fibrosis-related pathways across normal and cancer samples in both age groups (Figure 4D-F). First, we examined genes involved in collagen fibrillogenesis and found a significant increase in the expression of genes associated with collagens type I and III (*COL1A1*, *COL1A2*, and *COL3A1*) in cancer tissues with respect to normal for both age groups, while we found no differences between EO and AO cancers (**Figure S12**A). In contrast, we found a significant upregulation in the expression of collagen type VI (*COL6A1* and *COL6A2*) in EO cancer along with the expression of decorin (*DCN*), a small proteoglycan that regulates collagen fiber organization (Figure 4E). Next, we investigated genes involved in ECM remodeling, and found that EO cancers upregulated genes associated with matrix crosslinking, such as lysyl oxidase-like 1 (*LOXL1*) and tissue transglutaminase (*TGM2*), while matrix degradation genes, such as matrix metalloproteinases (*MMP2*, *MMP9*, and *MMP12*) had only modest differences in expression (Figure S12B). Instead, we detected a significant upregulation in the gene encoding tissue inhibitor of metalloproteinases 1 (*TIMP1*) in EO cancer, suggesting a shift in the proteolytic balance toward ECM accumulation (Figure 4D). We also observed elevated expression of genes involved in cell-matrix adhesion and signaling, such as thrombospondin 1 (*THBS1*), the integrin subunit α5 (*ITGA5*), and *CD44* (Figure S12C) in EO cancers. Lastly, EO CRC had upregulated genes involved in inflammation and angiogenesis, such as the C-X-C motif chemokine ligand 1 (*CXCL1*), the monocyte chemoattractant protein-1 (MCP-1 or *CCL2*), and the platelet-derived growth factor receptor beta (*PDGFRβ*), when compared to AO CRC (Figs. 4F and S12D-E). Our trascriptional analysis of Vimentin-positive cells confirmed that stromal cells in EO CRC exhibited a distinct pro-fibrotic transcriptional program, characterized by enhanced collagen deposition, crosslinking, adhesion signaling, and inflammatory remodeling. These features likely contribute to the fibrotic and mechanically altered TME that distinguishes EO CRC from AO CRC.

**Figure 4.**
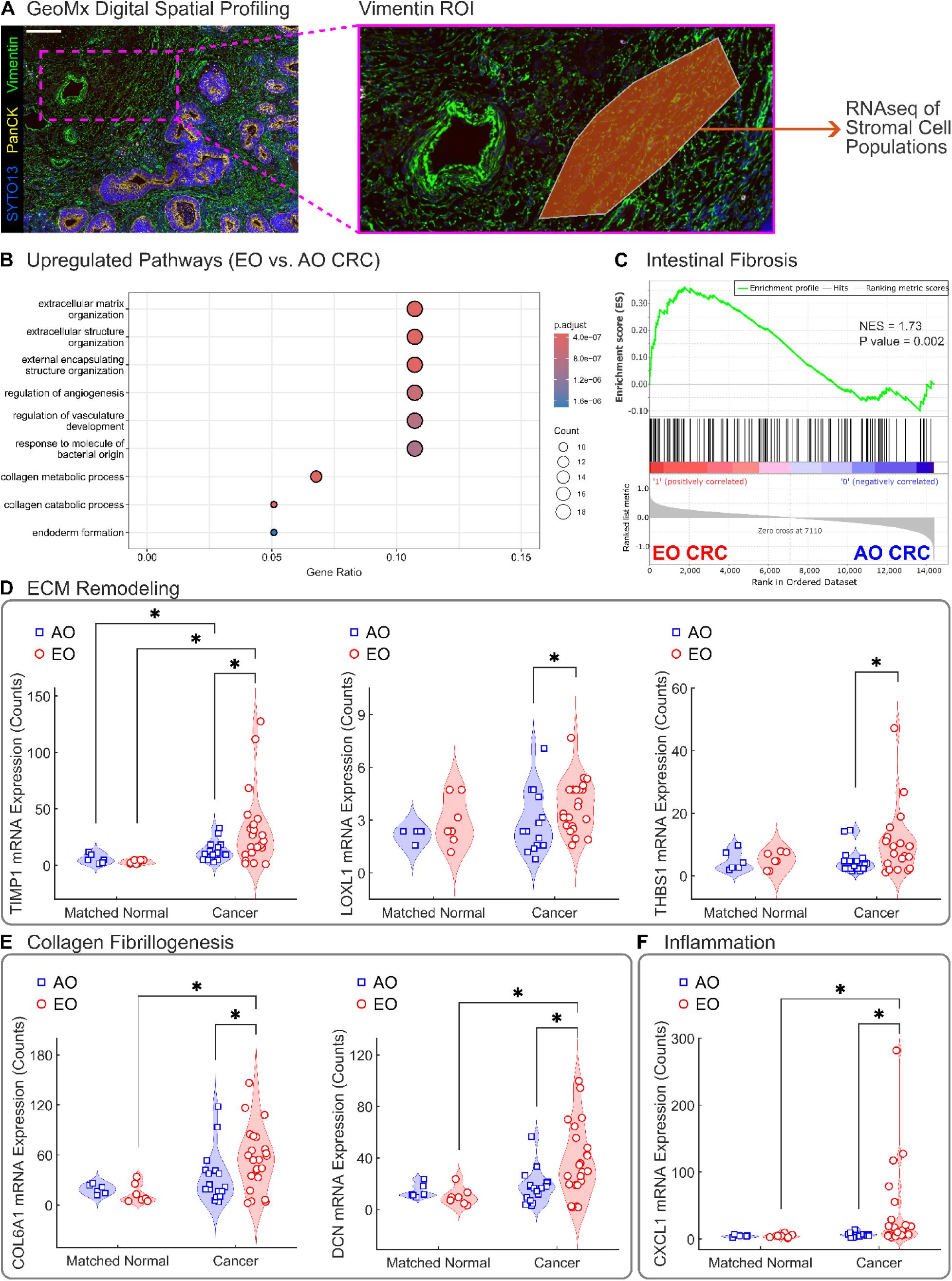
Spatial transcriptomic profiling reveals pro-fibrotic signatures in the EO CRC stroma. (**A**) Immunofluorescence staining of a patient tissue sample and a representative Vimentin-positive region of interest (ROI) used for digital spatial profiling (DSP) with the GeoMx® RNA assay. Pseudocoloring shows the SYTO13 nuclear marker in blue, the epithelial marker PanCK in yellow, and the stromal marker Vimentin in green. Scale bar, 100 μm. (**B**) Gene ontology (GO) analysis showing the top upregulated pathways in EO CRC compared to AO CRC. The x-axis represents the Gene Ratio (number of genes in the pathway divided by the total number of genes analyzed), the bubble size is proportional to the count of genes in each pathway, and the color intensity indicates the adjusted *P* value significance level. ECM and collagen remodeling are highlighted as key biological processes significantly enriched in the stroma of sporadic EO CRC. (**C**) Gene Set Enrichment Analysis (GSEA) confirms upregulation of genes associated with intestinal fibrosis.^[67]^ Normalized enrichment score (NES = 1.73) and *P* value (*P* = 0.002) demonstrate significant upregulation of intestinal fibrosis in EO CRC. Expression levels of selected genes associated with (**D**) ECM remodeling (TIMP1, LOXL1, THBS1), (**E**) collagen fibrillogenesis (COL6A1, DCN), and (**F**) inflammation (CXCL1). Sample sizes indicate the number of ROIs analyzed for each group (n = 6 for AO Matched Normal, n = 7 for AO Cancer, n = 17 for EO Matched Normal, n = 23 for EO Cancer). Data are presented as violin plots with data points indicating individual ROIs. Statistical significance was assessed using a Scheirer-Ray-Hare test with Dunn’s post-hoc test and Bonferroni correction. * indicates significant differences (*P* < 0.05) between cancer and matched normal tissues within onset groups (AO, EO) and between onset groups within tissue types (matched normal, cancer).

### Epithelial cells in EO CRC exhibit enhanced proliferation and altered mechanosensing

The second step in our digital spatial profiling focused on epithelial cells by identifying PanCK-positive ROIs from cancer samples (9 ROIs from AO tissues and 10 ROIs from EO tissues) for spatial transcriptomic analysis (**Figure 5**A). Differential expression analysis identified 769 DEGs in epithelial cells in EO relative to AO cancer, with 421 genes upregulated and 348 genes downregulated in EO cancer (adjusted *P* value < 0.05, |log2 Fold Change| > 1, Figure 5B). Upregulation of genes encoding olfactomedin 4 (*OLFM4*) and mutiple histone proteins and variants (*H1.5*, *H2BC9*, *H2BC10*, *H3C3*, *H3C7*, *H3C8*) indicated enhanced cell proliferation in EO CRC (Figure 5B). To contextualize these findings, we assessed expression patterns associated with the consensus molecular subtype (CMS) classification of CRC.^[68,69]^ By focusing on the two intrinsic subtypes iCMS2 and iCMS3^[69]^ in our patient cohort, unsupervised hierarchical clustering revealed the expected dichotomy between these subtypes, with iCMS2 samples showing elevated expression of WNT/β-catenin and MYC signaling, while iCMS3 samples had upregulated MAPK pathway genes and inflammatory response signatures (**Figure S13**). Yet, the EO and AO cancer samples did not segregate into these two distinct subtypes (Figure S13).

**Figure 5.**
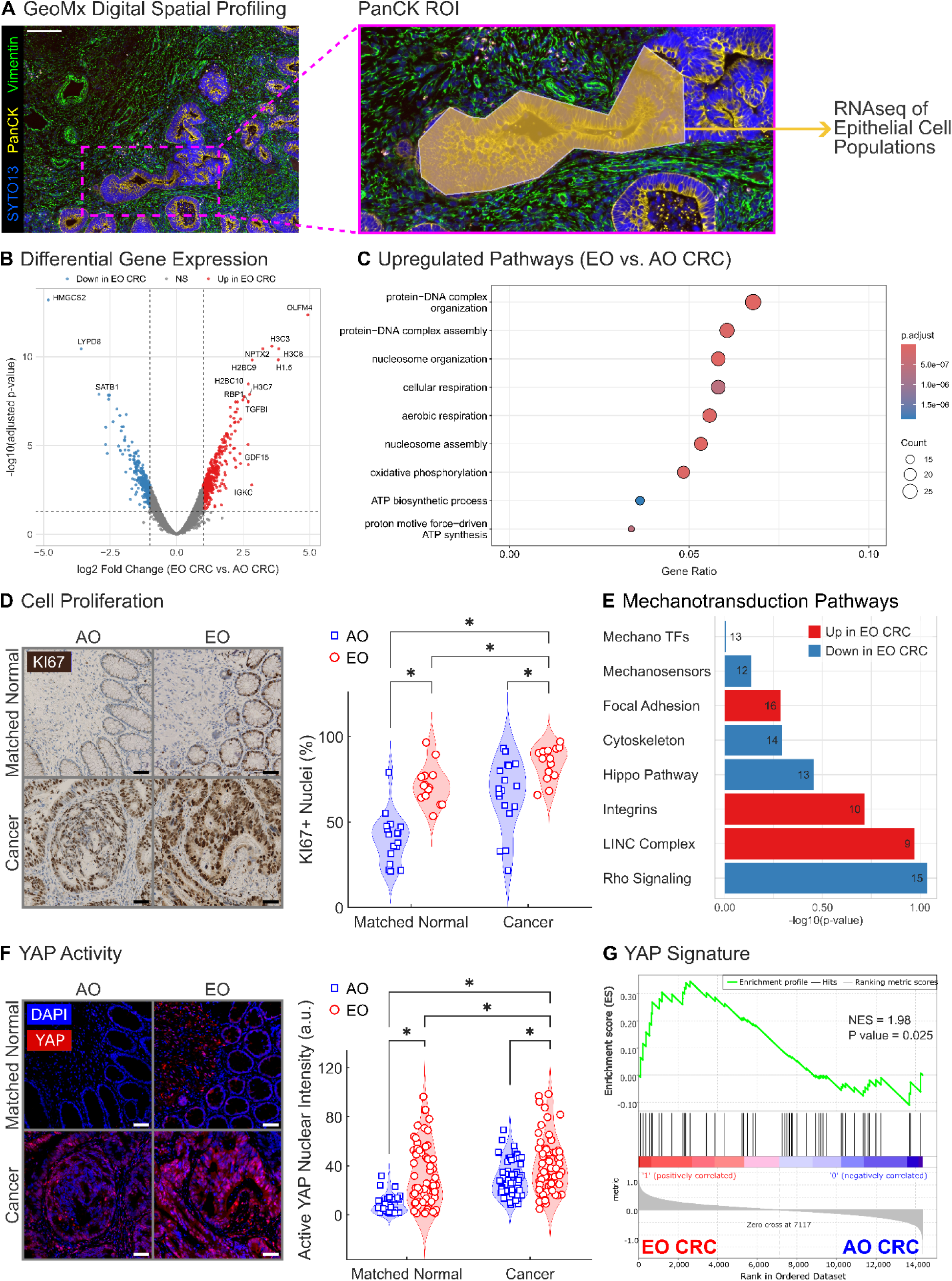
EO CRC epithelial cells exhibit enhanced proliferation and altered mechano-sensing. (**A**) Immunofluorescence staining of a tissue sample and a representative PanCK-positive region of interest (ROI) used for digital spatial profiling with the GeoMx® RNA assay. Pseudocoloring shows the SYTO13 nuclear marker in blue, the epithelial marker PanCK in yellow, and the stromal marker Vimentin in green. Scale bar, 100 μm. (**B**) Volcano plot highlights differential gene expression in PanCK-positive epithelial cells between EO and AO CRC. The x-axis represents log2 fold change (EO CRC vs AO CRC), and the y-axis shows the -log10 of adjusted *P* values. Red dots indicate genes significantly upregulated in EO CRC (log2 fold change > 1, adjusted *P* value < 0.05), blue dots indicate genes significantly downregulated in EO CRC (log2 fold change < -1, adjusted *P* value < 0.05), and grey dots represent non-significantly differentially expressed genes. (**C**) Gene ontology (GO) analysis showing the top upregulated pathways in EO CRC compared to AO CRC. (**D**) Ki67 immunohistochemistry staining of tissue samples confirmed increased cell proliferation in epithelial cells from EO CRC. (**E**) Mechanotransduction pathway enrichment analysis in epithelial cells comparing EO vs. AO CRC. Bar plot shows the significance (-log10 *P* value) of eight distinct mechanotransduction-related gene sets. Bars are colored according to pathway direction (red = upregulated in EO CRC, blue = downregulated in EO CRC), and numbers at the end of each bar indicate the number of genes in each pathway detected in our data. (**F**) YAP immunofluorescence demonstrates elevated active nuclear YAP in epithelial cells form EO CRC patient tissues. Staining of matched normal and cancer tissue samples using active (non-phosphorylated) YAP was followed by quantification of nuclear intensity for active YAP. (**G**) Gene Set Enrichment Analysis (GSEA) confirms upregulation of an established YAP signature.^[71]^ Normalized enrichment score (NES = 1.98) and *P* value (*P* = 0.025) demonstrate significant upregulation of YAP activity in EO CRC. Data are presented as violin plots with data points indicating individual ROIs. Statistical significance was assessed using a Scheirer-Ray-Hare test with Dunn’s post-hoc test and Bonferroni correction. * indicates significant differences (*P* < 0.05) between cancer and matched normal tissues within onset groups (AO, EO) and between onset groups within tissue types (matched normal, cancer).

GO analysis of epithelial cells revealed the enrichment of pathways related to cell division, mitotic processes, and metabolism in EO cancer (Figure 5C). Notably, the stem cell marker *OLFM4* was highly upregulated in EO CRC while another stem cell marker *LGR5* was highly downregulated in EO CRC (**Figure S14**). To confirm the observed trends in transcriptional data based on a limited number of epithelial ROIs, we conducted immunohistochemistry (IHC) staining of tissue samples using the cell proliferation marker Ki67 and found evidence of increased cell proliferation in both EO matched normal and EO cancer samples (Fig 5D). We noted that epithelial cells in EO CRC had a distinct pattern of increased proliferation, which mirrored the observed trends in tissue stiffness (cf. Figure 1). These data support the hypothesis that epithelial proliferation may be mechanically regulated in EO CRC.

We next analyzed mechanotransductive signaling using eight major mechanotransduction categories. We found unexpected downregulation of genes related to mechanosensitive transcription factors (Mechano TFs), mechanosensors, cytoskeletal components, canonical Hippo pathway, and Rho signaling in EO CRC (Figure 5E). Conversely, genes involved with focal adhesion complexes, integrins, and the linker of nucleoskeleton and cytoskeleton (LINC) complex demonstrated upregulation in EO CRC (Figure 5E), suggesting a complex set of alterations in the epithelial mechanosensing machinery. The selected mechanotransduction pathways did not reach statistical significance at an adjusted *P* value of 0.05 (Figure 5E). However, individual mechanotransduction genes within these pathways exhibited substantial expression changes (**Figure S15**, |log2 Fold Change| > 1). Given the established role of the Hippo pathway in regulating the proliferation of intestinal epithelial cells,^[70]^ the downregulation of *YAP1* (YAP) and *WWTR1* (TAZ), along with elevated expression of *LATS1* surprised us (**Figure S16**A). Furthermore, the TEA-domain (TEAD) transcription factors *TEAD1-4* displayed a mixed pattern of expression (Figure S16B), with upregulation in genes downstream of the Hippo pathway (Figure S16C). This findings suggested that the mechanosensitive effectors YAP/TAZ may be active in epithelial cells and partially responsible for the observed pattern of increased cell proliferation, while the canonical Hippo pathway is downregulated in EO CRC. To reconcile this apparent contradiction, we examined active (non-phosphorylated) YAP in tissue samples via immunofluorescence (IF). We found evidence of greater nuclear YAP activity in epithelial cells from both EO matched normal and EO cancer samples than in epithelial cells from their AO counterparts (Figure 5F). This finding indicated that YAP activity in EO CRC is higher than in AO CRC despite the downregulated canonical Hippo signaling, which suggests that YAP may be activated through Hippo-independent mechanisms, such as integrin-mediated mechanotranscduction. We conducted a GSEA using a validated YAP transcriptional signature^[71]^ and confirmed upregulation of YAP activity in EO CRC epithelial cells (Figure 5G). Taken together, these data support a model in which fibrotic remodeling in EO CRC directly contributes to epithelial activation and proliferation via YAP-mediated mechanotransduction.

### Microenvironmental stiffness enhances epithelial proliferation through YAP activity

To isolate the contribution of mechanical cues to epithelial cell proliferation, independent of other in vivo factors, we used both two-dimensional (2D) and three-dimensional (3D) in vitro models. First, we utilized CRC cell lines cultured on polyacrylamide (PA) substrates of graded stiffness^[72–74]^ to investigate mechanobiological regulation of CRC cell proliferation. Based on the material stiffness values measured from primary patient samples (Figs. 1F and 1I), we restricted our analysis to relatively moderate substrate stiffnesses, which differed from other studies.^[75–77]^ The soft hydrogels used to replicate healthy tissues displayed a Young’s modulus of 0.1-0.8 kPa while the stiff hydrogels used to replicate fibrotic tissues displayed a Young’s modulus of 3.2-4.9 kPa (**Figure S17**). The colon cancer cell lines HT29 (44-year-old patient, used as a model of EO CRC) and SW480 (50-year-old patient, used as a model of AO CRC) were seeded on collagen- and Matrigel-coated PA gels of graded stiffness for 72 hours (**Figure S18**). We measured the cellular aspect ratio, percentage of EdU-positive cells, and nuclear intensity of active YAP. We found that SW480 cells had greater mechanoresponsiveness than the HT29 cells in both morphology and proliferation (Figure S18). Importantly, SW480 cells cultured on collagen-coated PA gels, which reproduces the stromal environment, remained highly proliferative and elongated regardless of substrate stiffness (Figure S18B). Conversely, SW480 cells cultured on Matrigel-coated PA gels, which reproduces the native basement membrane, exhibited stiffness-dependent changes in morphology and proliferation as a function of increasing substrate stiffness (Figure S18B). In both cell lines, YAP activity appears to be modulated by the substrate stiffness rather than by the specific ECM ligand (Figure S18). Therefore, our data suggests that mechanosensitive differences in CRC cell behavior occur exclusively on Matrigel-coated substrates within the stiffness range characteristic of EO CRC pathophysiology.

To better understand the mechanobiological regulation of cell proliferation, we focused on the mechanisms underlying stiffness-dependent changes in CRC cell lines cultured on Matrigel-coated PA substrates (**Figure 6**). Across multiple biological replicates, HT29 cells displayed modest changes in aspect ratio and EdU incorporation with increasing stiffness, with significant changes in morphology and proliferation occurring only on glass (Figure 6C and 6E). Instead, SW480 cells exhibited increasingly elongated morphology and enhanced proliferation with increasing stiffness, as indicated by higher aspect ratios and EdU incorporation on both stiff gels and glass (Figure 6D and 6F). Both cell lines revealed higher YAP activity only on glass (Figure 6C-F), consistent with previously reported rigidity thresholds.^[78]^ To clarify whether YAP activity regulates the observed stiffness-dependent changes, we used verteporfin (VP) to block YAP-TEAD interactions.^[79]^ We found that VP concentrations of 1-2 μg/mL preserved cell viability while reducing the cell aspect ratio and inducing cell rounding (**Figure S19**). Treatment with these VP concentrations on substrates matching the stiffness of primary CRC tissues confirmed that YAP activity mediates the stiffness-dependent changes in both cell morphology (Figure 6G-H and **S20**) and proliferation (Figure 6I-J and S20) in HT29 and SW480 cell lines. Therefore, despite differences in mechanosensitive responsiveness across ECM coatings and cell types, elevated substrate stiffness promoted cancer cell proliferation through YAP activity in both HT29 and SW480 models.

**Figure 6.**
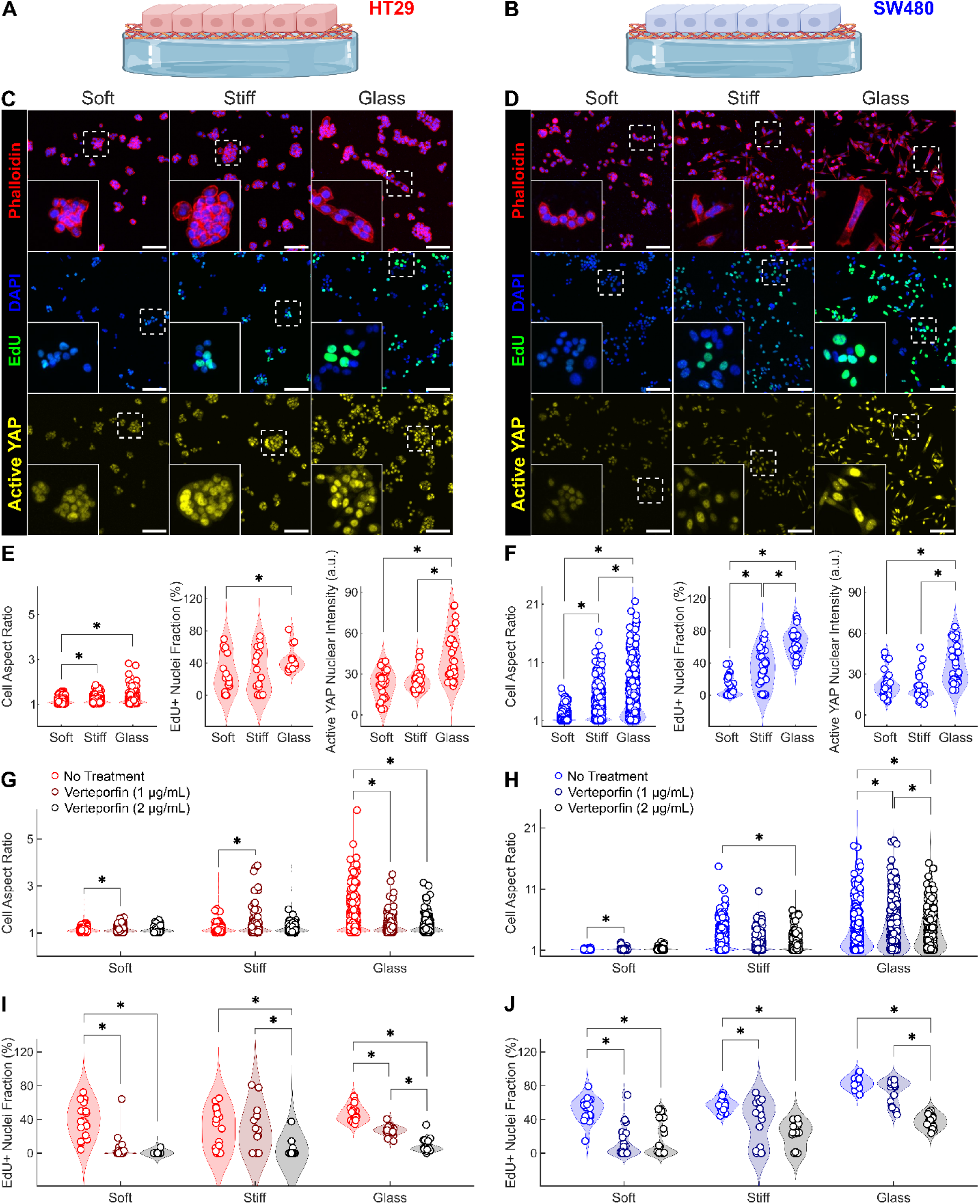
2D microenvironmental stiffness alters epithelial morphology and proliferation via YAP activity in CRC cell lines. Schematic representation of (**A**) HT29 cells (44-year-old female) and (**B**) SW480 cells (50-year-old male) cultured on 2D polyacrylamide (PA) substrates of graded stiffness (cf. Figure S17A) coated with Matrigel to investigate mechanobiological regulation of CRC cells. Immunofluorescence (IF) staining for phalloidin (red), DAPI (blue), EdU (green), and active YAP (yellow) on (**C**) HT29 and (**D**) SW480 cells cultured on graded substrate stiffness for 72 hours. Glass served as a positive control for high stiffness. Scale bars, 100 μm. (**E**) Quantification of HT29 cell morphology, proliferation, and YAP activity from IF images. The aspect ratio of HT29 cells (n = 543 for soft PA, n = 631 for stiff PA, and n = 736 for glass) was quantified using 6 biological replicates and 6 ROIs per replicate. EdU incorporation in HT29 cells (n = 17 for soft PA, n = 18 for stiff PA, and n = 15 for glass) was quantified using 3 biological replicates and 5-to-6 ROIs per replicate. The fluorescence intensity of active YAP in HT29 cell nuclei (n = 30 for soft PA, n = 30 for stiff PA, and n = 30 for glass) was quantified using 5 biological replicates and 6 ROIs per replicate. (**F**) Quantification of SW480 cell morphology, proliferation, and YAP activity from IF images. The aspect ratio of SW480 cells (n = 771 for soft PA, n = 769 for stiff PA, and n = 659 for glass) was quantified using 6 biological replicates and 6 ROIs per replicate. EdU incorporation in SW480 cells (n = 30 for soft PA, n = 36 for stiff PA, and n = 36 for glass) was quantified using 5-to-6 biological replicates and 6 ROIs per replicate. The fluorescence intensity of active YAP in SW480 cell nuclei (n = 30 for soft PA, n = 30 for stiff PA, and n = 30 for glass) was quantified using 5 biological replicates and 6 ROIs per replicate. (**G, H**) Changes in cell morphology between control (NT = No Treatment) and verteporfin-treated (VP1 = 1 µg mL^−1^, VP2 = 2 µg mL^−1^) HT29 cells (n = 380 for NT, n = 341 for VP1, n = 315 for VP2 on soft PA; n = 447 for NT, n = 441 for VP1, n = 305 for VP2 on stiff PA; n = 480 for NT, n = 450 for VP1, n = 364 for VP2 on glass) and (**H**) SW480 cells (n = 321 for NT, n = 372 for VP1, n = 716 for VP2 on soft PA; n = 510 for NT, n = 348 for VP1, n = 807 for VP2 on stiff PA; n = 407 for NT, n = 428 for VP1, n = 720 for VP2 on glass) were quantified using 3 biological replicates and 4-to-6 ROIs per replicate. (**I, J**) Associated changes in cell proliferation upon verteporfin treatment of HT29 cells (n = 18 for NT, n = 18 for VP1, n = 18 for VP2 on soft PA; n = 18 for NT, n = 16 for VP1, n = 18 for VP2 on stiff PA; n = 18 for NT, n = 18 for VP1, n = 18 for VP2 on glass) and SW480 cells (n = 18 for NT, n = 18 for VP1, n = 18 for VP2 on soft PA; n = 18 for NT, n = 18 for VP1, n = 18 for VP2 on stiff PA; n = 17 for NT, n = 17 for VP1, n = 18 for VP2 on glass). Data are presented as violin plots with data points indicating individual measurements. Statistical significance was assessed using Scheirer-Ray-Hare test followed by Dunn’s post-hoc test with Bonferroni correction. * indicates statistically significant differences (*P* < 0.05) between groups. Schematics created using Biorender.com.

To assess the role of 3D mechanosensitivity using clinically relevant models, we used established methods^[80–82]^ to derive epithelial organoids from a subset of patients within our patient cohort (Table S1 and S2). Organoids from unique AO and EO CRC patients (**Table S3**) were cultured in hyaluronic acid-tyramine (HA-Tyr) hydrogels^[82]^ of different concentrations (2.5 mg mL^−1^ and 8 mg mL^−1^), which corresponded to different stiffness values (Figure S17). After 7 days of culture, organoids in stiff hydrogels had larger sizes regardless of the tissue of origin (AO vs. EO) (Figure 7B). We used organoid viability as a proxy for proliferative capacity and found that changes in viability accompanied the observed changes in organoid size (Figure 7C). These in vitro findings demonstrate that increased microenvironmental stiffness may facilitate epithelial proliferation, independent of the underlying mutational status or age of the tumor of origin. These results support our ex vivo observations and suggest that microenvironmental stiffening may directly contribute to epithelial activation and facilitate tumor progression in fibrotic EO CRC. **DISCUSSION**

**Figure 7.**
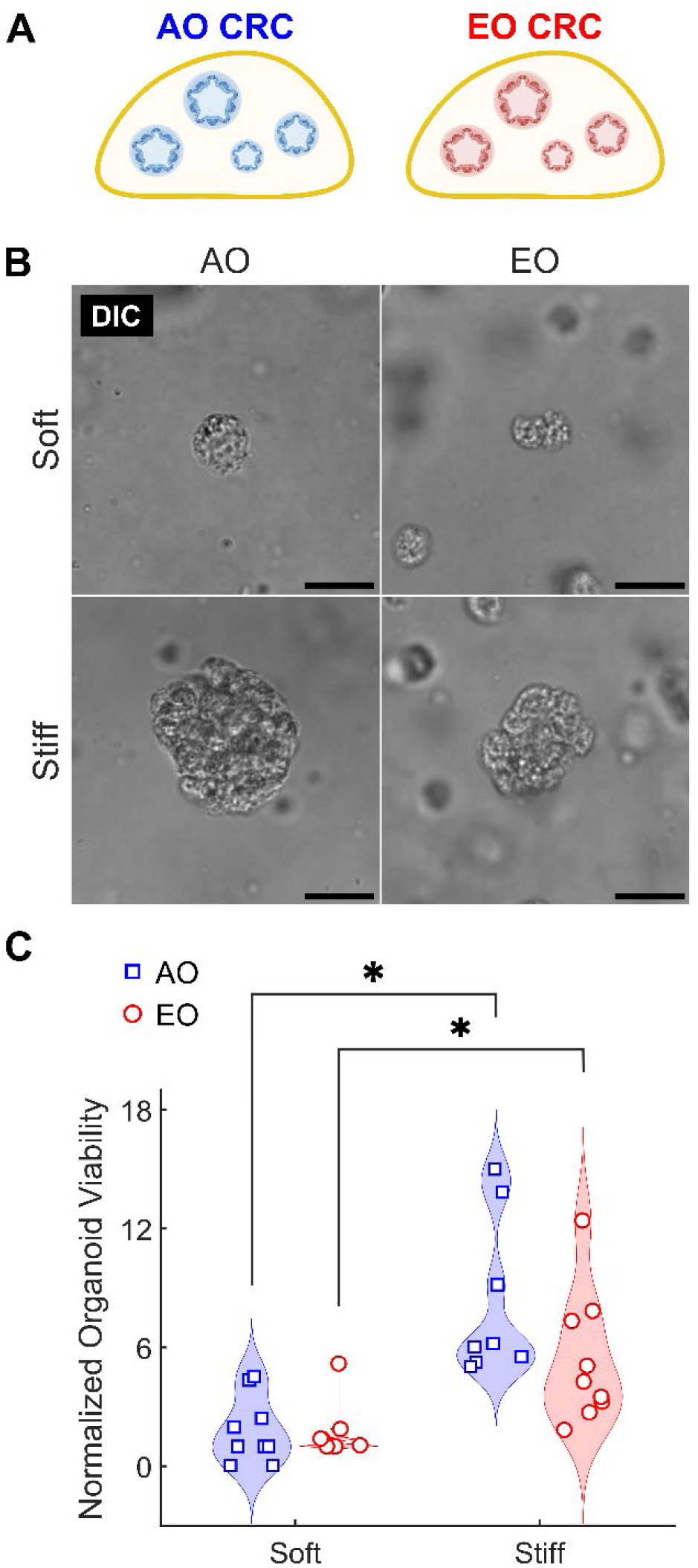
3D microenvironmental stiffness enhances viability of patient-derived CRC organoids. (**A**) Schematic representation of 3D organoids derived from AO and EO CRC patients cultured in hyaluronic acid-tyramine (HA-Tyr) hydrogels with tunable stiffness (cf. Figure S17B). (**B**) Representative DIC images of patient-derived organoids from AO and EO CRC patients cultured in soft and stiff HA-Tyr hydrogels for 7 days. Scale bars, 20 μm. (**C**) Quantification of normalized organoid viability in soft and stiff hydrogels using CellTiter-Glo® 3D assay. Viability increases with substrate stiffness in both patient groups. Each group (AO CRC and EO CRC) included 3 biological replicates (patient donors) and 3 technical replicates per donor (3 organoid-seeded hydrogels), resulting in a total of n = 9 per group. Data are presented as violin plots with data points indicating individual measurements. Statistical significance was assessed using Scheirer-Ray-Hare test followed by Dunn’s post-hoc test with Bonferroni correction. * indicates statistically significant differences (*P* < 0.05) between groups. Schematics created using Biorender.com.

Although the incidence and mortality of EO CRC are rapidly increasing worldwide,^[83]^ its etiology remains poorly understood.^[5,9]^ Because most cases of sporadic EO CRC are multifactorial, involving both environmental and lifestyle factors,^[18]^ there is a pressing need to uncover mechanistic insights and to identify predictive biomarkers capable of stratifying patients at high risk for metastatic disease and recurrence, particularly among stage II and III patients. In this study, we identify tissue stiffening and fibrotic remodeling as consistent biophysically measurable hallmarks of EO CRC. Furthermore, we link these mechanical changes to collagen remodeling in the colonic stroma and to mechanobiological changes in the colonic epithelium (Figure 8).

**Figure 8.**
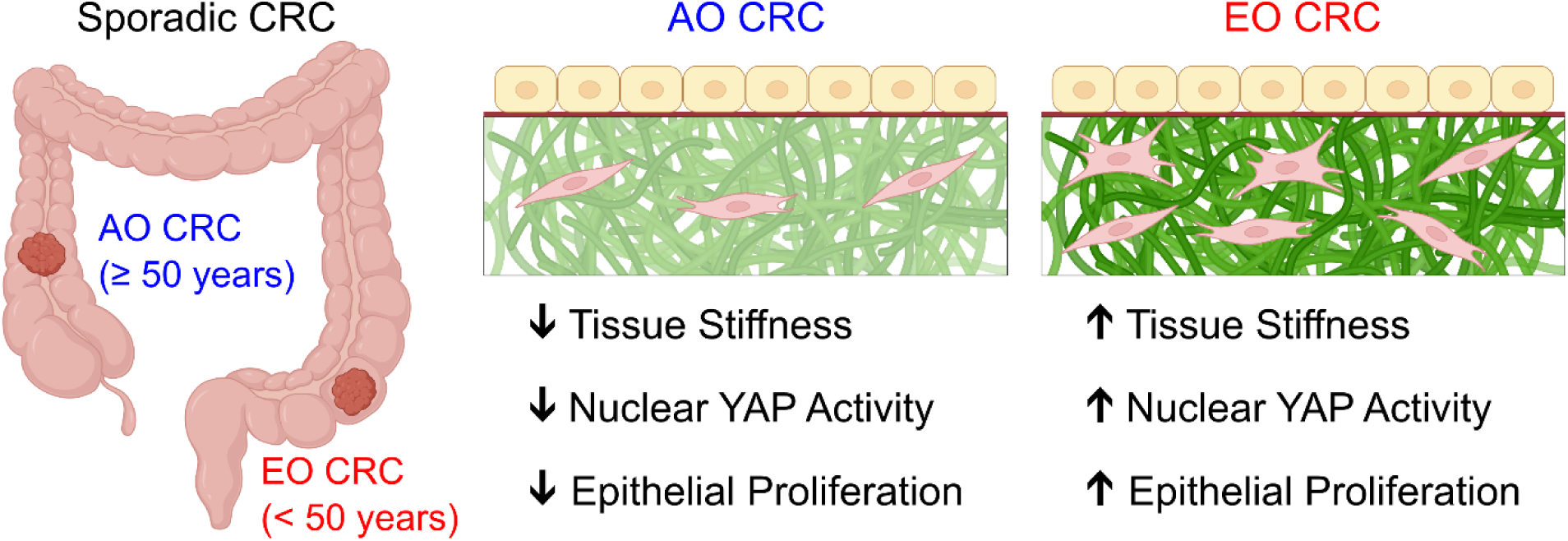
Summary of biomechanical and mechanobiological hallmarks of EO CRC. Using primary patient samples, our study quantified biomechanical, histological, transcriptional, and phenotypic differences between AO CRC and EO CRC. We find that EO CRC is associated with changes in collagen organization due to the activity of stromal cells, which results in increased tissue stiffness as measured via mesoscale indentation and macroscale compression. These changes in connective tissue mechanics result in increased nuclear activity of YAP and increased epithelial proliferation. Created using Biorender.com.

We hypothesized that fibrosis and mechanical stiffening of colorectal tissues serve both as early and active drivers of EO CRC progression. Based on this premise, our first objective was to compare tissue biomechanics between EO CRC and AO CRC patient samples. Although several reports have examined altered tissue mechanics in primary^[35–38,41]^ and metastatic^[84,85]^ CRC, most relied on atomic force microscopy (AFM), which probes tissue stiffness at nano- and microscale resolution but requires sample processing steps including adhesive mounting^[41]^, dehydration^[36]^, and cryo-sectioning^[37,38,84,85]^ that may alter the native tissue mechanics. Additionally, AFM captures localized mechanical properties of cell and ECM structures rather than the broader tissue-level mechanical properties relevant for diagnostic and therapeutic applications. For these reasons, here we used mesoscale indentation^[86]^ and macroscale compression,^[43]^ both of which require minimal tissue preparation and enable evaluation of bulk mechanical behavior. Our multiscale mechanical measurements of primary CRC tissues yielded stiffness values under 3 kPA, which represents an average of the heterogeneous mechanical landscape that AFM resolves at higher resolution, including soft cellular regions (<1 kPa) and stiffer ECM-rich structures (∼10-20 kPa). Differences between normal and cancer tissues were more modest than previously reported^[35,41,37]^ possibly due to patient heterogeneity or differences in testing methodology. Notably, both mesoscale and macroscale approaches revealed significantly elevated stiffness in EO CRC tissues relative to AO CRC (Figure 1).

While tissue stiffness – quantified as the Young’s elastic modulus – can be extracted from both mesoscale indentation and macroscale compression under the assumption of linear elastic behavior, characterizing nonlinear elastic properties and viscoelastic behavior requires large deformations and time-resolved measurements, respectively. To quantify the nonlinear viscoelastic properties of CRC tissues, we adopted stress-relaxation testing in macroscale compression paired with a nonlinear Maxwell-Wiechert model.^[44,45]^ Our data revealed increased tissue viscosity as an additional biophysical signature of EO CRC (Figure 1 and S4), a distinct biomechanical feature not previously reported. Given the emerging recognition of matrix viscoelasticity as a critical regulator of cellular and tissue dynamics,^[87,88]^ we propose that this observed increase in viscosity in EO CRC represents an important mechanobiological characteristic that warrants further investigation.

Strikingly, we found that not only cancerous tissues but also normal tissues from EO patients exhibited elevated stiffness and higher viscosity than tissues from AO patients (Figure 1), suggesting that mechanical remodeling may precede malignant transformation and may contribute to the development of EO CRC. Indeed, quantitative PSR staining and POL imaging revealed that EO normal tissues were enriched with red (thicker, most tightly packed and mature) collagen fibers, while they displayed less orange and yellow (intermediate thickness and packing) collagen fibers, with minimal green (thinnest, loosely packed and less mature) collagen fibers (Figure 2 and S6). By examining the microstructural organization of collagen via multiphoton SHG imaging, we found that EO normal tissues were characterized by an increased SHG intensity, a reduced fiber length, and similar fiber width and orientation with respect to AO normal tissues (Figure S7). Collectively, these results reveal that normal tissues from EO patients contain shorter, denser, and therefore more tightly packed collagen fibers, which are likely responsible for the observed increase in the stiffness of matched normal tissues. Although it is possible that matched normal tissues (retrieved 5 cm away from the tumor) were impacted by pre-neoplastic changes, the structural and mechanical differences observed between AO and EO normal tissues raised the possibility that fibrotic remodeling and tissue stiffening may represent microenvironmental cues that predispose malignant transformation of a normal epithelium in EO CRC. To investigate the structural causes of altered tissue mechanics in cancer samples, we combined quantitative histological and immunofluorescence analyses (Figure 2 and 3). EO cancer tissues exhibited thinner, longer, and more aligned collagen fibers than AO cancer tissues (Figure S5-S7). At the same time, EO cancer tissue exhibited elongated stromal cells and overall greater expression of the mesenchymal marker Vimentin (Figure S8-S10). Notably, the architectural remodeling of collagen in EO cancer resembles the radially aligned tumor-associated collagen signatures observed in breast cancer,^[89]^ which correlates with increased invasiveness and poor prognosis.^[56]^ This parallel suggests that collagen alignment by stromal cells may provide a biophysical mechanism to explain the aggressive, invasive behavior characteristic of EO CRC.

Since collagen remodeling in the stroma is primarily driven by CAFs, and fibroblast activation protein (FAP)-positive CAFs are enriched at the invasive margin of human EO CRC tissues,^[90]^ we used spatial transcriptomic analysis to explore the transcriptional signatures of Vimentin-positive stromal cells (Figure 3). Although canonical CAF markers (*ACTA2*, *FAP*, *PDGFA*, *PDGFB*) were not selectively enriched in our samples, we observed upregulation in ECM remodeling genes in EO CRC. The most significantly upregulated genes were associated with ECM crosslinking (*LOXL1/2*, *TGM2*), MMP inhibition (*TIMP1*), and increased cell-ECM interactions (*THBS1*, *ITGA5*, *CD44*), thereby supporting the accumulation of a stiffer and more aligned ECM in EO CRC (Figure 4 and S12). In parallel, elevated expression of pro-inflammatory mediators (*CXCL1*, *CCL2*) and immunomodulatory ECM proteins (*COL6A1*) support the presence of a pro-inflammatory, immune-recruiting stromal environment.^[91]^ In PanCK-positive epithelial cells, spatial transcriptomics revealed upregulation of genes involved in cell proliferation and metabolism in EO CRC (Figure 5A-D). Among genes involved in cell proliferation, the stem cell marker *OLFM4* was significantly upregulated while another stem cell marker *LGR5* was significantly downregulated in EO CRC. Notably, the same pattern in stem cell marker expression was independently observed in intestinal stem cells cultured on substrates of increasing stiffness.^[75]^ At the same time, it has been shown that LGR5-negative colon cancer stem cells are characterized by reduced stiffness and increased motility than their LGR5-positive counterparts.^[77]^ In addition, we observed upregulation of the neuronal pentraxin 2 (*NPTX2*) gene by epithelial cells (Figure 5B), which has been associated with proliferation, epithelial-to-mesenchymal transtion, and metastasis in CRC.^[92]^ Together, our findings suggest that epithelial cells in EO CRC may be more responsive to mechanical cues from the local tissue microenvironment. While it is known that matrix stiffness mediates stemness characteristics in CRC,^[93]^ mechanobiological investigations focused on the functional role played by LGR5 and OLFM4 could provide further insights into EO CRC pathogenesis and progression. Given emerging evidence linking mechanical signaling to stem cell behavior and cancer development, studies examining whether these markers influence cellular responses to mechanical cues – including effects on metabolism and proliferation – may reveal novel mechanistic pathways relevant to EO CRC.

Analysis of the expression patterns associated with AO and EO CRC epithelial cells show that cancer samples within our patient cohort did not segregate into the intrinsic subtypes iCMS2 and iCMS3 (Figure S13). This finding further supports that AO and EO CRC may not be distinguished based on intrinsic epithelial features but rather based on microenvironmental factors. Indeed, we found that epithelial proliferation in EO CRC tissues is associated with enhanced nuclear YAP activity (Figure 5). Despite our finding that canonical Hippo signaling was transcriptionally downregulated in EO CRC (Figure 5E and S16), immunofluorescence revealed greater nuclear localization of active YAP protein in both matched normal and cancerous EO epithelia (Figure 5F). GSEA confirmed coordinated upregulation of YAP target genes in EO CRC relative to AO CRC (Figure 5G). The seemingly contradictory findings from spatial transcriptomics (downregulated Hippo signaling) and immunofluorescence (active YAP in cell nuclei) of EO CRC epithelial cells may be explained through the ability of integrins, focal adhesions, and the LINC complex^[78]^ to activate YAP in response to fibrotic tissue stiffening independently from activation of the Hippo pathway.^[94]^ In response to YAP activation, negative feedback in the regulatory gene network may be triggered and affects the expression of the Hippo pathway genes, resulting in the observed downregulation of the canonical Hippo pathway (Figure S16). This suggests that enhanced YAP activity in EO CRC epithelial cells likely results from post-transcriptional protein stabilization through integrin signaling and actomyosin contractility,^[95]^ rather than regulation through the canonical Hippo pathway. While striking, it is possible that the observed increase in YAP activity and proliferation in epithelial cells from EO CRC patients may not be caused by alterated tissue mechanics but rather to other confounding factors, such as patient age.^[96]^ For this reason, we moved on to investigate the mechanobiological regulation of colonic cancer cells using both cell lines and primary organoids (Figure 6 and 7).

To directly test the effect of matrix stiffness on epithelial cell proliferation, we examined stiffness-dependent changes in colonic cancer cell morphology and proliferation using both cell lines and primary organoids (Figure 6 and 7). We selected the HT29 and SW480 models as MSS colorectal adenocarcinoma cell lines characterized by distinct genetic profiles^[97]^ and physical properties.^[98]^ When cultured on 2D PA hydrogels, SW480 cells (a model of AO CRC) were more mechanosensitive than HT29 cells (a model of EO CRC), exhibiting increased elongation and proliferation on stiffer substrates (Figure 6A-F and S18). While prior literature demonstrates that cell spreading and proliferation responses to substrate stiffness vary between cell lines,^[99]^ we found that the observed mechanosensitive changes also depend on the ECM coating used to functionalize PA substrates. In fact, stiffness-dependent changes emerged only on Matrigel-coated substrates, while both aspect ratio and proliferation of cells cultured on collagen I-coated substrates were consistently elevated regardless of substrate stiffness (Figure S18). Coating substrates with Matrigel better recapitulates the basement membrane environment of colonic epithelial cells than coating with collagen I and therefore is more relevant as a model of CRC initiation. Importantly, we focused our in vitro analysis on a moderate stiffness range (0.1-to-4.9 kPa) that reflects mechanical properties measured in primary EO CRC tissue samples (Figure 1), with the goal of reproducing the mechanobiological responses of epithelial cells in EO CRC. Other studies reported stiffness-mediated changes in CRC proliferation and motility using collagen-coated PA substrates of stiffnesses up to 38 kPa^[100]^ or 126 kPa.^[95]^ Instead, here we show that stiffness-mediated changes in CRC phenotype can be achieved on Matrigel-coated PA substrates within a pathophysiologically relevant stiffness range. Therefore, our findings emphasize that both ECM composition and stiffness influence the mechanobiology of AO and EO colon cancer cells.

To determine whether YAP activity mediates the increased epithelial proliferation observed in response to tissue stiffening, we employed our 2D PA culture system to interrogate YAP activity in vitro. First, we observed that the fluorescence intensity of active (non-phosphorylated) YAP was elevated only in CRC cells cultured on glass (Figure 6C-F). Our IF data are consistent with a minimal rigidity threshold of ∼5 kPa required for YAP nuclear translocation.^[78]^ However, a recent report identified drastic morphological changes occurring at a rigidity threshold as low as ∼2 kPa, accompanied by changes in YAP activity.^[101]^ To demonstrate that YAP regulates the observed stiffness-dependent changes in CRC cells, we ablated YAP-TEAD interactions using the pharmacological agent verteporfin. We found that YAP inhibition using verteporfin suppressed stiffness-mediated changes in CRC cell morphology and proliferation in both HT29 and SW480 cell lines (Figure 6G-J). Our data are consistent with reports documenting phenotypic and morphological changes upon YAP knockdown on substrates of supraphysiological stiffness.^[95]^ Functionally, YAP is known to confer a proliferative and invasive phenotype to CRC cells,^[102]^ potentially explaining how tissue stiffening facilitates EO CRC initiation and malignancy. Overall, our in vitro data using CRC cell lines cultured on PA hydrogels align well with our observations from patient tissues and further provide a mechanistic link between tissue stiffening, YAP activity, and CRC proliferation.

Finally, we validated the impact of microenvironmental stiffness on cancer cell proliferation using 3D primary organoids (Figure 7) and found that hydrogel stiffness dictates proliferation rather than the tissue of origin (AO vs. EO CRC). In fact, patient-derived organoids cultured for 7 days in stiff hydrogels were consistently more proliferative than matched organoids cultured for the same period in soft hydrogels. This observation validates our proposed mechanism by showing that the differences in epithelial cell proliferation observed in human tissues (Figure 5D) can be reproduced by tuning solely the mechanical properties of the TME (Figure 7C).

The primary limitation of this study is a small sample size, comprising 19 AO CRC and 14 EO CRC tissue pairs (each containing matched normal and cancer samples). Additionally, some patients did not yield complete biomechanical datasets due to insufficient tissue size and lack of visible tissue layers, or damage during preparation and testing. Our sample set also had a notable absence of rectal cancers in the AO CRC group, likely reflecting more aggressive chemoradiation therapy that reduces tumor size and requires prioritizing tissue for essential clinicopathological analyses over research use. Furthermore, we performed spatial transcriptomic analysis on a limited number of ROIs and it was not consistently conducted on both matched normal and cancer tissues within PanCK-positive ROIs. Finally, it should be noted that the matched normal tissues were obtained from areas adjacent to the tumor and may exhibit field effects that could influence our comparative analyses. Despite these constraints, a coherent mechanobiological framework emerges from our multiscale approach integrating biomechanical measurements, quantitative histology, cellular phenotype analysis, spatial transcriptomics, and in vitro modeling. To the best of our knowledge, this work therefore represents the first comprehensive mechanobiological characterization of EO CRC. We demonstrate that pro-inflammatory and pro-fibrotic remodeling drives tissue stiffening, leading to enhanced epithelial YAP activity and proliferation. We propose that these mechanobiological signatures play a key role in promoting EO CRC development and progression (Figure 8).

Future investigations should focus on three key areas: (1) elucidating the biochemical and biomechanical crosstalk between stromal and epithelial cells during EO CRC initiation and progression, (2) systematically modulating both microenvironmental stiffness and viscosity to define their respective contributions to epithelial stemness, phenotypic plasticity, and invasion, and (3) evaluating compositional and mechanical biomarkers as early predictors of EO CRC progression for improved diagnostic and therapeutic strategies.

## MATERIALS AND METHODS

### Patients

We obtained tissue specimens from patients who underwent operative resection for colorectal cancer at either the Cleveland Clinic (IRB 13-1159) or at the University of Texas Southwestern Medical Center (IRB 2021-0161). Written informed consent was obtained from all patients for the collection and use of tissue samples for research purposes. The inclusion criteria were as follows: individuals aged above 18 years, decisionally competent, non-pregnant, non-incarcerated, with colorectal cancer undergoing exploration with resection of bowel. Patient demographics, stage, and cancer location are summarized in Table S1 and S2. Our analysis focused on patients with MSS colorectal tumors, which we confirmed using standard immunohistochemistry for MLH1, MSH2, MSH6, and PMS2. At the time of resection, we also retrieved matching normal tissues at least 5 cm away from the tumor. We separated tumor and matching normal tissues into specimens for mechanical testing, histological analysis, and organoid preparation. We stained FFPE tissue sections with H&E, and a board-certified pathologist (ZC) in the Department of Pathology at UT Southwestern evaluated them.

### Multiscale biomechanical testing

We froze freshly excised specimens for ex vivo mechanical evaluation at -80°C and stored them until testing. Preliminary experiments ensured that fresh and frozen tissues were mechanically indistinguishable. After thawing each patient sample in a water bath (37°C), we used a biopsy punch to isolate two to three cylindrical specimens (3 mm in diameter) per sample. We subjected the resulting tissue plugs to mechanical testing using a commercial system (Cell Scale, Waterloo, ON, Canada) and established protocols^[42,43]^ to determine their material properties at different length scales. We used MATLAB R2024b (Mathworks, Natick, MA) to carry out all calculations, group averaging, plotting, and biomechanical modeling , unless otherwise specified. We subjected each specimen to four local indentation tests and to one global unconfined compression. In both indentation and compression testing, we estimated the forces generated by tissues by measuring the deflection of a microbeam of known elastic modulus (411 GPa) and length (57–59 mm). The microbeam diameter Φ dictates its overall force resolution. Therefore, we used a smaller diameter for local indentation tests (Φ = 0.0762, 0.1016, or 0.1524 mm) while we used a larger diameter for global compression tests (Φ = 0.3048, 0.4064, or 0.5588 mm). For each specimen, we determined the precise microbeam diameter Φ that best matches individual tissue properties via preliminary tests. Prior to each experiment, we incubated the indenter and compression platen in 1% bovine serum albumin for 30 minutes to minimize tissue adhesion.

#### Local indentation

We fixed a short microbeam of diameter Φ = 0.0762 mm at the end of the longer force-sensing microbeam in a 90° configuration to create a cylindrical indenter (cf. Figure 1C-D). We performed indentation testing in 5 incremental steps, each with a magnitude equal to 1% of the original height of the sample and interspersed with 1-minute hold phases (Figure S2A-D). We plotted the steady-state force (*F*) from the hold phases against the local tissue displacement (*δ*) and used these graphs to fit the classical Sneddon’s solution from linear elasticity,^[103,104]^

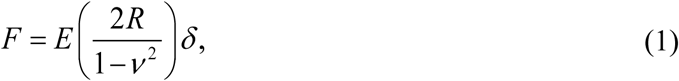

where *E* represents the unknown Young’s modulus, *R* = 38.1 µm is the radius of the cylindrical indenter and *ν* = 0.5 is the Poisson’s ratio for an incompressible linear material. Using this method the calculated values of *E* indicate the local material properties of human colorectal tissues.

#### Global compression

We fixed a square platen with area 6 mm × 6 mm to the force-sensing microbeam, thereby enabling compression of entire specimens (Figure S3A-C). Since the mechanical behavior of soft tissues is pseudoelastic,^[105]^ we used a preconditioning protocol involving 10 cycles of compression up to a deformation of 20% of the initial height of the cylindrical specimen with a strain rate of 0.5 % s^−1^ to yield consistent behaviors under loading. Unconfined compression testing was subsequently performed in 8 incremental steps, each with a magnitude equal to 2.5% (for a total of 20% deformation) interspersed with 2-minute hold phases (Figure S3D). In addition to enabling force estimation by measuring the microbeam deflection, the side camera allowed us to quantify the specimen geometry (i.e., height and diameter) at each time point during the test (Figure S3E). Measurements of initial height ( *H*_0_ ) and radius ( *R*_0_ ) of the undeformed specimen and of current tissue height ( *h_c_* ) and radius ( *r_c_* ) of the deformed specimen allowed us to calculate radial and axial strain components as *ε_rr_* = (*r_c/_ R*_0_ ) −1 and *ε _zz_* = (*h_c_* / *H*_0_ ) −1 , respectively. Finally, the axial Cauchy stress was calculated as *σ* = *F* (*π r_c_*^2^ ) . We determined the true Poisson’s ratio *ν* = −*ε_rr_* /*ε_zz_* by finding the best fit line between the axial and radial strain (Figure S3F). We fitted steady-state values of the equilibrium Cauchy stress (Figure S3G-H) using a neo-Hookean strain energy density function,^[106]^

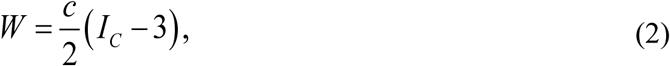

where *c* represents an unknown material parameter, and *I_C_* represents the first invariant of the Cauchy-Green tensor. Details regarding parameter estimation via nonlinear least square minimization can be found elsewhere.^[42]^ To enable comparisons between local and global experiments, we determined the global Young’s modulus as *E* = 2*c* (1+*ν* ) , where the values for *c* and *ν* where independently estimated for each specimen from unconfined compression data.

#### Viscoelastic modeling

To fully capture the nonlinear stress-relaxation behavior displayed by colorectal tissues under compression, we implemented a modified Maxwell-Wiechert model.^[44,45]^ Such model consists of a nonlinear spring in parallel to 2 Maxwell elements, each consisting of a linear spring and a linear dashpot, as shown in the model schematic (Figure S4A-B). According to this model, the time-dependent stress response of tissue plugs under compression can be described by the following equation,

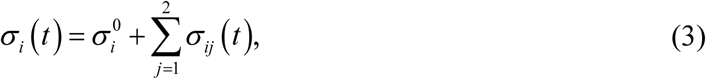

where *i* = 1,…,8 indicates each of the loading steps and *j* = 1, 2 indicates each of the Maxwell elements. The stress response of the nonlinear spring *σ*_i_^0^ is described by the following equation,

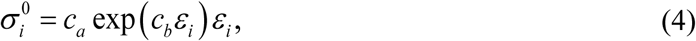

where *c_a_* is a stress-like material parameter quantifying the stiffness of the nonlinear spring and *c_b_* represents a dimensionless parameter indicating the nonlinearity of the stress-strain response. The symbol *ε_i_* indicates the discrete values taken by the axial strain component *ε _zz_* at the end of each stress-relaxation step *i*. Finally, the stress response by the Maxwell elements *σ _ij_* (*t* ) is described by the following equations, valid for each compression step ( *t_i_* ≤ *t* ≤ *t_i_*_+1_ ):

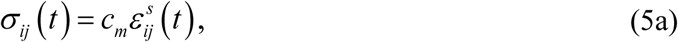

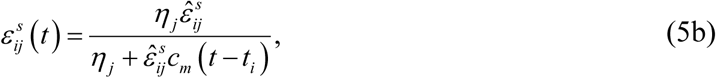

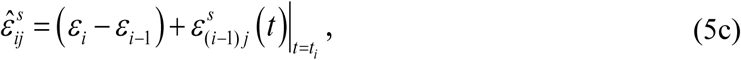

where *c_m_* is a stress-like material parameter quantifying the stiffness of the linear spring while *η _j_* (with *j* = 1, 2 ) are the coefficient of viscosity (with units of stress × time) for standard dashpots indicating the amount of viscous damping. Following Tuttle at al.,^[45]^ we assumed that the 2 Maxwell elements have the same stiffness *c_m_* but different viscosities *η*_1_ and *η*_2_ . The symbol 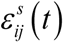 indicates the strain experienced at the *i*-th compression step by the spring within the *j*-th Maxwell element. Equation (5) was solved under the following initial conditions: 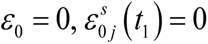 . Details regarding model parameters and their numerical estimation can be _1_ found elsewhere.^[45]^ Briefly, we used a two-step fitting procedure: we first determined the parameters for the nonlinear spring ( *c_a_* and *c_b_* ) by fitting equilibrium data (Figure S4C), then determined the parameters for the Maxwell elements ( *c_m_* , *η*_1_ , and *η*_2_ ) by fitting transient responses (Figure S4D).

### Quantitative histology

Histological examination of human colorectal tissues included using FFPE blocks sectioned at 4 µm and staining them with MTC and PSR. We imaged histology slides, using a Nikon TI2-E inverted microscope equipped with a DS-RI2 color CMOS camera (Nikon, Tokyo, Japan) and custom analyzer and polarizer filters. Complete cross-sectional views of the stained tissues were obtained by acquiring images using a 4× objective and by stitching individual images using the NIS-Elements software. We obtained tiled images using the following settings: 2424 × 2424 pixels at a resolution of 0.734 µm pixel^−1^, and a 15% overlap. For each image, we cropped five representative regions (1 mm × 1 mm) and processed them independently. We analyzed the resulting images using a custom MATLAB script described elsewhere^[48,49,107]^ to quantify the area fractions of collagen from MTC images and to identify the fraction of thick (red), intermediate (orange-yellow), or thin (green) birefringent collagen fibers from PSR images. Briefly, image analysis first consisted of global thresholding to saturate the background to a uniform white (for MTC staining) or black (for PSR staining) value and to eliminate variations in background illumination. We performed pixel-specific thresholding in the HSL (hue–saturation– lightness) color space. Specifically, we classified blue pixels from MTC images (H = 150 – 249°, S = 0 – 1, and L = 0.01 – 1) as total collagen. We identified collagen of varying thickness from PSR images by separating the hue of red (H = 324 – 12°), orange (H = 13 – 52°), yellow (H = 53 – 72°), and green (H = 73 – 180°) pixels while keeping saturation and lightness fixed (S = 0.1 – 1, and L = 0.1 – 0.93). We report results in terms of percent area fractions, by computing the ratio of positively identified pixels to total number of pixels in the tissue sections.

### Multiphoton microscopy

We captured the architecture of fibrous collagen by imaging H&E-stained slides with a FV MPE-RS multiphoton microscope (Olympus, Tokyo, Japan) in the UT Dallas Imaging & Histology core. The laser beam (Mai-Tai HP DeepSee, Spectra Physics, Santa Clara, CA) was focused on the slides through a 25× water-immersion objective (Olympus, 1.05 N.A., 2 mm working distance) mounted in upright configuration. Following prior work,^[42,43]^ we used an excitation wavelength of 880 nm and collected the SHG signal from the collagen matrix using a 410-455nm bandpass filter. Acquisition of tiled image stacks used the following settings: 512 × 512 pixels at a resolution of 0.994 µm pixel^−1^, a pixel dwell time of 2 µs, a stack size of 30 µm with 5 µm steps, and a 10% overlap. We analyzed the resulting SHG images of whole tissue cross-sections using the ImageJ plugin Ridge Detection^[55]^, alongside a custom written MATLAB script. For each image, we cropped five representative regions (1 mm × 1 mm) and processed them independently. For each region, we calculated the average signal intensity by averaging the SHG signal intensity across all pixels. Then we converted images to an 8-bit format and analyzed them using the following setting for Ridge Detection: line width = 3, high contrast = 100, low contrast = 10, lower threshold = 0.34, upper threshold = 4.25, minimum line length = 0, maximum line length = 100. Individual fibers in the regions were detected as ridges, and the associated fiber length and width were calculated. We reported the mean fiber length and width per image (Figure S7). To quantify fiber orientation, we defined an Orientation Index (*OI*) as *OI* = 1− circular variance , where the circular variance was calculated using the *circ_var* function of the CircStat MATLAB toolbox.^[108]^ The circular variance measures the directional spread of fiber angles, ranging from 0 (all fibers aligned) to 1 (completely random orientation). Consequently, the OI ranges from 0 for randomly oriented fibers to 1 for perfectly aligned fibers.

### Spatial transcriptomics

We quantified local RNA expression with spatial context using the GeoMx® RNA assay.^[65,66]^ Described below are the steps in the experimental and analysis pipeline.

#### Sample preparation

We selected 3 sporadic EO CRC and 3 sporadic AO CRC samples, along with matching normal tissues based on age at resection and on inclusion of both epithelial and stromal content with verification by our pathologist (ZC; see patient characteristics for those included in this analysis in Table S1 and S2). We mounted fresh cuts of 5 µm FFPE sections on charged slides, baked, and stained them using the BOND RX/RXm Fully Automated IHC/ISH Stainer from Leica Biosystems, following the manufacturer’s automated slide preparation user manual. We conducted immunofluorescent labeling of distinct cell compartments using the following markers: epithelial cells via pan-cytokeratin (PanCK, Novus NBP2-33200AF532-AF532; 1:40); non-immune stromal cells via Vimentin (Santa Cruz sc373717-AF594, 1:150); leukocytes via CD45 (Nanostring CD45-AF647, clone D9M8I, CST cat number 13917; 5 µg mL^− 1^), cell nuclei via SYTO13 (Nanostring; 1:10). After staining, we immediately scanned all slides. In total, we selected 72 ROI, including: AO Matched Normal (6 for Vimentin), AO Cancer (9 for PanCK, and 17 for Vimentin), EO Matched Normal (7 for Vimentin), EO Cancer (10 for PanCK, and 23 for Vimentin). Section AO2 did not have a matched normal and could not be used due to technical compromise. Overall, we identified 59 ROIs from cancers and 13 from matched normal tissues for extraction and sequencing.

#### Digital Spatial Profiling (DSP)

After hybridization with RNA probes conjugated to barcoded oligonucleotide tags with an ultraviolet (UV) photocleavable linker and staining with fluorescent morphology markers (PanCK, Vimentin, CD45, SYTO13), we loaded the slides onto a GeoMx® (NanoString Technologies, Seattle, WA, USA) digital spatial profiler (DSP) and scanned them. We imaged the labeled tissues and identified specific ROIs based on cellular localization and fluorescent tagging. We defined ROIs as containing at least 30 nuclei, delimited by the morphological markers and the accompanying fluorochrome. We further segmented some ROIs into more areas of illumination (AOI) based on segmentation of morphological markers. Finally, we successively exposed the selected AOIs to UV light to release the barcoded oligos; we aspirated them and dispensed them into a collection plate for library preparation for next-generation sequencing (NGS). We subjected the isolated RNA to bulk RNA sequencing.

#### Library preparation and sequencing

We prepared the NGS libraries following the NanoString GeoMx pipeline. In brief, after the collection was complete, we dried the aspirates in the collection plate at 65°C for 1 hour in a thermal cycler with the lid open and resuspended them in 10 µL of nuclease-free water. We mixed 4 μL of rehydrated aspirates with 2 μL of 5×PCR Master Mix and 4 μL of SeqCode primers, and performed PCR amplification with 18 cycles. We then pooled the PCR products equally and purified twice with 1.2×AMPure XP beads (Beckman Coulter, Brea, CA, USA). We evaluated the final libraries and quantified them using Agilent’s high-sensitivity DNA kit and Invitrogen’s Qubit dsDNA HS assay, respectively. Using the NanoString GeoMx worksheet, we calculated total sequencing reads per DSP collection plate. The libraries were subjected to 2×38 bp paired-end sequencing on an Illumina NovaSeq 6000 system with a 100-cycle S1 kit.

#### Bulk RNA-seq data analysis

We used the Nanostring GeoMx Human Whole Transcriptome Atlas platform to profile 72 ROIs from all slides. This panel profiles the whole transcriptome by targeting 18,677 unique transcripts from human protein-encoding genes plus negative controls from the External RNA Controls Consortium. The panel excludes uninformative high-abundance RNAs such as ribosomal subunits and includes RNA probes designed for Illumina NGS readout with the Seq Code library prep. Digital count conversion files were generated using the GeoMx® NGS Pipeline (Version 2.3.3.10) from FASTQ sequencing files and uploaded into the GeoMx DSP Data Analysis Suite for spatially resolved expression analysis and visualizations. We performed standard quality control using default settings per the manufacturer’s guidelines. All 72 segments passed quality control analysis. Gene filtering retained 14,364 out of 18,677 genes (76.94%) after removing targets with expression at or lower than the selected threshold value at or above specific frequency (5% in our case). We applied Q3 normalization to account for differences in sequencing yield, enabling biological comparison of gene expression across the segments. The normalized count density distributions across individual segments were grossly similar, and there was no observable bias by either pathologic grade or epithelial histology.

We performed differential expression analysis between groups using the *limma* package in R (version 4.4.3) after log2 transformation (with a pseudocount of 1) of the pre-normalized expression data. To maintain the spatial resolution inherent in the GeoMx data while accounting for patient effects, we implemented a ROI-level analysis that treated individual ROIs as observations while incorporating patient IDs as covariates in the statistical model. We fitted a linear model with patient onset type (EO vs. AO) as the main effect of interest and patient ID as a covariate to account for within-patient correlation. This approach preserves spatial heterogeneity information while controlling for patient-specific effects. We used an empirical Bayes method to moderate the standard errors of the estimated log-fold, thus resulting in more stable inference and improved power, especially considering the small numbers of samples. We identified DEGs using thresholds of adjusted *P* value < 0.05 and absolute log2 Fold Change > 1.0. To adjust for multiple testing, we used the Benjamini and Hochberg ^[109]^ method to control false discovery rate (FDR). We performed pathway enrichment analysis using GO gene sets and implemented with the *clusterProfiler* package using the org.Hs.eg.db annotation database for gene identifier mapping. We identified significantly enriched pathways using a *P* value threshold of 0.05 and a q value (FDR) threshold of 0.1. For the epithelial compartment, we performed a comprehensive pathway analysis using the Correlation Adjusted MEan RAnk gene set test (CAMERA).^[110]^ We curated eight distinct mechanotransduction-related gene sets to represent key functional modules of cellular mechanosensing: (1) mechanosensors (ion channels and transmembrane proteins that directly sense mechanical forces); (2) focal adhesion and cell-ECM adhesion proteins; (3) integrins; (4) cytoskeletal components; (5) Rho GTPase signaling proteins; (6) LINC complex components; (7) Hippo pathway components; and (8) mechanosensitive transcription factors (Mechano TFs). For each mechanotransduction gene set, CAMERA calculated a competitive gene set test statistic that evaluated whether genes in the set were more highly differentially expressed between EO and AO samples compared to genes outside the set. Finally, we conducted GSEA^[111]^ using the GSEA (v14) algorithm (https://www.gsea-msigdb.org/gsea/index.jsp). Expression patterns were visualized using the R function *ggplot2*. Correlation analysis between ECM genes was performed using Spearman’s rank correlation. We optimized visualizations for publication quality, with consistent color schemes, with red indicating genes upregulated in EO CRC and blue indicating genes upregulated in AO CRC.

### Cell segmentation and phenotype identification

To achieve a better understanding of the structural and functional relationships between epithelial and stromal cell populations, we used the high-resolution immunofluorescence images acquired for DSP to conduct single-cell segmentation of human CRC tissues. First, we annotated immunofluorescence images using DeepCell Label,^[112]^ a browser-based graphical user interface for editing cell annotations and for training the neural network for single-cell segmentation. Next, we used Mesmer,^[57]^ a deep-learning algorithm to generate automated whole-cell and nuclear boundaries. Briefly, we used nuclear images (SYTO13) to identify the nucleus of each cell, while we used cytoplasmic images (PanCK, Vimentin) to define the area occupied by each cell. Because of the low intensity in the CD45 channel in all images, we decided to exclude the immune marker from this analysis thus avoiding erroneous segmentations. The deep-learning model produces predictions for the centroid and boundary of each nucleus and cell captured in the original image. This method allowed us to segment 2,155,046 cells from 11 patient-derived samples. After segmentation, we extracted single-cell morphological features (area, major and minor axis length, eccentricity, solidity, extent, and orientation) and fluorescence intensity of each marker using the *scikit-image* Python library. The cell aspect ratio was calculated by dividing the length of the major axis by the length of the minor axis. All features were z-score normalized^[113]^ across the dataset to facilitate cross-sample comparison of marker expression (PanCK and Vimentin) across groups and to enable clustering analysis. Finally, we identified distinct cellular phenotypes via unsupervised Louvain clustering,^[58]^ which was implemented using the Python library *Scanpy*.^[114]^ We first constructed a k-nearest neighbor graph based on Euclidean distances computed from the z-normalized feature space, including both morphological features and marker intensities. A resolution of 0.1 was chosen to avoid over-clustering and to identify major cell phenotypes with robust morphological and marker-based separation.^[115]^ Using this approach, we found that all cells within our dataset clustered into four groups.^[116]^ Depending on the average marker expression and the average cell area, the four clusters were assigned to the following cell types: epithelial-large (high PanCK, low Vimentin, high cell area), epithelial-small (high PanCK, low Vimentin, low cell area), stroma (low PanCK, high Vimentin, high cell area), immune (low PanCK, high Vimentin, low cell area). These four phenotypes were validated by superimposing the cell cluster labels on the original immunofluorescence images (Figure S9A-D).

### Immunofluorescence and immunohistochemistry

We stained FFPE blocks sectioned at 4 µm (prepared by the Tissue Management Shared Resources at UT Southwestern) for YAP in human colorectal tissues. Tissue sections were deparaffinized and subjected to antigen retrieval using a tris/borate/EDTA buffer (pH 9.0). The slides were cooled to room temperature and washed twice with PBS. Sections were incubated in 0.3% H_2_O_2_ in PBS for 10 minutes at room temperature and then blocked with 10% serum and permeabilized with 0.3% Triton X-100 in PBS for 15 minutes at room temperature. We stained the slides with anti-active-YAP1 (Abcam, ab205270, 1:2000) at 4°C overnight. The next day, we incubated the slides at room temperature for 1 hour before being washed 3 times with PBS and incubated them with AF594 conjugated secondary antibody (Invitrogen, A32740, 1:1000) for 1 hour at room temperature. After three rounds of PBS wash, we incubated the slides with DAPI (1:2000 in PBS) for 15 minutes at room temperature. Lastly, we mounted the slides using Prolong Gold Antifade (Invitrogen P36931) and kept them in the dark at 4°C and dried overnight. We captured images using an automated widefield epifluorescence slide scanner with LED excitation (Zeiss Axioscan.Z1) at 20× by the Whole Brain Imaging Core at UT Southwestern. We developed a custom ImageJ macro to isolate epithelial regions manually using a brush tool to exclude non-epithelial cells from subsequent analyses. We quantified the fluorescent intensity of nuclear active YAP as discussed in the section “*Quantitative confocal microscopy*”.

We stained for Ki67 in human colorectal tissues using FFPE blocks sectioned at 4 µm obtained from the Tissue Management Shared Resource core at UT Southwestern following standard protocols.^[117]^ Tissue sections were deparaffinized and subjected to antigen retrieval using a sodium citrate buffer containing 0.05% Triton X-100 (pH 6.0). We stained the slides using an antibody for Ki67 (Proteintech 27309-1-AP, 1:5000) and imaged using a Whole Slide Scanner (NanoZoomer by Hamamatsu) at 20× magnification. We quantified Ki67+ cells using an automatic cell detection tool in QuPath-v0.5.0,^[118]^ an open source software for digital pathology image analysis. Briefly, IHC image files were opened in QuPath as Brightfield H-DAB images. Epithelial cancer cell regions were selected using the brush tool. QuPath’s Positive Cell Detection tool was used to quantify the number of DAB positive nuclei (Ki67+) on cancer regions using the hematoxylin OD image as the detection image.

### Cell culture experiments

We prepared polyacrylamide (PA) gels by mixing solutions of 40% acrylamide and 2% bis-acrylamide (Bio-Rad, Hercules, CA) using established protocols.^[119,120]^ Briefly, we treated the bottom of 12-well glass bottom plates (Cellvis P12-1.5H-N) with 1 mL of bind silane solution (80 µL glacial acetic acid, 50 µL 3-(trimethoxysilyl)propyl methacrylate, and 200 mL deionized water) for 1 hour at room temperature. We then rinsed the glass bottom plates thoroughly with distilled water and dried with an air gun. Top coverslips (18 mm in diameter) were passivated by treating them with 0.1 M NaOH for 1 hour at room temperature, rinsed 3 times with deionized water, and then dried with compressed N_2_. Soft PA gels consisted of 75 µL acrylamide and 24 µL bis-acrylamide, while stiff gels were prepared by mixing 188 µL acrylamide and 19 µL bis-acrylamide. In both cases, we added 0.5 µL TEMED and 100 µL of 5% ammonium persulfate solution, and sterile deionized water to achieve a 1 mL solution. We placed a 200 µL droplet in each well and covered them with a passivated coverslip to spread the liquid uniformly across the bottom of the well. We allowed the PA to polymerize for 30 minutes under vacuum and then removed the top coverslip by using fine forceps. We assessed the mechanical properties of the resulting PA gels using the unconfined compression protocol described above (Figure S17). We treated polymerized PA gels with 440 μL of a 1:50 (v/v) mixture of Sulfosuccinimidyl 6-(4’-azido-2’-nitrophenylamino)hexanoate (Sulfo-SANPAH) (ProteoChem) and 50 mM HEPES solution under a UV light for 10 min. We then incubated PA gels in either 0.2 mg mL^−1^ collagen I (Corning, 354249) or 1.5% growth factor– reduced Matrigel (Corning, 354230) at 37°C for 30 minutes. We cultured the colon cancer lines HT29 (ATCC, HTB-38, 44-year-old female) and SW480 (ATCC, CCL-228, 50-year-old male) in DMEM (Corning, 10013CV), supplemented with 10% fetal bovine serum (ATCC, 302020) and 1% penicillin/streptomycin (ATCC, 302300). We authenticated both cell lines by short-tandem repeat profiling using a commercial kit (ATCC, 135-XV). Cell cultures were maintained in humidified incubators at 37°C and 5% CO_2_ in T75 flasks and were monitored regularly under a phase contrast microscope. We routinely tested the cells for mycoplasma contamination using the MycoAlert kit (Lonza, LT07-218) to ensure that cultures remained contamination-free. At the time of passage, we seeded HT29 cells at a density of 50 × 10^3^ cells cm^−2^ and SW480 cells at a density of 40 × 10^3^ cells cm^−2^ due to their different rates of proliferation. We allowed the cells to grow on PA gels for one day and then cultured in serum-free DMEM. After 72 hours of culture on PA gels of varying stiffnesses, we fixed the cells using 4% paraformaldehyde (PFA) in PBS for 15 minutes, permeabilized them using 0.2% Triton X-100 for 15 minutes and blocked using 2.5% normal horse serum for 15 minutes. We washed the samples 3 times with PBS and incubated them with a primary anti-active-YAP1 (Abcam, ab205270, 1:2000) antibody overnight on a shaker at 4°C. After washing, we incubated the samples in Alexa Fluor 594 conjugated Goat anti-Rabbit secondary antibody (Invitrogen, A32740, 1:1000) for 1 hour at room temperature and counterstained them with Alexa Fluor 647 phalloidin (Invitrogen, A22287, 1:1000) and DAPI (Invitrogen, D1306, 1:1000). We assessed cell proliferation using the Click-iT EdU Cell Proliferation Kit (Thermo Fisher, C10637). Briefly, we prepared a 2x working solution of EdU (Component A) in serum-free DMEM and added it by replacing 500 μL of medium in each well. After incubation, we fixed the cells with 4% PFA for 15 minutes, permeabilized with 0.2% Triton X-100 for 15 minutes and washed thoroughly with PBS. We prepared the Click-iT reaction cocktail containing Alexa Fluor 488 azide according to manufacturer’s protocol, and added 200 μL to each gel surface for 30 minutes at room temperature, and shielded the wells from light to enable click chemistry detection of incorporated EdU. For visualization, we then counterstained the samples with DAPI (Invitrogen, D1306, 1:1000). We inhibited YAP activity by culturing HT29 and SW480 cells in serum-free DMEM supplemented with verteporfin (Sigma, SML0534) at concentrations of 1, 2, 4 or 8 μg mL^−1^ dissolved in 0.5% DMSO. Cell viability was assessed using the LIVE/DEAD Viability/Cytotoxicity Kit (Invitrogen, L3224). After 48 hours of incubation with verteporfin, the cell culture medium was removed and cells were washed with 1 mL of sterile PBS. Next, 400 µL of a solution containing 2 µM calcein AM and 4 µM ethidium homodimer-1 (EthD-1) in sterile PBS was added to cells for 30 minutes, after which cells were immediately imaged. Phalloidin and EdU staining on verteporfin-treated cells were carried out as described above.

### Quantitative confocal microscopy

We imaged the fixed and stained samples using a Nikon AX confocal microscope and a line series acquisition mode to minimize crosstalk between the 5 fluorescent channels: 405nm excitation / 420-470nm emission (DAPI), 488nm excitation / 500-530nm emission (EdU), 561 excitation / 570-616nm emission (Active YAP), 640nm excitation / 660-750nm emission (Phalloidin), and transmitted detector for differential interference contrast (DIC) imaging. We acquired image stacks using the following settings: 2x line averaging, 1024 × 1024 pixels at a resolution of 0.576 μm pixel^−1^ in x-y, 11 z-slices with 11μm step size covering 121μm total depth. Once acquired, we transformed each stack into a maximum intensity projection image and analyzed the individual channels using custom ImageJ macros. For cell proliferation analysis, we used maximum intensity projections with intensity thresholding (DAPI: 50-255, EdU: 100-255) followed by watershed segmentation and particle analysis (size: 55-5,000 μm²) to identify total nuclei and proliferating cells. Cell proliferation was quantified as the percentage of EdU-positive nuclei relative to total DAPI-positive nuclei. To quantify active YAP intensity in cell nuclei, we identified nuclear regions in epithelial cells by intensity thresholding of DAPI (75-255) and measured active YAP fluorescence intensity within these nuclear compartments. We quantified cell morphology by outlining the phalloidin-positive cytoplasmic region of individual cells using manual tracing with a polygon tool. For each cell, the aspect ratio was calculated by dividing the length of the major axis by the length of the minor axis. We analyzed a minimum of 5 cells and a maximum of 70 cells per ROI, depending on cell density. For viability analysis, we imaged cells using the following fluorescent channels: 488 nm excitation / 500-530 nm emission (calcein AM, live cells) and 561 nm excitation / 570-616 nm emission (EthD-1, dead cells). We quantified cell viability by applying intensity thresholding to maximum intensity projections (live cells: 50-255, dead cells: 50-255), followed by watershed segmentation and particle analysis to identify individual cells. Particle analysis parameters included size: 50-800 μm² and circularity: 0.5-1.0. We calculated percent cell viability as the ratio of live cells to total cells, expressed as a percentage.

### Organoid experiments

We established and characterized human CRC patient-derived epithelial organoids as described previously.^[80–82]^ We selected three sporadic EO CRC and three sporadic AO CRC samples based on age at resection, ability to be expanded in vitro, and tumor stage diversity to enhance generalizability of our findings (see patient characteristics for those included in this analysis in Table S1 and S2). Briefly, we isolated the cells from the surgically resected colorectal cancer specimens and plated them in Matrigel (Corning, 354230) in 24-well Nunclon Delta plates (ThermoFisher, 142475). Following Matrigel polymerization, we overlaid the organoids with 500 µL of “complete” medium composed of advanced DMEM/F12 (Gibco, 12634) supplemented with 20% FBS, L-WRN medium, 100x GlutaMAX^TM^ (Gibco, 35050), SB431542 (Tocris, 1614), Y27632 (Tocris, 1254) and 1% penicillin/streptomycin (Gibco, 15140). We maintained the organoid cultures in humidified incubators at 37°C with 5% CO_2_ and monitored them regularly under a phase contrast microscope. We changed the culture medium every two days. Authentication of all organoid models occurred by short-tandem repeat profiling at the UT Southwestern Genomics Core via the PowerPlex Fusion System, and we regularly tested for mycoplasma contamination via the MycoAlert kit (Lonza, LT07-710). We encapsulated patient-derived organoids in 25 µL Tyramine-Substituted Hyaluronate (TS-HA; synthesized at the Cleveland Clinic).^[82]^ Two TS-HA concentrations (2.5 mg mL^−1^ and 8 mg mL^−1^) were used to make hydrogel beads. Briefly, we washed 10^3^ organoids with PBS and resuspended them in 24 µL of activated TS-HA at room temperature. Addition of Horseradish Peroxidase (Sigma-Aldrich; cat# P8250-5KU; at 1:500 ratio) activated the TS-HA solution. We used sterile 12-well silicon isolators (Sigma-Aldrich; cat# GBL665506) of 25 µL each as molds for individual beads. TS-HA crosslink reaction with 1 µL of 0.03% hydrogen peroxide was achieved individually in-well. We washed the hydrogel beads with organoid media and incubated them at 37°C in a 5% CO_2_ incubator for 20 to 30 minutes to remove residual chemicals, and then changed the medium. We used the TS-HA beads for experiments within 48 hours. We used a CellTiter-Glo® 3D Cell Viability assay (Promega, G9681) to measure viability of encapsulated organoids in TS-HA at both 2.5 mg mL^−1^ and 8 mg mL^−1^ from both AO and EO CRC patients. Briefly, we incubated organoid-containing hydrogel beads overnight in multi-well clear-bottom opaque plates and added an equal volume of CellTiter-Glo® 3D Reagent to the volume of cell culture medium present in each well. After vigorous shaking, we incubated the plates at room temperature for an additional 25 minutes to stabilize the luminescent signal. We recorded luminescence using a Molecular Devices Spectramax (Model ID3) plate reader. To account for patient-to-patient variability, we normalized the raw luminescence readings acquired from a given patient organoids with respect to the lowest reading on soft (2.5 mg mL^−1^) TS-HA hydrogels. This approach allowed us to quantify relative changes in cell viability.

### Short-tandem repeat fingerprinting

We conducted short-tandem repeat analysis to confirm that patient-derived organoids cultured in vitro preserved the same genomic identity as the corresponding FFPE tissues (Table S3). Once established, we compared the organoid DNA isolate (DNeasy Blood & Tissue Kit, Qiagen 69506) to genomic DNA isolated from the companion FFPE sections (GeneRead DNA FFPE Kit Qiagen 180134). The University of Texas Southwestern McDermott Center Sequencing Core (Department Eugene McDermott Center for Human Growth and Development) confirmed the unique origin of each primary epithelial organoid prior to use in the 3D culture assay outlined above.

### Statistical analysis

We present experimental data as mean ± standard deviation (SD), as box plots, or as violin plots with individual data points showing independent measurements. We present raw biomechanical measurements as mean ± standard error of the mean (SEM) error envelopes to visualize average trends across experimental groups. No statistical method was used to predetermine sample size, and investigators were not blinded during experiments or outcome assessment. We assessed data normality using the Shapiro-Wilk test and density plots, which revealed non-normal distribution patterns. The Scheirer-Ray-Hare test was used as a non-parametric alternative to two-way ANOVA to assess differences between: (i) cancer and matched normal samples and between AO and EO tissues; (ii) cell proliferation based on ligand adhesion (Collagen vs. Matrigel), pharmacological treatment (No Treatment vs. Verteporfin), and PA stiffness (Soft, Stiff, and Glass); and (iii) organoid viability between AO and EO organoids cultured in TS-HA hydrogels of tunable stiffness (Soft vs. Stiff). Additionally, we used a Kruskal-Wallis test to determine significant differences in cell aspect ratio, in the fraction of EdU-positive nuclei, and active YAP fluorescence intensity across three PA stiffnesses (Soft, Stiff, and Glass). For post-hoc multiple comparisons, Dunn’s test with Bonferroni correction identified significant differences between specific groups. Results were considered statistically significant at *P* < 0.05. All analyses were performed using R (version 4.4.3).

## ACKNOWLEDGMENTS

We thank Elena Seymour and Reyan Ghanim for conducting preliminary experiments on polyacrylamide substrates and Nathaniel Roberts for assisting with cell morphology quantifications. We also thank Manal Ali for assistance with the collection of both fresh and fixed primary tissues, and Dr. Dan Zhao for initial in vitro experiments. We gratefully acknowledge Dr. Huocong Huang for helpful discussions and guidance on the bioinformatic analysis of spatial RNA sequencing.

## Funding

University of Texas at Dallas Office of Research and Innovation through the CoBRA program (JF, EHH).

University of Texas at Dallas, Bioengineering Department Startup funds (JF). University of Texas at Dallas, Bioengineering Researcher Award (NCH).

University of Texas Southwestern, Department of Surgery Startup funds (EHH, MHB).

University of Texas Southwestern Early Onset Colorectal Cancer Initiative (Simmons Comprehensive Cancer Center) (EHH).

National Institutes of Health grant R01 CA234307 (EHH). National Institutes of Health grant U01 CA214300 (EHH).

Burroughs-Wellcome Trust (MHB).

American Society of Colon and Rectal Surgeons Resident Research Initiation Grant (MHB).

The work reported in this publication was supported by the National Cancer Institute of the National Institutes of Health under award number P30 CA142543 (CML).

University of Texas Whole Brain Microscopy Facility RRID:SCR_017949, and Axioscan 7 Award number 1S10ODO032267-01 to Denise Ramirez (Core Director).

## Author Contributions

Conceptualization: JF, EHH

Methodology: JF, RCF, IR, ARJ, SR, VDV, CML

Investigation: JF, NCH, MHB, VVN, AK-S, HA, AK, AF, GC, CZ, MY

Pathology: ZC

Visualization: JF, NCH, VVN, MHB, CZ, MY

Supervision: JF, EHH

Writing – original draft: NCH, MHB, VVN, JF, EHH

Writing – review & editing: JF, EHH

## Data and materials availability

All data in the main text or in the supplementary materials are available upon request. Data and codes supporting the findings of this study are available from the corresponding authors upon reasonable request. Primary human tissues have been used throughout this manuscript and only histological blocks are available via deidentified Material Transfer Agreement (MTA). The GeoMx bulk transcriptome data on the human colon cancer and matched normal tissues, with PanCK, Vimentin, and CD45 stained and retrieved clusters have been deposited into the Gene Expression Omnibus and are available via GSE281413.

