## Supplementary Material for "Biomechanical Phenotyping Reveals Unique Mechanobiological Signatures of Early-Onset Colorectal Cancer"

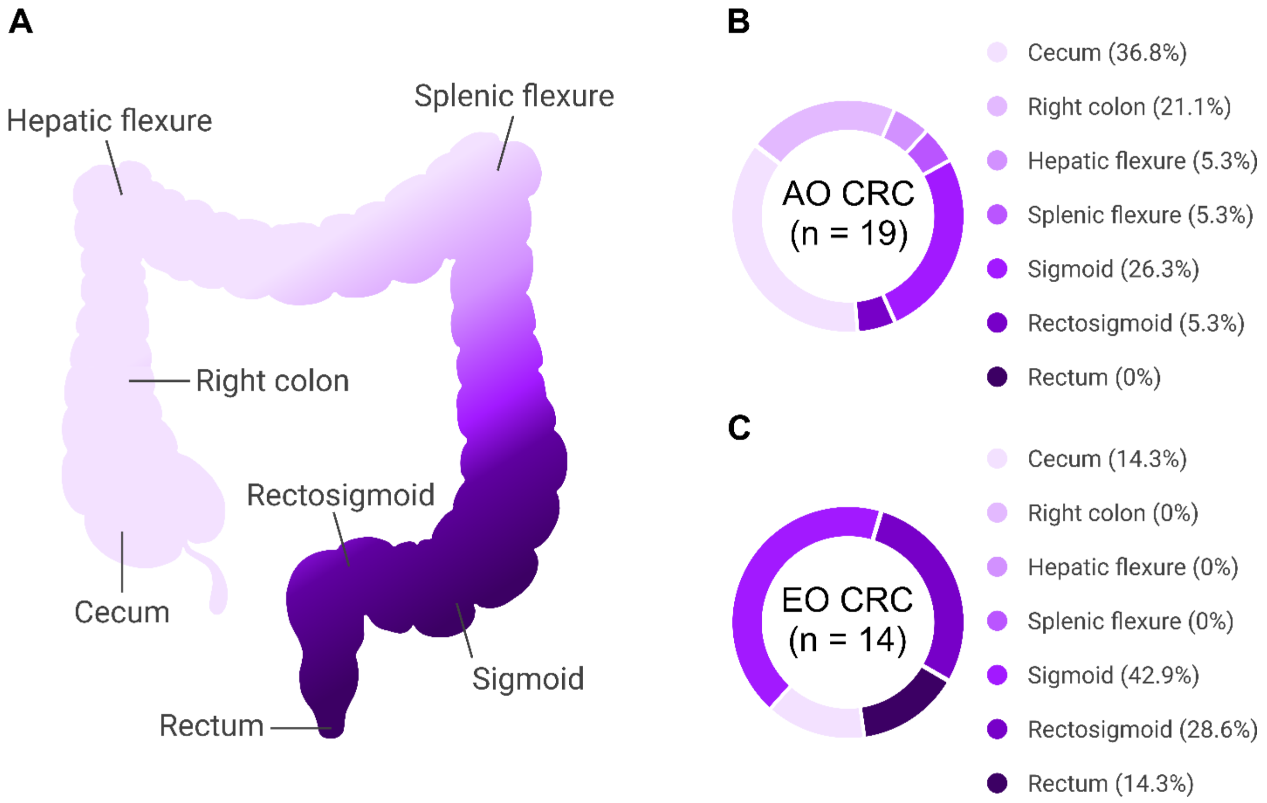


**Figure S1. Sporadic EO CRC favors distal anatomic locations.** (**A**) Schema of the human colon with key anatomical locations labeled. The anatomy of the colon is visually indicated using shades of purple, with light purple indicating the most proximal location (cecum) and dark purple indicating the most distal location (rectum). Pie charts of the tumor locations for the (**B**) AO CRC and (**C**) EO CRC patient cohorts. Overall samples sizes are indicated at the center of each pie chart and the percent tumor occurrence for each anatomical location is listed to the right of the pie chart. Detailed information on the AO CRC and EO CRC patient cohorts can be found, respectively, in Table S1 and S2.

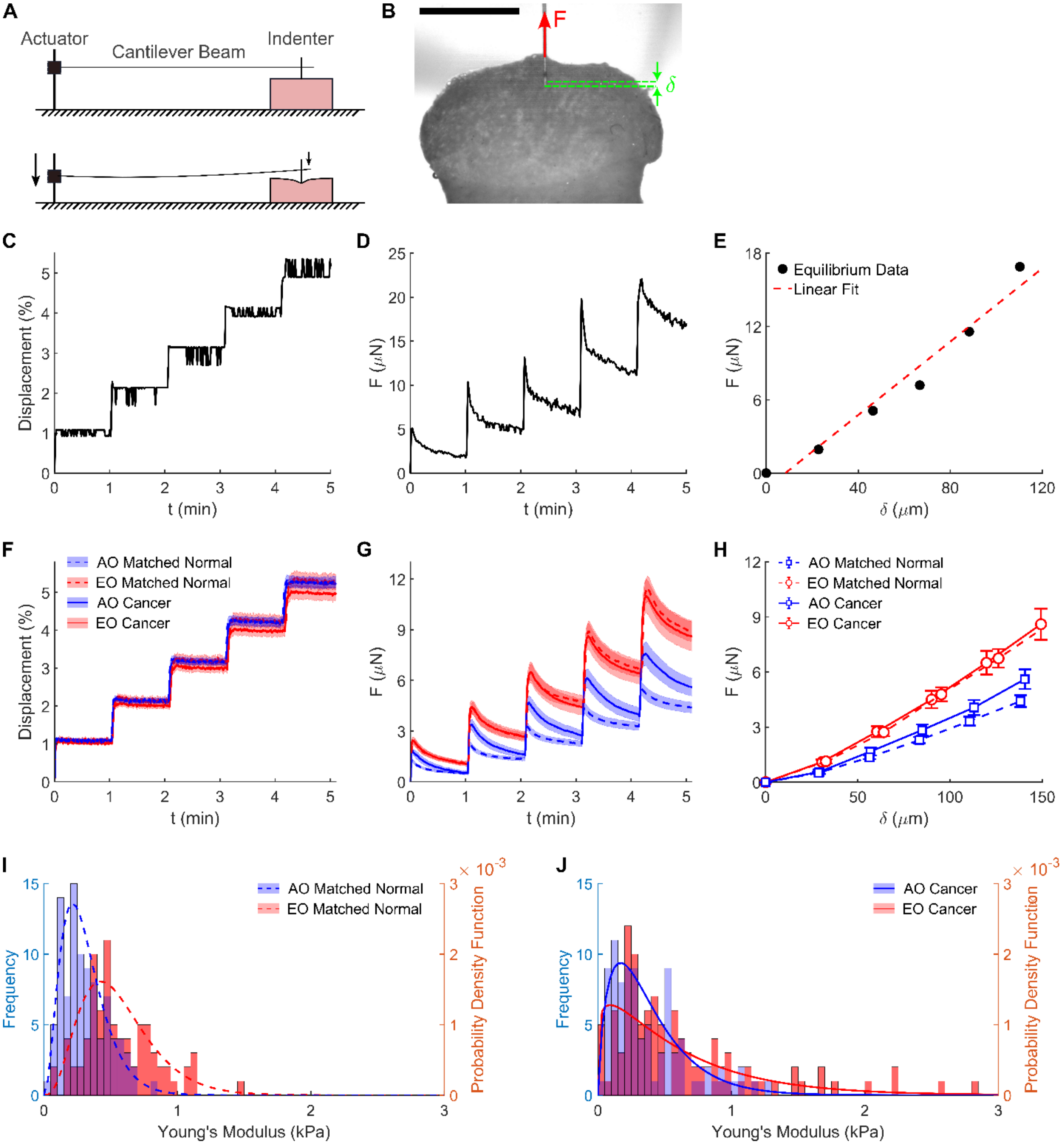


**Figure S2. Analysis of local (mesoscale) tissue mechanics via indentation testing.** (**A**) Schematic representation of the approach used to assess the local tissue stiffness by pairing a cantilever beam with an indenter (top) and using an actuator to locally deform the tissue (bottom). (**B**) The imposed indentation (δ) and the resulting force (F) measured from the cantilever deflection are schematically superimposed on a camera caption of a representative colorectal tissue specimen.

Scale bar, 2 mm. (**C**) Representative indentation data show that tissue displacement was imposed in 5 incremental steps, each with a magnitude equal to 1% of the original tissue height of and interspersed with 1-minute hold phases. (**D**) Representative force evolution in response to the 5 stress-relaxation steps, and (**E**) the associated steady-state force values graphed versus the tissue indentation and fitted with a linear best fit line, which determines the local Young's modulus. Average (**F**) indentation time-course, (**G**) force time-course, and (**H**) steady-state force vs. indentation for the four groups (n = 94 for AO Matched Normal, n = 89 for AO Cancer, n = 80 for EO Matched Normal, n = 102 for EO Cancer). Data are presented as mean ± SEM. Local stiffness distributions for (**I**) matched normal and (**J**) cancer tissues comparing the raw data histograms and the fitted Gamma distribution functions between AO and EO samples. In both matched normal and cancer tissues the local stiffness distributions are broader and right-ward shifted for EO tissues, which indicates an increase in local tissue stiffness.

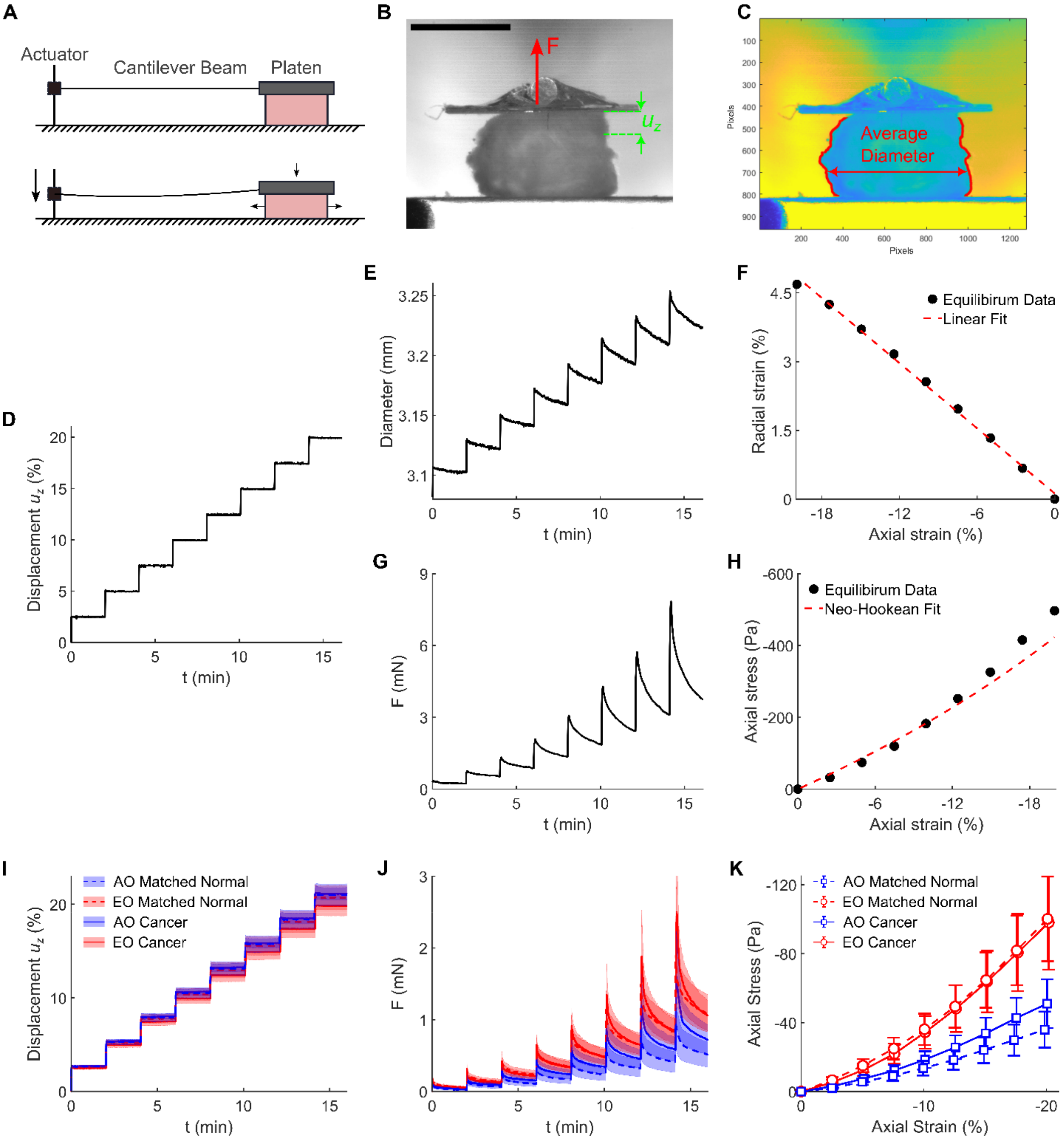


**Figure S3. Analysis of global (macroscale) tissue mechanics via unconfined compression testing.** (**A**) Schematic representation of the approach used to assess the global tissue stiffness by pairing a cantilever beam with a 6 mm × 6 mm platen (top) and using an actuator to compress the tissue (bottom). (**B**) The imposed displacement ($u_z$) and the resulting force (F) measured from the cantilever deflection are schematically superimposed on a camera caption of a representative colorectal tissue specimen. Scale bar, 2 mm. (**C**) Automated quantification of the average diameter of cylindrical specimens for the purpose of determining radial expansion upon axial compression (also known as Poisson's effect). (**D**) Representative unconfined compression data show that tissue compression was performed in 8 incremental steps, each with a magnitude equal to 2.5% the original tissue height interspersed with 2-minute hold phases. (**E**) Representative diameter evolution in response to the 8 stress-relaxation steps, and (**F**) the associated radial strains at equilibrium plotted against the axial strain at equilibrium for the purpose of determining the tissue's Poisson's ratio (dashed red line indicates the linear fit to data). (**G**) Representative force evolution in response to the 8 stress-relaxation steps, and (**H**) the associated steady-state axial stress graphed versus the axial strain and fitted using a nonlinear Neo-Hookean model (cf. Equation 2) to determine the global Young's modulus. Average (**I**) percent axial displacement time-course, (**J**) force time-course, and (**K**) steady-state axial stress vs. axial strain curves for the four groups (n = 21 for AO Matched Normal, n = 21 for AO Cancer, n = 19 for EO Matched Normal, n = 21 for EO Cancer). Data are presented as mean ± SEM.

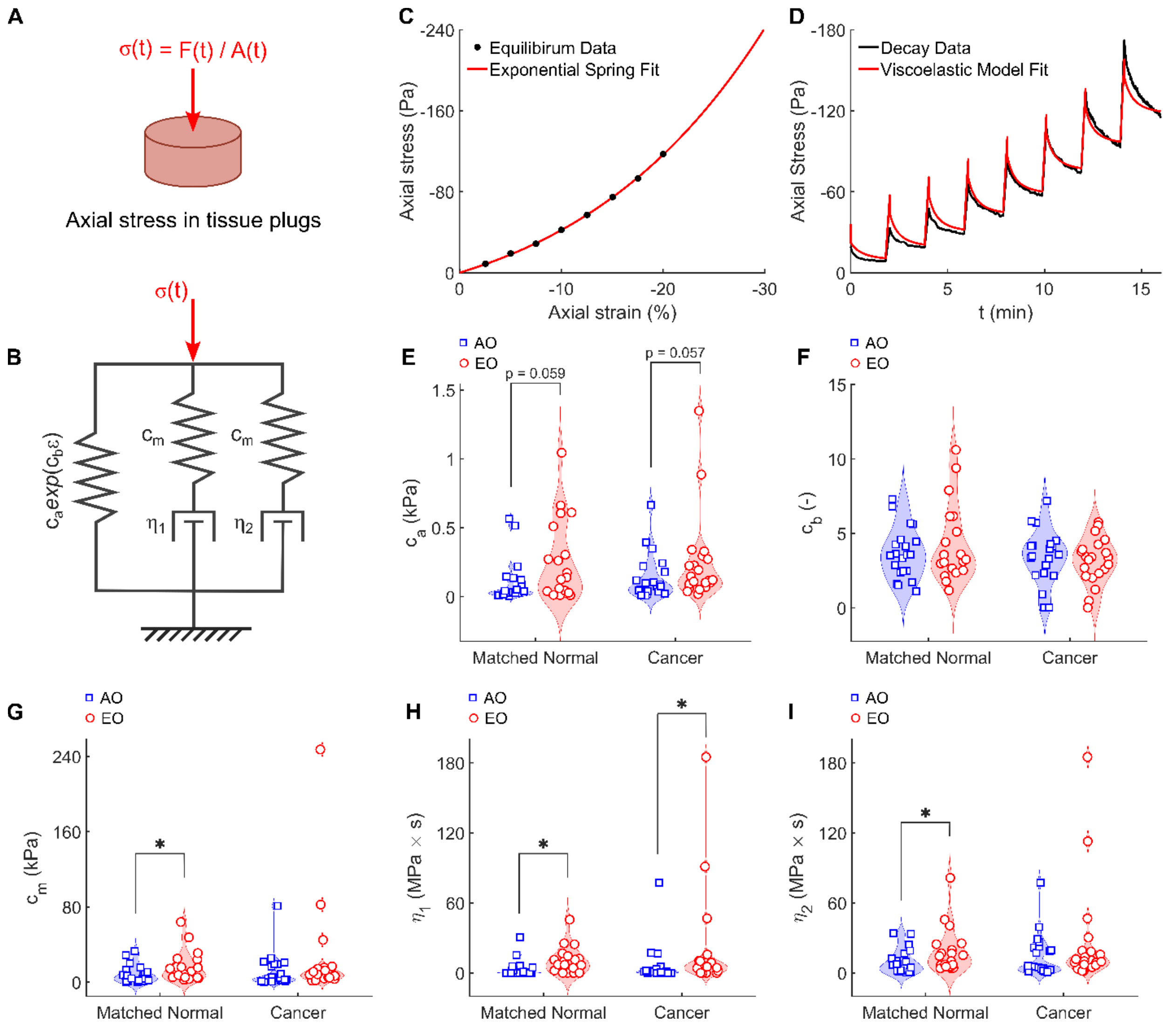


**Figure S4. Viscoelastic modeling of unconfined compression data.** (**A**) The axial stress in cylindrical tissue plugs was estimated by using the cross-sectional area evolution A(t) (estimated from the diameter evolution, cf. Figure S3E) to normalize the measured force evolution F(t) (Figure S3G). The resulting axial stress time-course was fitted using a modified Maxwell-Wiechert viscoelastic model shown schematically in (**B**). Representative (**C**) steady-state axial stress vs. axial strain curve and (**D**) force time-course data (black dots and lines) with superimposed viscoelastic model fits (orange lines). Bar plots show differences in the estimated model parameters between the four groups: (**E**) nonlinear spring stiffness $c_a$, (**F**) nonlinear spring exponent $c_b$, (**G**) Maxwell spring stiffness $c_m$, (**H**) Maxwell fast dashpot viscosity $\eta_1$, (**I**) Maxwell slow dashpot viscosity $\eta_2$ for the four groups (n = 21 for AO Matched Normal, n = 21 for AO Cancer, n = 19 for EO Matched Normal, n = 21 for EO Cancer). Data are presented as violin plots with data points indicating individual measurements. Statistical significance was assessed using a Scheirer-Ray-Hare test with Dunn's post-hoc test and Bonferroni correction. * indicates significant differences ($P < 0.05$) between cancer and matched normal tissues within onset groups (AO, EO) and between onset groups within tissue types (matched normal, cancer).

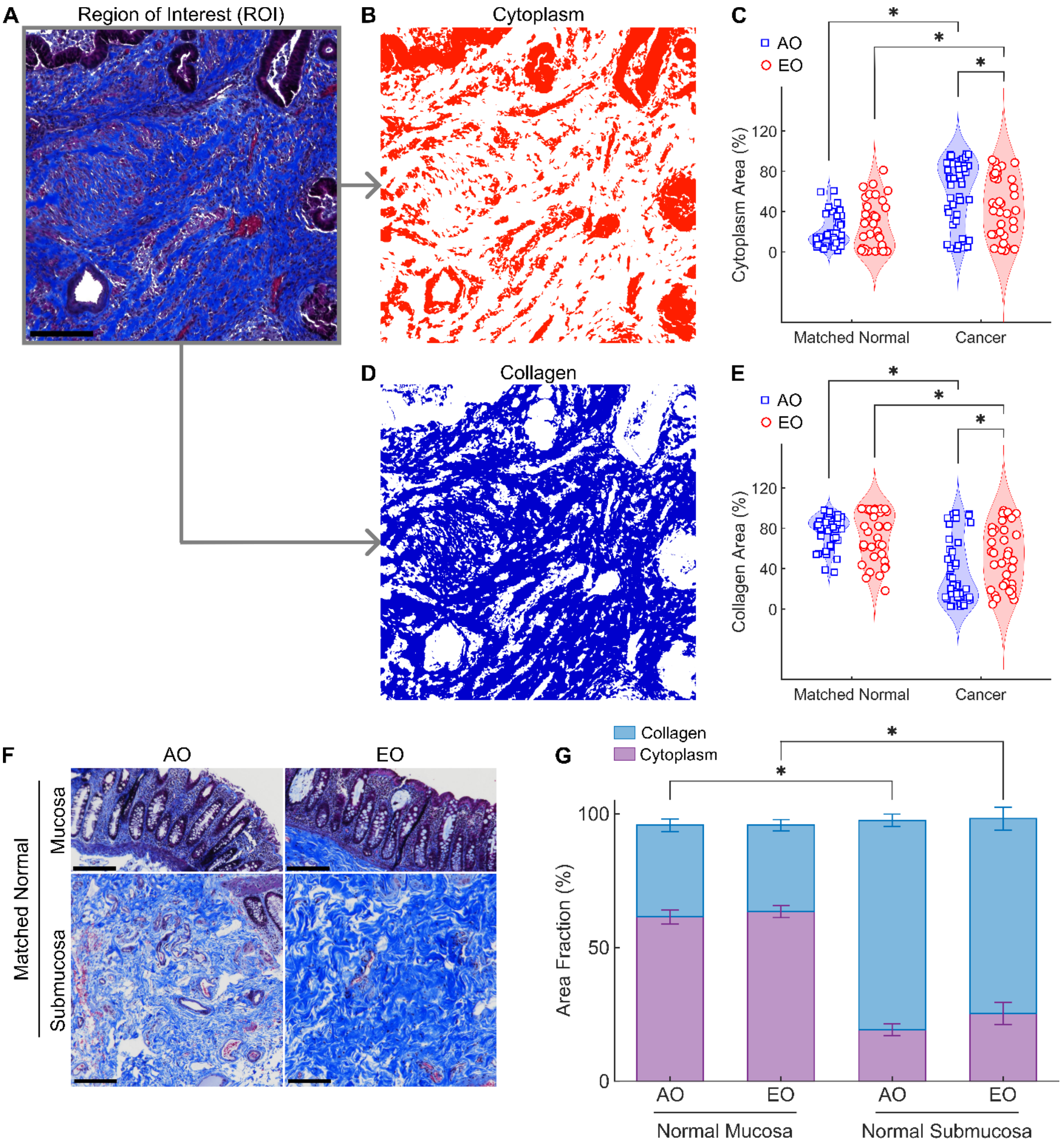


**Figure S5. Masson's Trichrome (MTC) image analysis and results.** (**A**) Representative region of interest (ROI) obtained from an FFPE block sectioned at 4 μm and stained using MTC. (**B**) Binary mask displaying pixels identified as cytoplasm (red-purple staining) and (**C**) associated violin plot showing differences in the cytoplasm area fraction between the four groups. (**D**) Binary mask displaying pixels identified as collagen (blue staining) and (**E**) associated violin plot showing differences in the collagen area fraction between the four groups. n = 45 for AO Matched Normal, n = 45 for AO Cancer, n = 35 for EO Matched Normal, n = 35 for EO Cancer. The comparison of cytoplasm/collagen area fractions was conducted between the normal submucosa and the cancer stroma to focus on collagen-rich areas in both normal and cancer tissues. To support this comparison, we compared (**F**) representative MTC images and (**G**) the associated quantifications of cytoplasmic and collagen area fractions to show that the normal mucosa is rich in cytoplasm while the normal submucosa is rich in collagen. Scale bars, 200 µm. Data comparing normal mucosa and submucosa are presented as mean ± SEM. Data comparing normal vs. cancer samples are presented as violin plots with data points indicating individual measurements. Statistical significance was assessed using a Scheirer-Ray-Hare test with Dunn's post-hoc test and Bonferroni correction. * indicates significant differences ($P < 0.05$) between cancer and matched normal tissues within onset groups (AO, EO) and between onset groups within tissue types (matched normal, cancer).

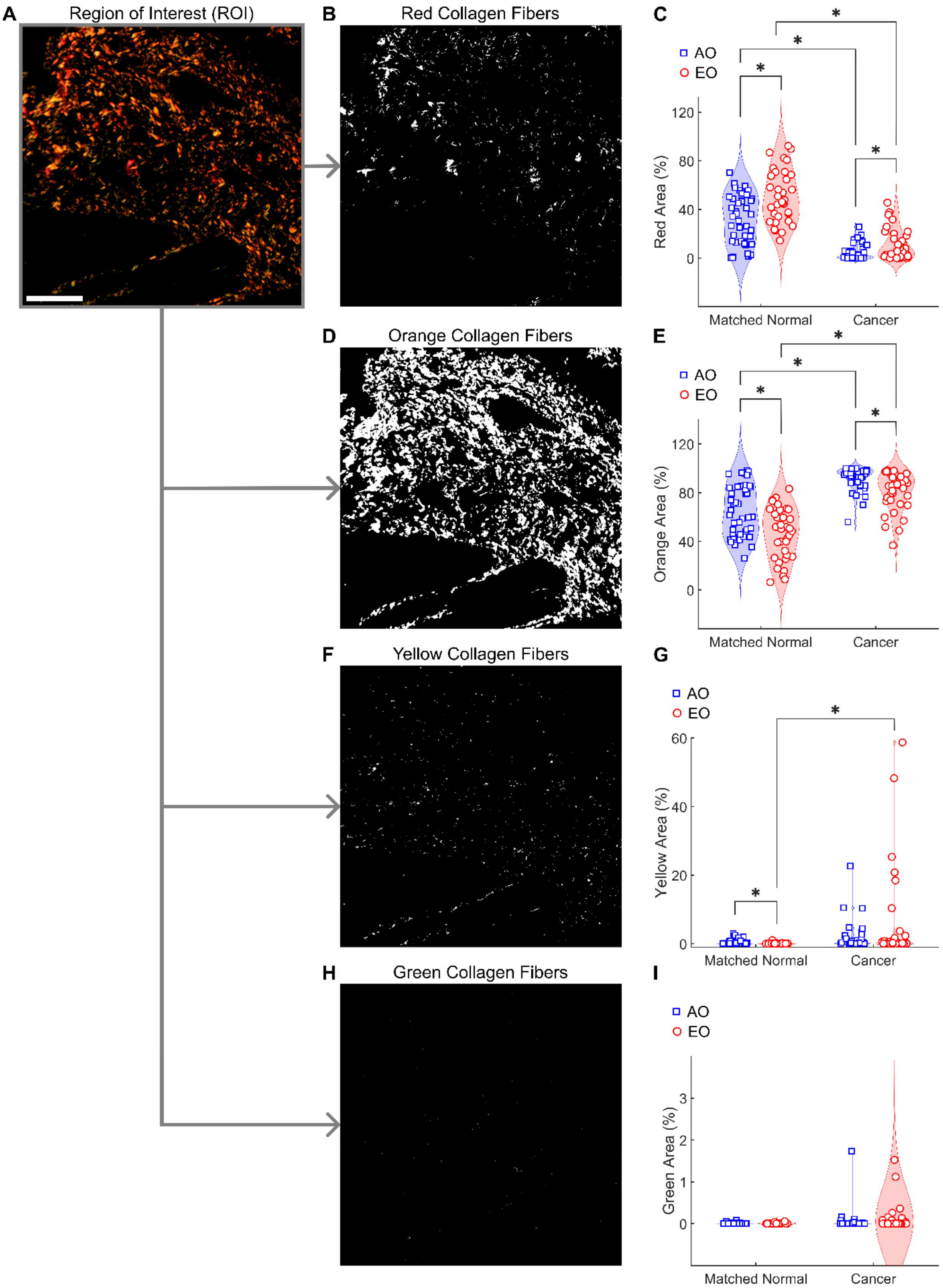
A
Region of Interest (ROI)
B
Red Collagen Fibers
C
AO
EO
Red Area (%)
Matched Normal
Cancer
D
Orange Collagen Fibers
E
Orange Area (%)
F
Yellow Collagen Fibers
G
Yellow Area (%)
H
Green Collagen Fibers
I
Green Area (%)

**Figure S6. Picrosirius Red (PSR) image analysis and results.** (**A**) Representative Region of Interest (ROI) obtained from an FFPE block sectioned at 4 µm and stained using PSR. Images were acquired under polarized light illumination where collagen fibers appear colored in red, orange, yellow, and green based on their birefringence. Scale bar, 200 µm. A colorimetric analysis was used to quantify the distribution of collagen fibers from thick (red) to thin (green). (**B**) Binary mask displaying pixels identified as red collagen fibers and (**C**) associated violin plot showing differences in the red collagen area fraction between the four groups. (**D**) Binary mask displaying pixels identified as orange collagen fibers and (**E**) associated violin plot showing differences in the orange collagen area fraction between the four groups. (**F**) Binary mask displaying pixels identified as yellow collagen fibers and (**G**) associated violin plot showing differences in the yellow collagen area fraction between the four groups. (**H**) Binary mask displaying pixels identified as green collagen fibers and (**I**) associated violin plot showing differences in the green collagen area fraction between the four groups. n = 45 for AO Matched Normal, n = 45 for AO Cancer, n = 35 for EO Matched Normal, n = 35 for EO Cancer. Data are presented as violin plots with data points indicating individual measurements. Statistical significance was assessed using a Scheirer-Ray-Hare test with Dunn's post-hoc test and Bonferroni correction. * indicates significant differences ($P < 0.05$) between cancer and matched normal tissues within onset groups (AO, EO) and between onset groups within tissue types (matched normal, cancer).

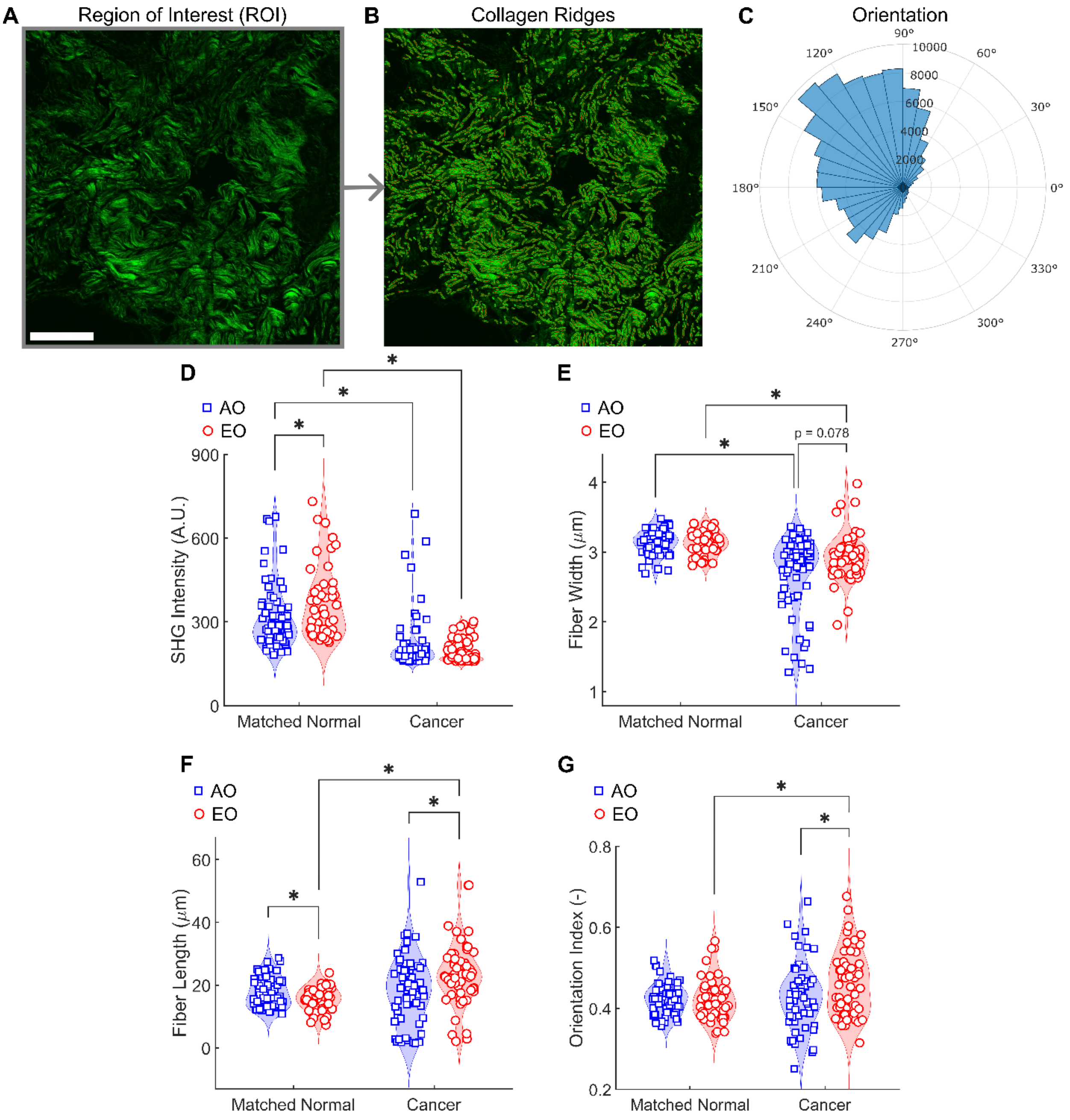


**Figure S7. Second Harmonic Generation (SHG) image analysis and results. (A)** Representative Region of Interest (ROI) obtained from an FFPE block sectioned at 4 µm and stained using H&E. Images were acquired under multiphoton excitation. Scale bar, 200 µm. (**B**) Collagen fibers identified as ridges (using the ImageJ plugin Ridge Detection) are shown superimposed over the raw image and were used to quantify the local collagen fiber architecture.

(**C**) Polar histogram showing the orientation of the ridges for the representative image was used to calculate an Orientation Index (OI, see Methods) to quantify collagen alignment. Violin plots show differences in (**D**) overall SHG signal intensity, as well as in collagen fiber (**E**) width, (**F**) length, and (**G**) OI between the four groups (n = 65 for AO Matched Normal, n = 64 for AO Cancer, n = 47 for EO Matched Normal, n = 55 for EO Cancer). Data are presented as violin plots with data points indicating individual measurements. Statistical significance was assessed using a Scheirer-Ray-Hare test with Dunn's post-hoc test and Bonferroni correction. * indicates significant differences ($P < 0.05$) between cancer and matched normal tissues within onset groups (AO, EO) and between onset groups within tissue types (matched normal, cancer).

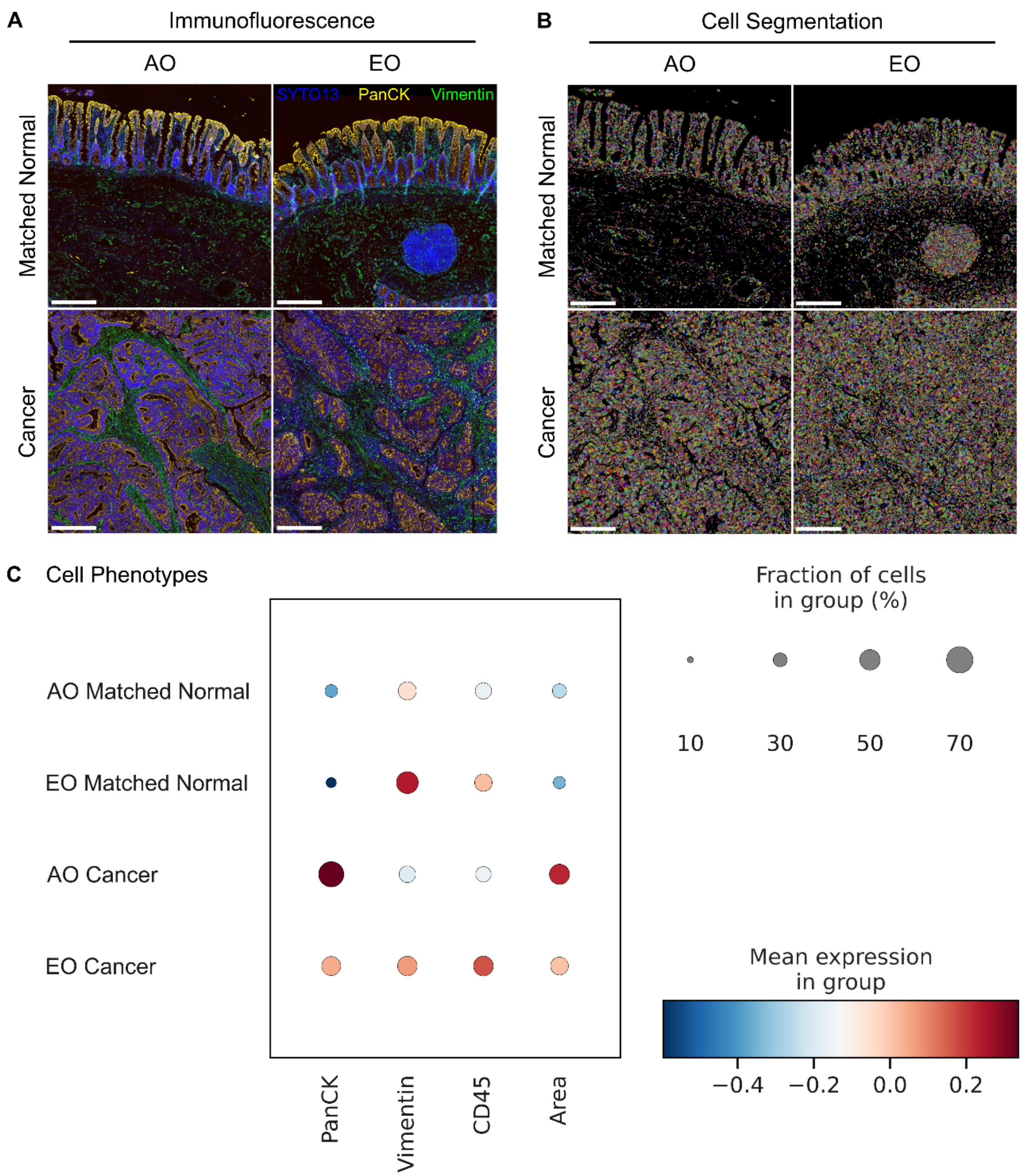


**Figure S8. Cellular features separated via single-cell segmentation.** (**A**) Immunofluorescence images showing SYTO13 in blue, PanCK in yellow, and Vimentin in green. (**B**) Segmented images showing individually segmented cells labeled using different colors. Scale bars, 200 μm. (**C**) The dot plot shows differences in cellular features between the four groups. The dot color indicates the z-normalized expression of each cell feature within each group, while the dot size indicates the fraction of cells within the group.

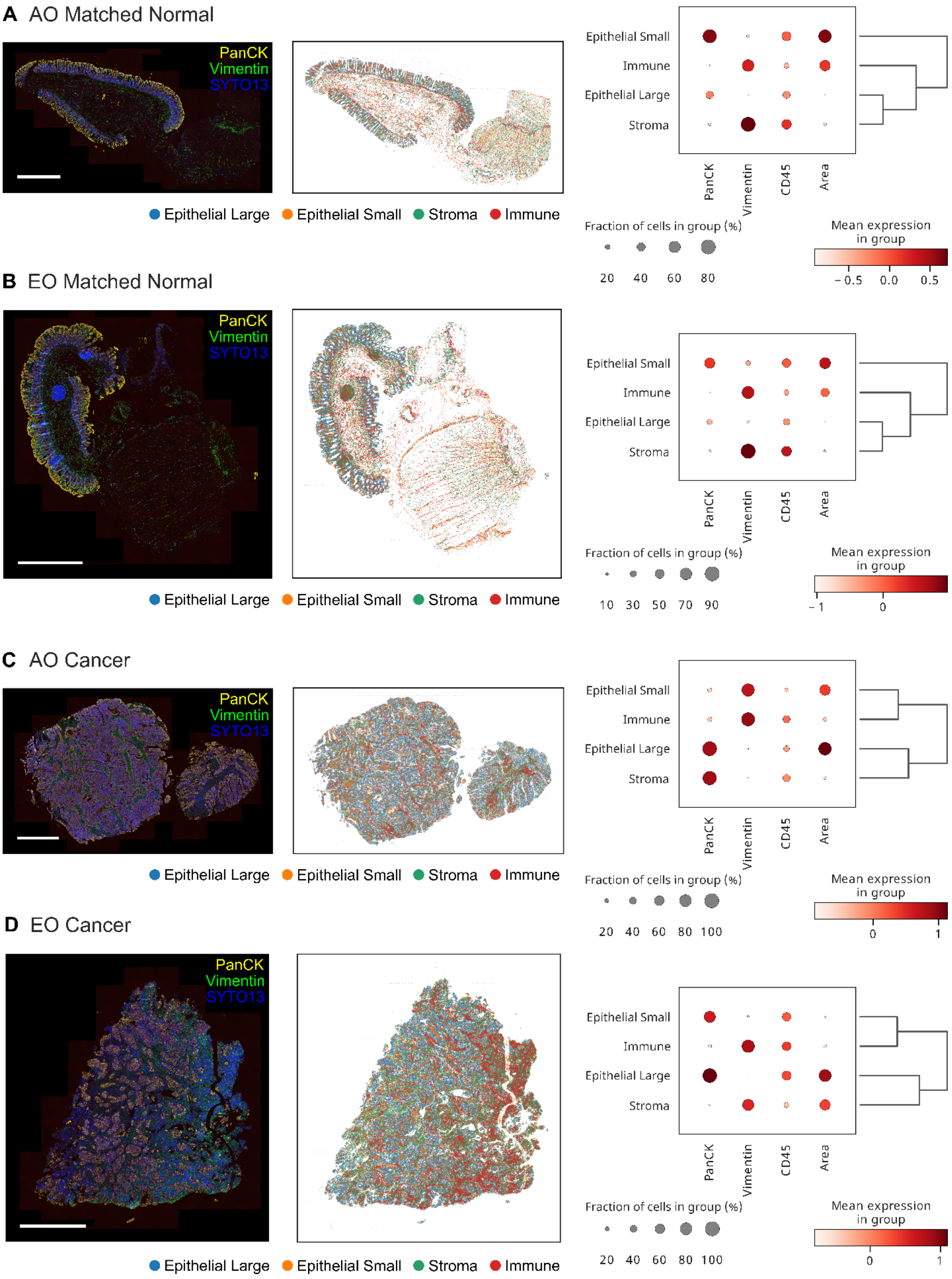
A AO Matched Normal
B EO Matched Normal
C AO Cancer
D EO Cancer
PanCK
Vimentin
SYTO13
Epithelial Large
Epithelial Small
Stroma
Immune
Epithelial Small
Immune
Epithelial Large
Stroma
PanCK
Vimentin
CD45
Area
Fraction of cells in group (%)
20 40 60 80
Mean expression in group
−0.5 0.0 0.5
10 30 50 70 90
−1 0
20 40 60 80 100
0 1

**Figure S9. A cell clustering analysis reveals distinct cellular phenotypes in CRC.** Representative tissue images show the raw immunofluorescence (left), the spatial distribution of cell phenotypes identified via unsupervised Louvain clustering (middle) and a dot plot quantifying the expression of individual features (right) for (**A**) AO Matched Normal, (**B**) EO Matched Normal, (**C**) AO Cancer, and (**D**) EO Cancer. The four cellular phenotypes are the following: epithelial large (PanCK-high, Vimentin-low, cell area-high), epithelial small (PanCK-high, Vimentin-low, cell area-low), stroma (PanCK-low, Vimentin-high, cell area-high), and immune (PanCK-low, Vimentin-high, cell area-low). Scale bars, 1 mm.

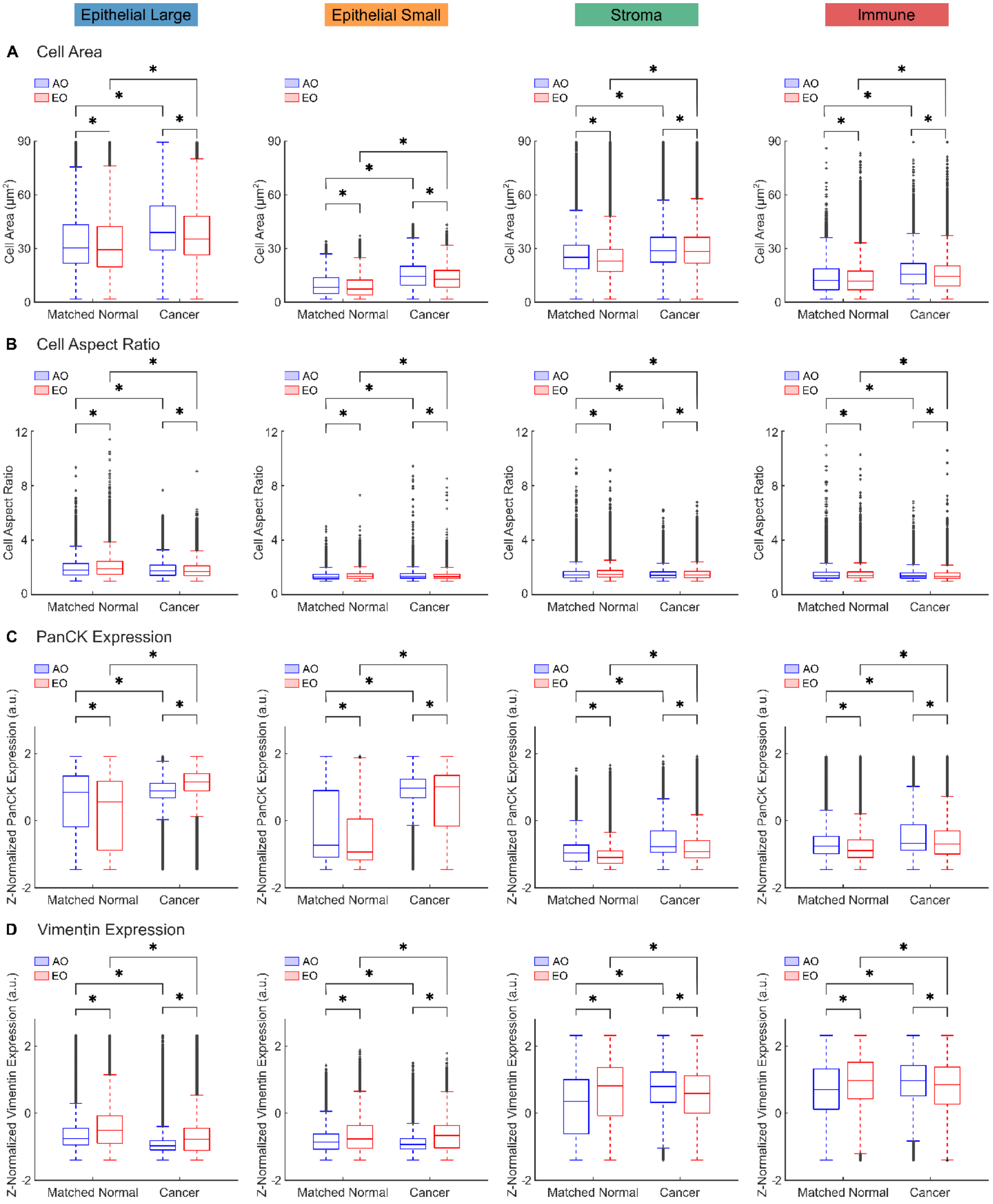


**Figure S10. Quantification of single cell features for individual phenotypes in AO and EO CRC.** Box plots for (**D**) cell area, (**E**) cell aspect ratio, (**F**) PanCK expression, and (**G**) Vimentin separated for the four cell phenotypes: epithelial large (left, n = 32,936 for AO Matched Normal, n = 262,137 for AO Cancer, n = 61,101 for EO Matched Normal, n = 206,389 for EO Cancer), epithelial small (middle-left, n = 45,357 for AO Matched Normal, n = 186,585 for AO Cancer, n = 94,247 for EO Matched Normal, n = 176,501 for EO Cancer), stroma (middle-right, n = 47,526 for AO Matched Normal, n = 134,497 for AO Cancer, n = 101,933 for EO Matched Normal, n = 203,185 for EO Cancer), immune (right, n = 50,862 for AO Matched Normal, n = 170,774 for AO Cancer, n = 122,345 for EO Matched Normal, n = 258,671 for EO Cancer). The expressions of epithelial and stromal markers were z-normalized to facilitate comparisons across a large dataset. Statistical significance was assessed using a Scheirer-Ray-Hare test with Dunn's post-hoc test and Bonferroni correction. * indicates significant differences ($P < 0.05$) between cancer and matched normal tissues within onset groups (AO, EO) and between onset groups within tissue types (matched normal, cancer).

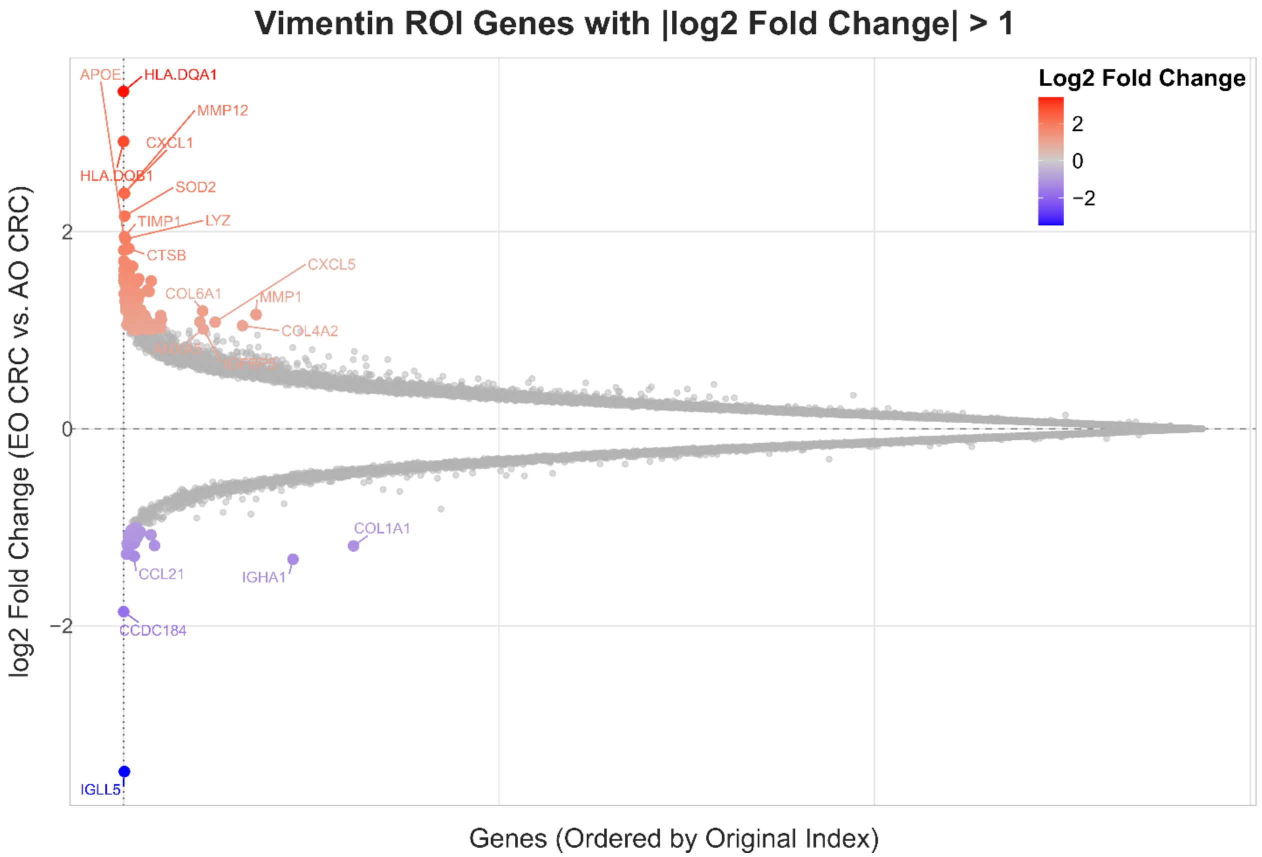


**Figure S11. Differentially expressed genes in stromal cells of EO CRC with respect to AO CRC.** Distribution of gene expression fold changes in vimentin-positive stromal cells in EO vs. AO CRC samples. The x-axis represents genes ordered by their original dataset index, and the y-axis shows log2 fold change values. Small grey points represent all genes analyzed, while colored points highlight genes exceeding the fold change threshold (|log2 Fold Change| > 1). Points are colored according to their fold change value (red = upregulated in EO , blue = downregulated in EO). Among the upregulated genes in the EO stroma are several ECM-related genes including collagens (COL6A1), ECM remodeling enzymes (LOXL1, TIMP1, MMP12), and chemokines (CXCL1), consistent with a pro-fibrotic phenotype.

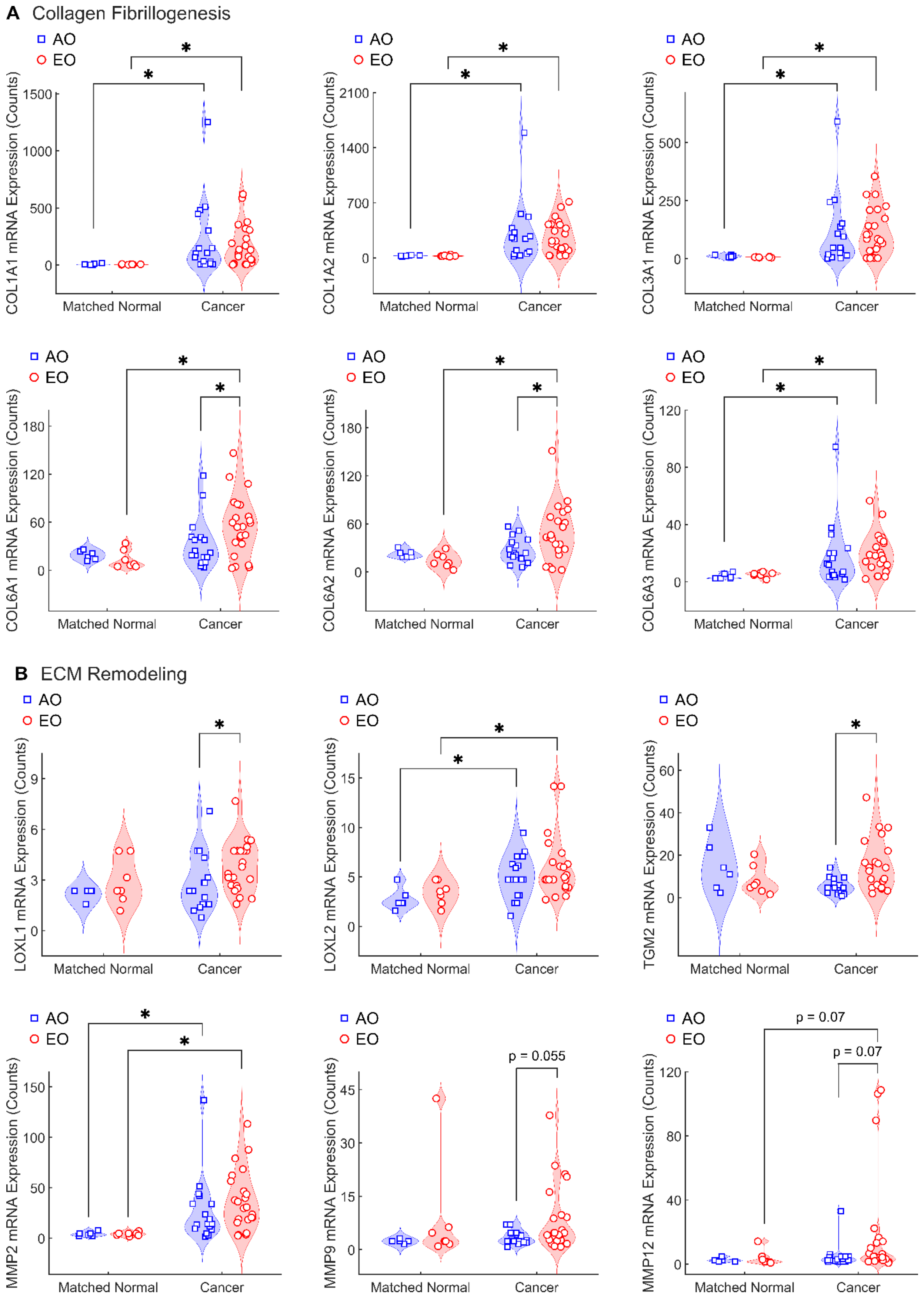
A Collagen Fibrillogenesis
AO
EO
*
COL1A1 mRNA Expression (Counts)
COL1A2 mRNA Expression (Counts)
COL3A1 mRNA Expression (Counts)
COL6A1 mRNA Expression (Counts)
COL6A2 mRNA Expression (Counts)
COL6A3 mRNA Expression (Counts)
Matched Normal
Cancer
B ECM Remodeling
LOXL1 mRNA Expression (Counts)
LOXL2 mRNA Expression (Counts)
TGM2 mRNA Expression (Counts)
MMP2 mRNA Expression (Counts)
MMP9 mRNA Expression (Counts)
MMP12 mRNA Expression (Counts)
p = 0.055
p = 0.07
p = 0.07

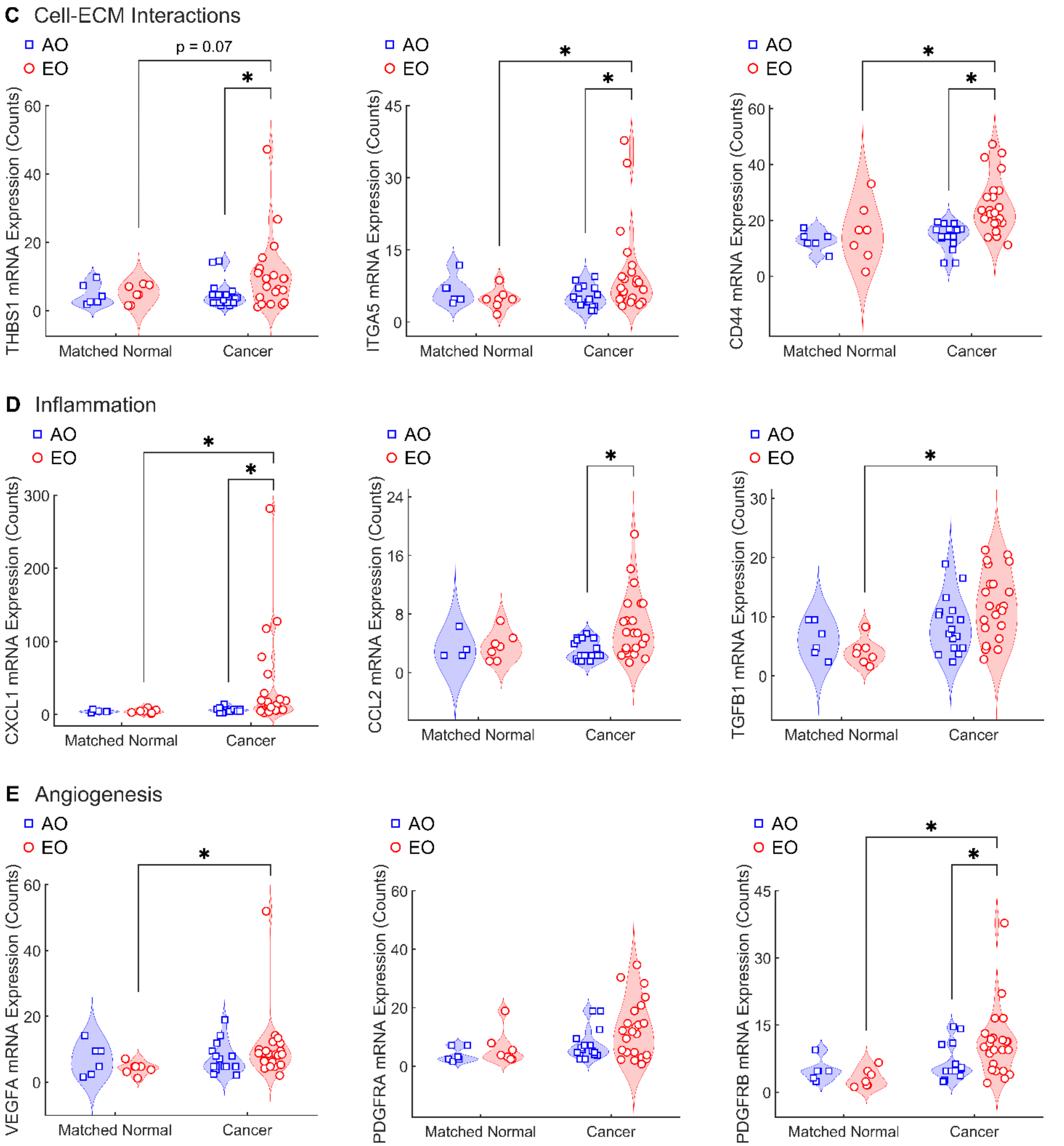


**Figure S12. Gene expression patterns in stromal cells from AO CRC and EO CRC patients.** Expression levels of selected genes associated with (**A**) collagen fibrillogenesis, (**B**) ECM remodeling, (**C**) cell-ECM interactions, (**D**) inflammation, and (**E**) angiogenesis. Data are presented as violin plots with data points indicating individual measurements. Statistical significance was assessed using a Scheirer-Ray-Hare test with Dunn's post-hoc test and Bonferroni correction. * indicates significant differences ($P < 0.05$) between cancer and matched normal tissues within onset groups (AO, EO) and between onset groups within tissue types (matched normal, cancer).

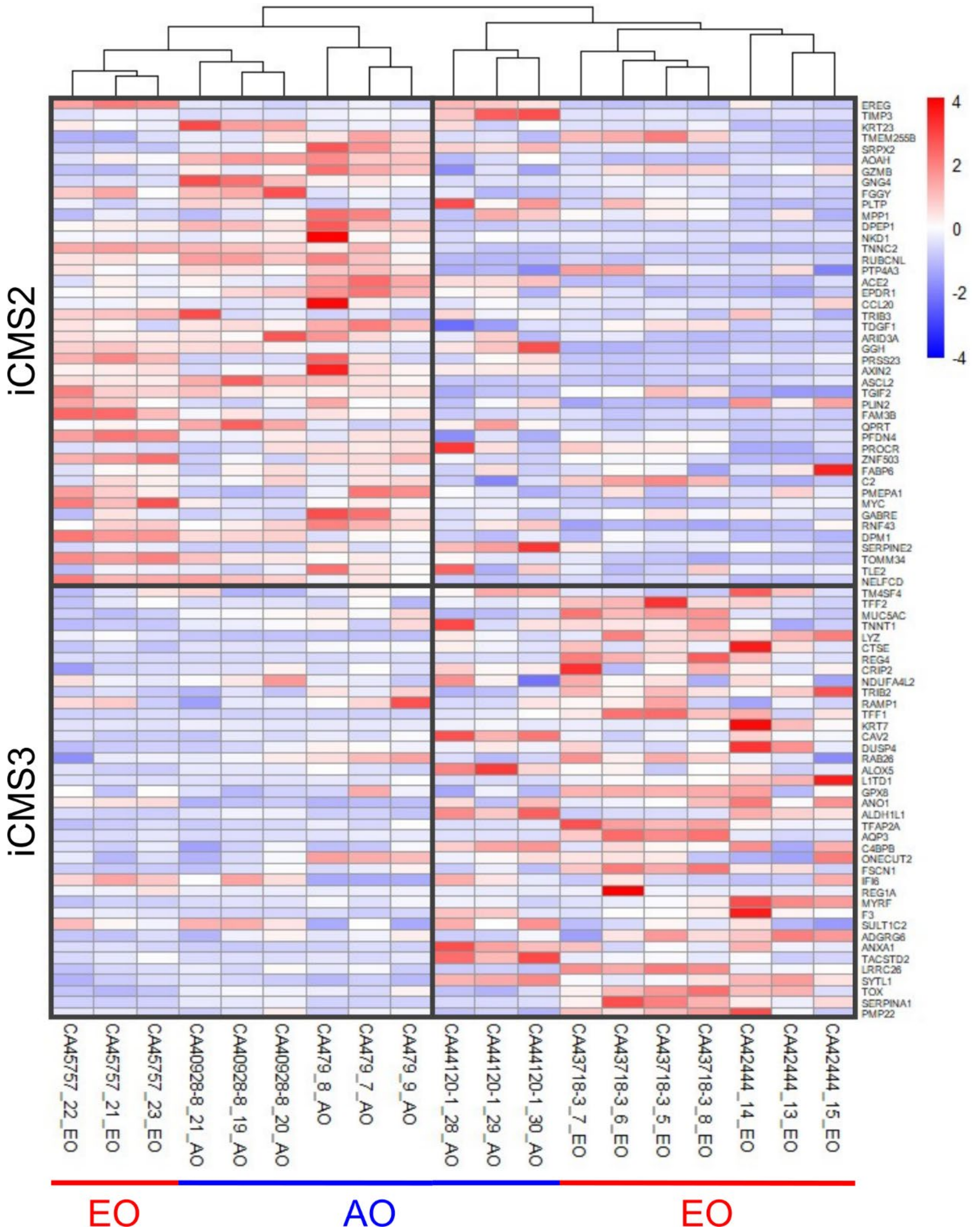


**Figure S13. Classification of colorectal cancer samples into iCMS subtypes based on epithelial cell gene expression.** Heatmap showing unsupervised hierarchical clustering of colorectal cancer samples based on intrinsic epithelial gene expression patterns using the top 100 iCMS marker genes from Joanito et al.[69] The analysis reveals two major clusters corresponding to iCMS2 (upper cluster) and iCMS3 (lower cluster) subtypes. Samples are represented in columns, with their IDs shown at the bottom along with age of onset (AO vs. EO). Marker genes are represented in rows. The color scale indicates expression levels, with red representing high expression and blue representing low expression. The clustering demonstrates distinct gene expression patterns between the two subtypes, with iCMS2 samples showing elevated expression of WNT/β-catenin and MYC pathway genes (upper section), while iCMS3 samples show upregulation of MAPK pathway genes and inflammatory response signatures (lower section).

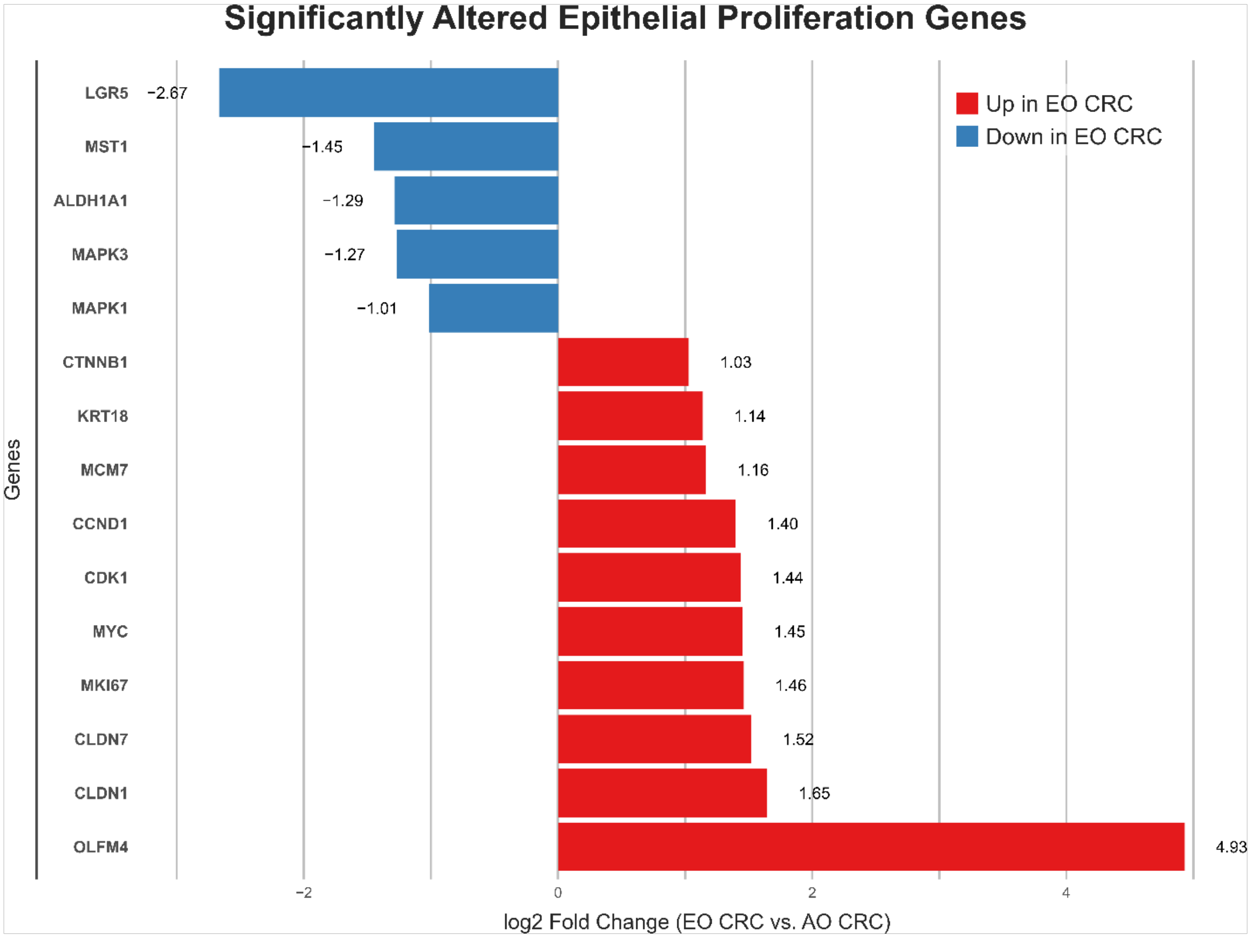


**Figure S14. Proliferation genes in epithelial cells of EO CRC with respect to AO CRC.** (A) Altered genes related to epithelial cell proliferation in EO CRC determined from PanCK-positive ROIs using a *limma*-based differential expression analysis with patient ID as a covariate. Significantly altered genes were filtered based on statistical significance (adjusted $P$ value < 0.05) and fold change magnitude (|log2 Fold Change| > 1). The horizontal bars represent log2 fold changes, with red bars indicating upregulation in EO CRC and blue bars indicating downregulation. Positive values (red) represent higher expression in EO CRC epithelial cells, while negative values (blue) show reduced expression compared to AO CRC. Numerical values indicate the precise log2 fold change for each gene.

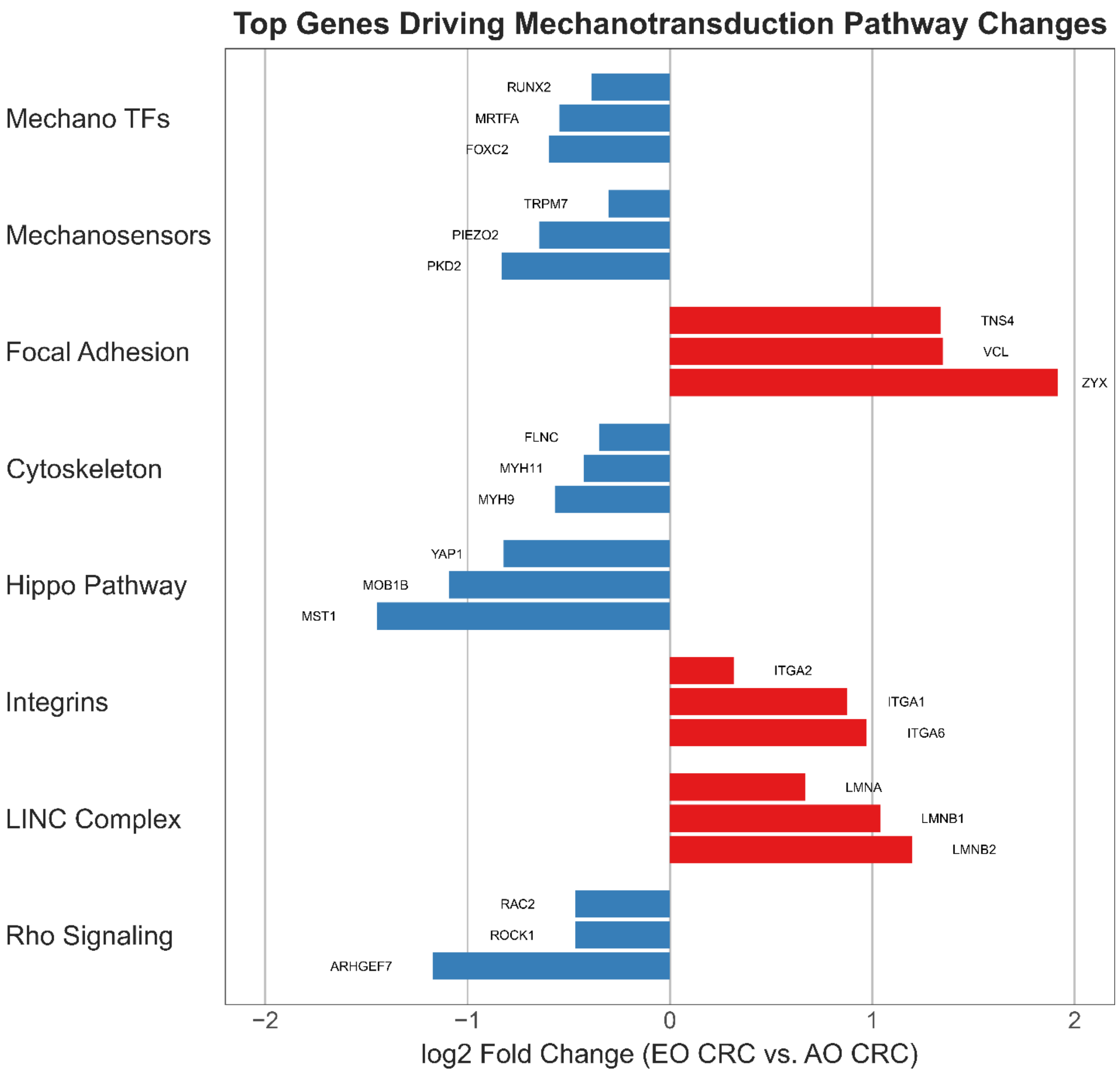


**Figure S15. Top genes driving mechanotransduction pathway changes in epithelial cells of EO CRC compared to AO CRC.** (**A**) Top three driver genes with the highest absolute log fold change values from each mechanotransduction pathway, consistent with the pathway's overall direction (red: upregulated in EO CRC, blue: downregulated in AO CRC). Using CAMERA gene set analysis,[110] we analyzed eight major mechanotransduction categories: Mechanosensitive Transcription Factors, Mechanosensors, Focal Adhesion, Cytoskeleton, Hippo Pathway, Integrins, LINC Complex, and Rho GTPase Signaling.

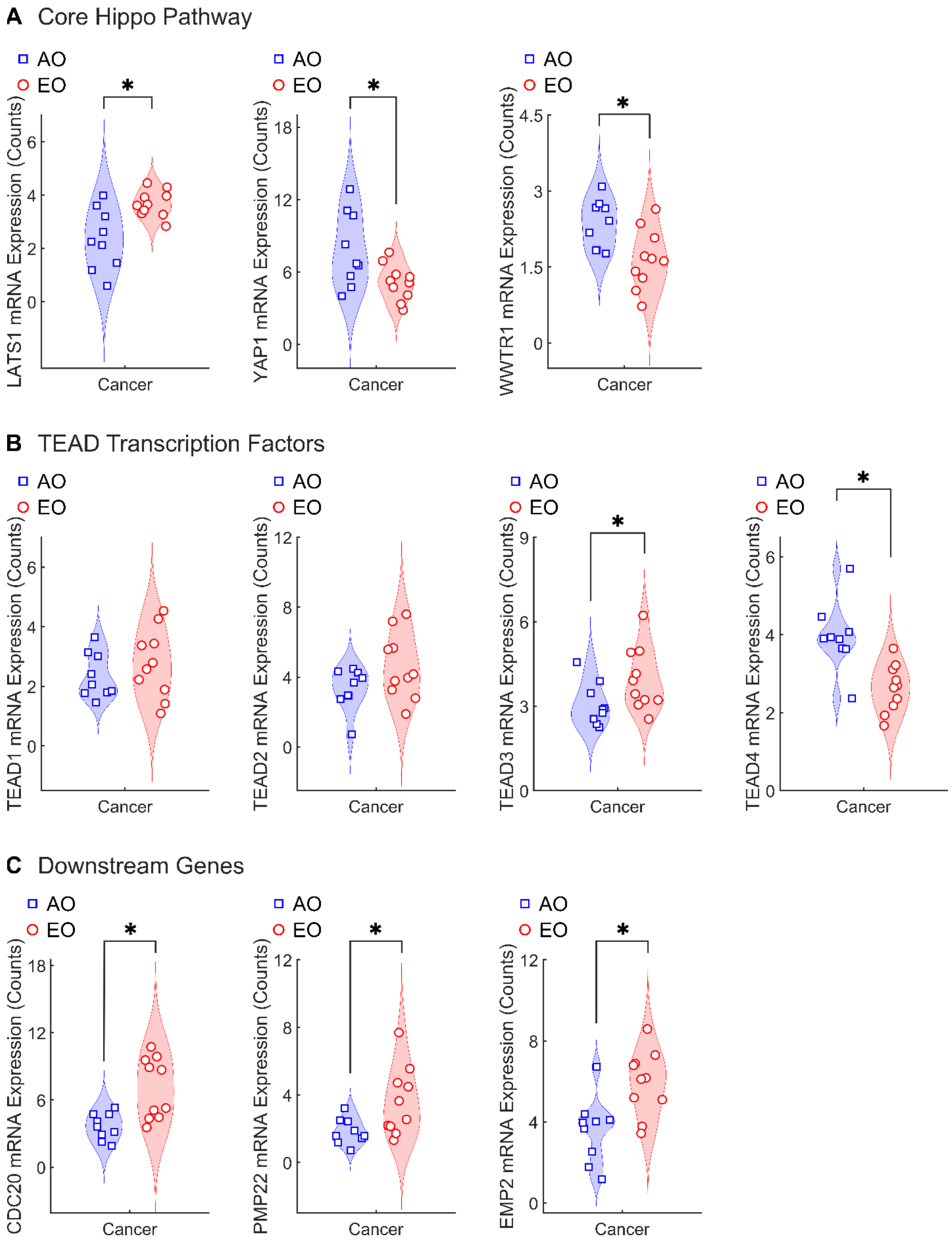


**Figure S16. Expression of Hippo-YAP genes in epithelial cells of EO CRC versus AO CRC.** (**A**) mRNA expression differences of key Hippo-YAP pathway components between EO CRC and AO CRC in PanCK-positive epithelial regions. The analysis is organized into three functional categories: (**A**) Core Hippo Pathway components (LATS1, YAP1, WWTR1), (**B**) TEAD transcription factors (TEAD1-4), and (**C**) Downstream target genes (CDC20, PMP22, EMP2). Violin plots with overlaid data points show the distribution of normalized mRNA counts from GeoMx spatial transcriptomics analysis of PanCK+ ROIs. Statistical significance was assessed using Dunn's test with Bonferroni correction. * indicates significant differences ($P$ < 0.05) between AO and EO onset groups within cancer tissues.

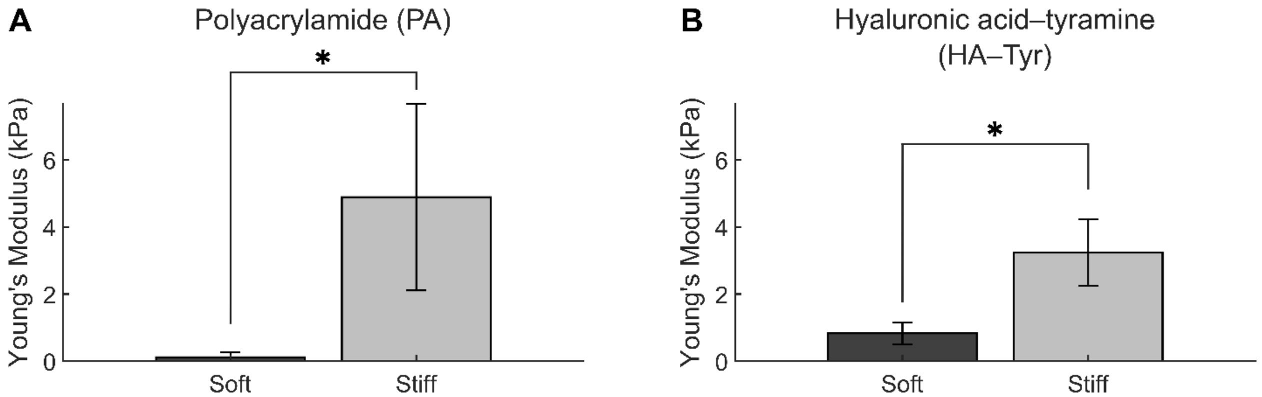


**Figure S17. Hydrogel stiffness used to reproduce colorectal tissue mechanics in vitro.** We reproduced the mechanical properties of primary colorectal tissues (Figure 1) by employing (**A**) polyacrylamide (PA) hydrogels for 2D in vitro cultures and (**B**) hyaluronic acid-tyramine (HA-Tyr) hydrogels for 3D in vitro cultures. We used unconfined compression to quantify the Young's modulus of soft ($E = 0.12 \pm 0.15$ kPa, n = 29) and stiff ($E = 4.89 \pm 2.78$ kPa, n = 28) PA gels, as well as that of soft (E = 0.8 ± 0.1 kPa, n = 7) and stiff (E = 3.2 ± 0.4 kPa, n = 7) HA-Tyr gels. Data are presented as mean ± SD. Statistical significance was assessed using Dunn's test with Bonferroni correction. * indicates statistically significant differences ($P < 0.05$) in pairwise stiffness comparisons within each substrate type.

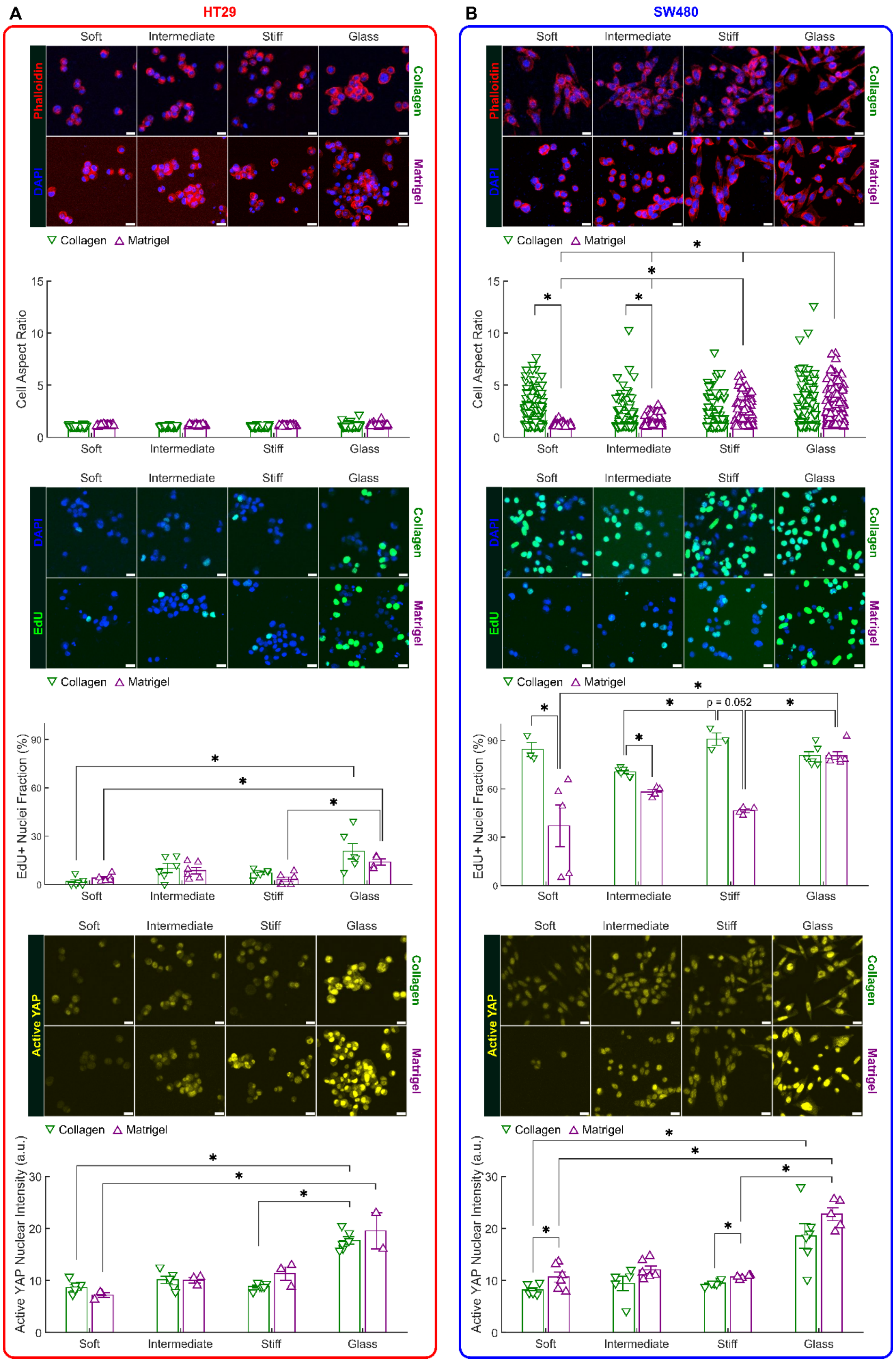
A
HT29
B
SW480
Soft
Intermediate
Stiff
Glass
Phalloidin
DAPI
EdU
Active YAP
Collagen
Matrigel
Cell Aspect Ratio
EdU+ Nuclei Fraction (%)
Active YAP Nuclear Intensity (a.u.)
p = 0.052

**Figure S18. Effect of substrate stiffness and adhesive ligand on CRC cell lines.** HT29 and SW480 cells were cultured on Collagen and Matrigel-coated polyacrylamide (PA) substrates of graded stiffness. PA gels were tuned to generate substrates with low (Soft: E = 43.9 ± 15.01 Pa, n = 3), medium (Intermediate: E = 282.1 ± 80.82 Pa, n = 4), and high (Stiff: E = 2672.7 ± 796.07 Pa, n = 4) stiffness based on direct measurements from primary tissues (cf. Figure 1F). We carried out immunofluorescence staining after 72 hours in culture using phalloidin (red), DAPI (blue), EdU (green), and active YAP (yellow). Scale bars, 20 µm. (**A**) Representative images for HT29 cells and associated quantifications of cellular aspect ratio (Collagen: n = 129 for Soft, n = 136 for Intermediate, n = 141 for Stiff, n = 128 for Glass; Matrigel: n = 48 for Soft, n = 51 for Intermediate, n = 59 for Stiff, n = 71 for Glass), EdU incorporation (Collagen: n = 6 for Soft, n = 6 for Intermediate, n = 6 for Stiff, n = 6 for Glass; Matrigel: n = 6 for Soft, n = 6 for Intermediate, n = 6 for Stiff, n = 4 for Glass), and YAP activity (Collagen: n = 6 for Soft, n = 6 for Intermediate, n = 6 for Stiff, n = 6 for Glass; Matrigel: n = 3 for Soft, n = 3 for Intermediate, n = 3 for Stiff, n = 2 for Glass). (**B**) Representative images for SW480 and associated quantifications of cellular aspect ratio (Collagen: n = 129 for Soft, n = 107 for Intermediate, n = 128 for Stiff, n = 108 for Glass; Matrigel: n = 118 for Soft, n = 179 for Intermediate, n = 143 for Stiff, n = 173 for Glass), EdU incorporation (Collagen: n = 3 for Soft, n = 6 for Intermediate, n = 3 for Stiff, n = 6 for Glass; Matrigel: n = 6 for Soft, n = 5 for Intermediate, n = 6 for Stiff, n = 6 for Glass), and YAP activity (Collagen: n = 6 for Soft, n = 5 for Intermediate, n = 6 for Stiff, n = 6 for Glass; Matrigel: n = 6 for Soft, n = 6 for Intermediate, n = 6 for Stiff, n = 5 for Glass). Data are presented as mean ± SEM with data points indicating individual measurements. Statistical significance was assessed using Scheirer-Ray-Hare test followed by Dunn's post-hoc test with Bonferroni correction. * indicates statistically significant differences ($P < 0.05$) between groups.

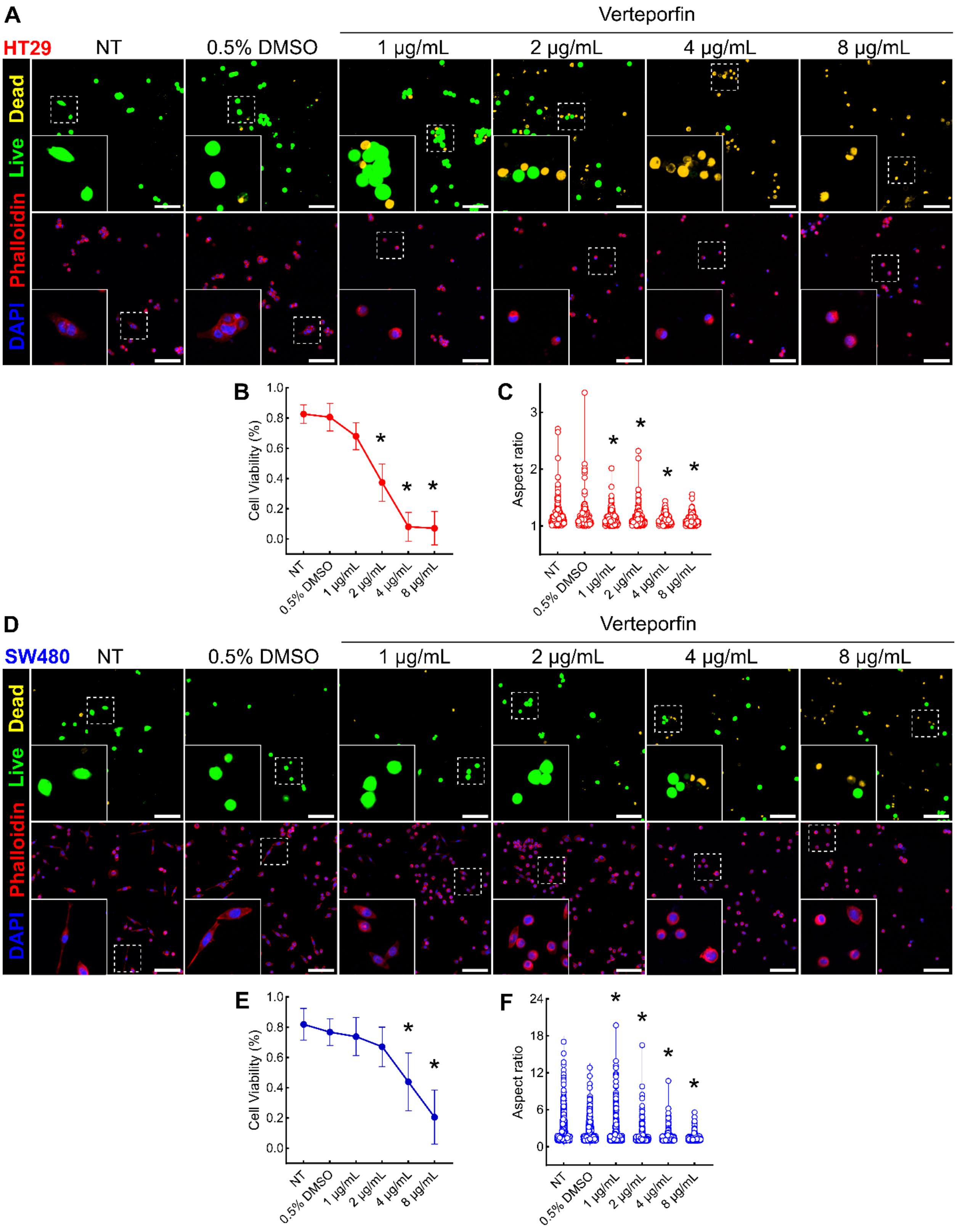
A
Verteporfin
HT29
NT
0.5% DMSO
1 μg/mL
2 μg/mL
4 μg/mL
8 μg/mL
Dead
Live
Phalloidin
DAPI
B
Cell Viability (%)
1.0
0.8
0.6
0.4
0.2
0.0
NT
0.5% DMSO
1 μg/mL
2 μg/mL
4 μg/mL
8 μg/mL
C
Aspect ratio
3
2
1
D
Verteporfin
SW480
E
F
24
18
12
6
0
*

**Figure S19. Dose-dependent effect of YAP inhibition with verteporfin on CRC cell lines.** Fluorescence staining of HT29 (**A**) and SW480 (**D**) cells cultured on Matrigel-coated glass for 72 hours. Cells received no treatment (NT), media with vehicle control (0.5% DMSO), or media with varying concentrations of verteporfin (1–8 μg $mL^{-1}$) during the final 48 hours of culture. Cells were stained for calcein AM (green, live), ethidium homodimer-1 (yellow, dead), DAPI (blue), and phalloidin (red). Scale bars, 100 μm. Cell viability quantification for HT29 (**B**) and SW480 (**E**) cells (n = 12 per group) using 2 technical replicates and 6 ROIs per replicate. Aspect ratio quantification for HT29 (**C**) and SW480 (**F**) cells. Sample sizes: HT29 (n = 277 for NT, n = 254 for DMSO, n = 266 for 1 μg/mL, n = 255 for 2 μg/mL, n = 232 for 4 μg/mL, n = 247 for 8 μg/mL); SW480 (n = 655 for NT, n = 679 for DMSO, n = 526 for 1 μg/mL, n = 586 for 2 μg/mL, n = 486 for 4 μg/mL, n = 337 for 8 μg/mL). Cell viability data are presented as mean ± SD, while cell aspect ratio data are presented as violin plots with data points indicating individual measurements. Statistical significance was assessed using Kruskal-Wallis test followed by Dunn's post-hoc test with Bonferroni correction. * indicates significant differences ($P < 0.05$) in pairwise comparisons with the NT group.

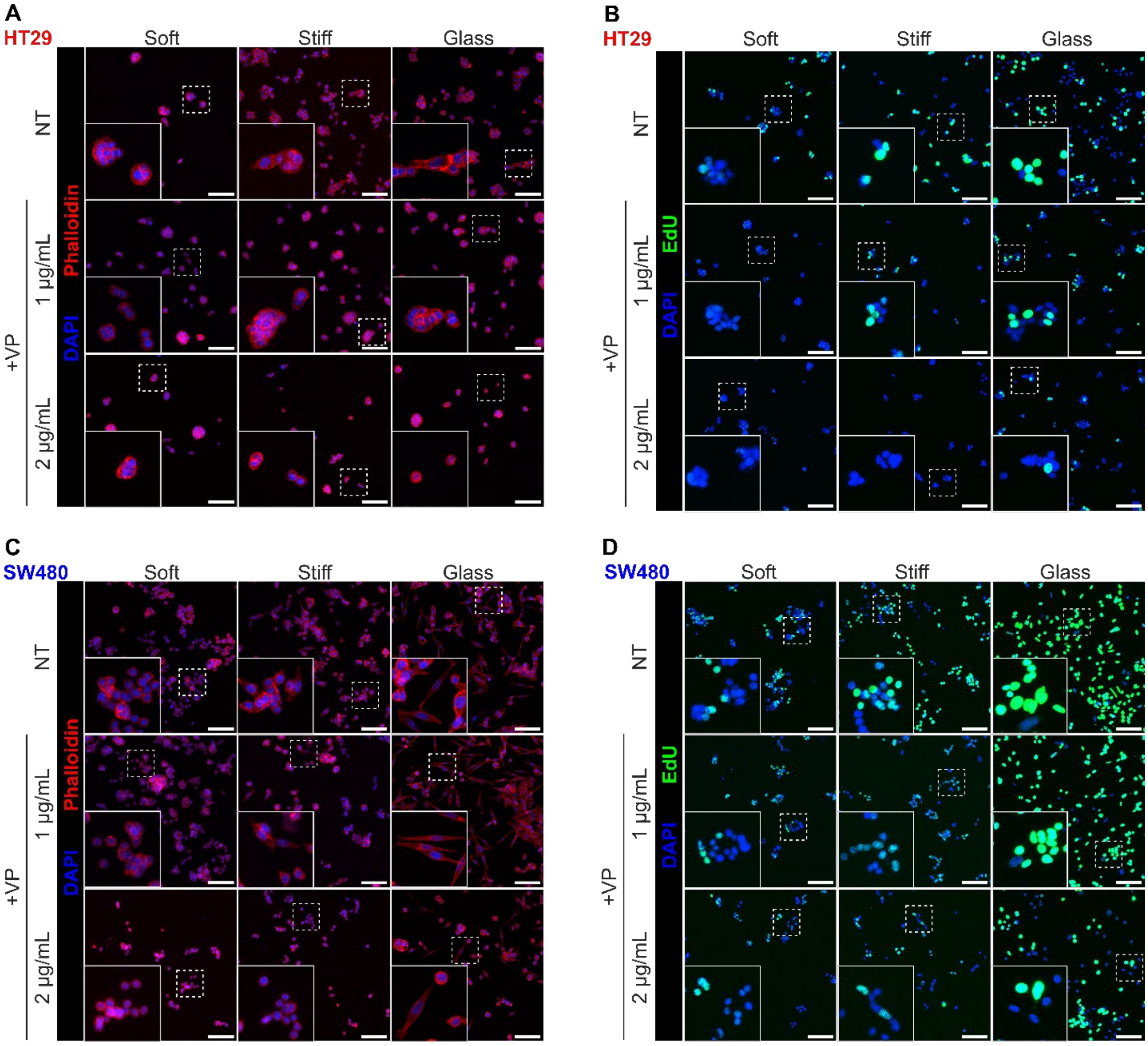


**Figure S20. Stiffness-dependent changes in cell morphology and proliferation of CRC cell lines upon YAP inhibition.** (**A, C**) Immunofluorescence staining for phalloidin (red) and DAPI (blue) in HT29 and SW480 cells. (**B, D**) EdU (green) and DAPI (blue) staining in HT29 and SW480 cells. Cells were cultured on graded substrate stiffness for 72 hours, with verteporfin treatment (1 or 2 µg mL$^{-1}$) applied during the final 48 hours. Scale bars, 100 µm. Quantifications are presented in Figure 6.

| AO CRC | | | | | | | |
|---|---|---|---|---|---|---|---|
| **Patient ID** | **Age (years)** | **BMI** | **Sex** | **Ethnicity** | **Race** | **Cancer Location** | **Stage** |
| AO_01 | 70 | 33.2 | M | Non-Hispanic | Black | Cecum | 2 |
| AO_02* | 75 | 32.3 | F | Hispanic | White | Sigmoid | 1 |
| AO_03 | 50 | 50.0 | F | N/A | N/A | Sigmoid | 3 |
| AO_04 | 55 | 36.8 | M | Hispanic | White | Right colon | 3 |
| AO_05* | 68 | 34.0 | M | Non-Hispanic | Black | Hepatic flexure | 3 |
| AO_06 | 71 | 25.0 | F | Non-Hispanic | Asian | Right colon | 1 |
| AO_07*† | 63 | 31.1 | M | Non-Hispanic | White | Sigmoid | 2 |
| AO_08† | 58 | 39.6 | M | Hispanic | White | Cecum | 1 |
| AO_09 | 60 | 26.5 | M | Non-Hispanic | White | Right colon | 4 |
| AO_10 | 69 | 27.4 | M | Hispanic | White | Right colon | 4 |
| AO_11 | 72 | 33.5 | M | Non-Hispanic | White | Cecum | 2 |
| AO_12† | 50 | 37.9 | M | Hispanic | White | Rectosigmoid | 3 |
| AO_13 | 78 | 22.6 | F | N/A | N/A | Cecum | 3 |
| AO_14 | 72 | 32.0 | M | Non-Hispanic | White | Cecum | 2 |
| AO_15 | 85 | 22.7 | F | Non-Hispanic | White | Sigmoid | 2 |
| AO_16 | 60 | 44.9 | F | Non-Hispanic | Black | Sigmoid | 2 |
| AO_17 | 85 | 24.1 | M | Non-Hispanic | Black | Cecum | 1 |
| AO_18 | 82 | 21.4 | F | Non-Hispanic | White | Splenic flexure | 3 |
| AO_19 | 66 | 43.9 | F | Non-Hispanic | White | Cecum | 3 |
| Mean | 68 | 32.6 | | | | | |
| STD | 11 | 8.2 | | | | | |

**Table S1. AO CRC patient demographic and tumor information.** Description of demographic and clinical characteristics of AO CRC patients involved in the study. Samples from patient IDs labeled with * were used to quantify local RNA expression using the GeoMx® RNA assay. Samples from patient IDs labeled with † were used to generate organoids for in vitro culture. N/A, not available.

| EO CRC | | | | | | | |
|---|---|---|---|---|---|---|---|
| Patient ID | Age (years) | BMI | Sex | Ethnicity | Race | Cancer Location | Stage |
| EO_01 | 45 | 30.0 | F | Non-Hispanic | Black | Rectum | 3 |
| EO_02* | 45 | 21.0 | M | Hispanic | White | Sigmoid | 2 |
| EO_03† | 41 | 27.0 | M | Non-Hispanic | Asian | Sigmoid | 3 |
| EO_04 | 46 | 28.2 | M | Hispanic | White | Rectosigmoid | 1 |
| EO_05 | 42 | 31.0 | F | Hispanic | White | Cecum | 2 |
| EO_06* | 40 | 24.3 | M | Non-Hispanic | White | Rectosigmoid | 2 |
| EO_07 | 45 | 29.0 | M | Non-Hispanic | Asian | Rectum | 2 |
| EO_08* | 42 | 23.0 | F | Non-Hispanic | Black | Sigmoid | 2 |
| EO_09† | 43 | 31.7 | M | Non-Hispanic | White | Sigmoid | 3 |
| EO_10† | 47 | 25.8 | M | Non-Hispanic | White | Rectosigmoid | 1 |
| EO_11 | 41 | 30.1 | F | Non-Hispanic | Black | Sigmoid | 3 |
| EO_12 | 41 | 21.5 | F | Non-Hispanic | Black | Cecum | 2 |
| EO_13 | 46 | 29.5 | M | Non-Hispanic | Asian | Rectosigmoid | 4 |
| EO_14 | 46 | 28.3 | M | Non-Hispanic | White | Sigmoid | 2 |
| Mean | 44 | 27.2 | | | | | |
| STD | 2 | 3.5 | | | | | |

**Table S2. EO CRC patient demographic and tumor information.** Description of demographic and clinical characteristics of EO CRC patients involved in the study. Samples from patient IDs labeled with * were used to quantify local RNA expression using the GeoMx® RNA assay. Samples from patient IDs labeled with † were used to generate organoids for in vitro culture.

| Patient ID | AMEL | CSF 1PO | D5S 818 | D7S 820 | D13S 317 | D16S 539 | D21S 11 | TH01 | TPOX | vWA | D1S 1656 | D2S 441 |
|---|---|---|---|---|---|---|---|---|---|---|---|---|
| AO_07 | X,Y | 11,12 | 12,13 | 8,12 | 11 | 8,11 | 27,30 | 7,9 | 8,11 | 17 | 16.3,17.3 | 14 |
| AO_08 | X,Y | 11,13 | 9 | 7,12 | 9,12 | 11 | 30,32.2 | 6,9.3 | 8,11 | 16,17 | 16.3,17.3 | 10,14 |
| AO_12 | X,Y | 11,12 | 12 | 11,14 | 11,12 | 9,12 | 31.2,32.2 | 7,9.3 | 8,9 | 16,17 | 16 | 10 |
| EO_03 | X,Y | 9 | 11 | 11 | 10,11 | 9,11 | 29,30 | 6 | 10,11 | 16 | 17 | 12 |
| EO_09 | X,Y | 12,13 | 11,12 | 7,8 | 11 | 12 | 28,31.2 | 6 | 8,10 | 17 | 11,15.3 | 11,14 |
| EO_10 | X,Y | 12,13 | 10,13 | 9,12 | 11,12 | 11 | 30.2,32.2 | 9.3 | 8 | 17,19 | 14,15 | 11,13 |

| Patient ID | D2S 1338 | D3S 1358 | D8S 1179 | D10S1248 | D12S391 | D18S51 | D19S433 | D22S1045 | DYS391 | FGA | PentaD | PentaE |
|---|---|---|---|---|---|---|---|---|---|---|---|---|
| AO_07 | 25 | 15,18 | 12,14 | 14 | 22,24 | 10,15 | 15.2,16 | 15,17 | 10 | 21,22 | 9,13 | 16,19 |
| AO_08 | 20,23 | 15,17 | 10,11 | 15,18 | 20 | 14,17 | 14,15 | 16 | 10 | 23,27 | 9,10 | 16,20 |
| AO_12 | 19,23 | 15,17 | 13,14 | 13,14 | 18 | 14,17 | 13,14 | 16,17 | 10 | 19,24 | 13 | 12 |
| EO_03 | 20,26 | 15 | 10 | 14,15 | 18,19 | 15,16 | 15.2,16 | 11,17 | 10 | 21,23 | 10,13 | 19 |
| EO_09 | 22,23 | 14,15 | 12,13 | 14,16 | 21,22 | 12,17 | 15,16.2 | 11 | 9 | 21,22 | 13,15 | 7,12 |
| EO_10 | 24,27 | 14,17 | 10,11 | 13,15 | 22,23 | 14,15 | 15 | 15,16 | 10 | 24,25 | 12 | 5,12 |

**Table S3. Short-tandem repeat fingerprinting.** Patient-derived epithelial organoids were compared to primary formalin-fixed tissues. When recovered from cryopreservation, samples demonstrated identical short-tandem repeat fingerprinting compared to matched primary samples, and unique fingerprints compared to one another.
